# Development of a chemical probe to enable characterization of the casein kinase 1γ subfamily

**DOI:** 10.1101/2025.09.10.675406

**Authors:** Jacob L. Capener, Thomas W. Kramer, Frances M. Bashore, Emily Flory, Fengling Li, Blair L. Strang, Alison D. Axtman

## Abstract

The casein kinase 1γ (CK1γ) subfamily, while severely understudied, is implicated in diverse disease-relevant pathways, including WNT signaling and human cytomegalovirus (HCMV) replication. While genetic tools exist to study CK1γ, the selective inhibition of CK1γ through pharmacological means remains underexplored. Chemical probes, or potent and selective inhibitors, remain one of the most powerful pharmacological tools for uncovering protein biology. Herein, we developed several novel assays for assessing target engagement with the CK1γ subfamily in cells. Enabled by these assays, we conducted a comprehensive structure–activity relationship (SAR) campaign to develop the first chemical probe, SGC-CK1γ-1, for the CK1γ subfamily. SGC-CK1γ-1, which was developed alongside a structurally related negative control compound, potently and selectively inhibited the CK1γ kinases in living cells, plus inhibited both WNT signaling and human cytomegalovirus replication.

## Introduction

The casein kinase 1 (CK1) family is composed of several kinases involved in fundamental biological processes. Among these kinases, the CK1γ subfamily, consisting of CK1γ1, CK1γ2, and CK1γ3, represents a poorly investigated group of “dark” kinases as defined by the National Institute of Health initiative, Illuminating the Druggable Genome^1,2^. The CK1γ subfamily members are highly conserved in their kinase domains, with 92% sequence homology^1^. Unlike the broader CK1 family, the CK1γ subfamily is anchored to the membrane via a conserved C-terminal palmitoylation site^3^. Although they have unique functions, very little is known about the CK1γ kinases when compared to the other CK1 family members. CK1γ kinases are mentioned in fewer research articles and patents than other CK1 family members (Figure 1A, B)^4^. Furthermore, many of the citations and patents that mention CK1γ kinases only refer to them in association with the actual protein of interest, often another CK1 isoform. Interestingly, the CK1γ kinases demonstrate some similar and some non-overlapping cellular functions when compared to one another and are distinct from the larger CK1 family^1,5^. Emerging literature supports that the cellular processes regulated in part by the CK1γ subfamily positions them as potential therapeutic targets^1,6,7,8,9,10,11,12,13,14^.

**Figure 1.**
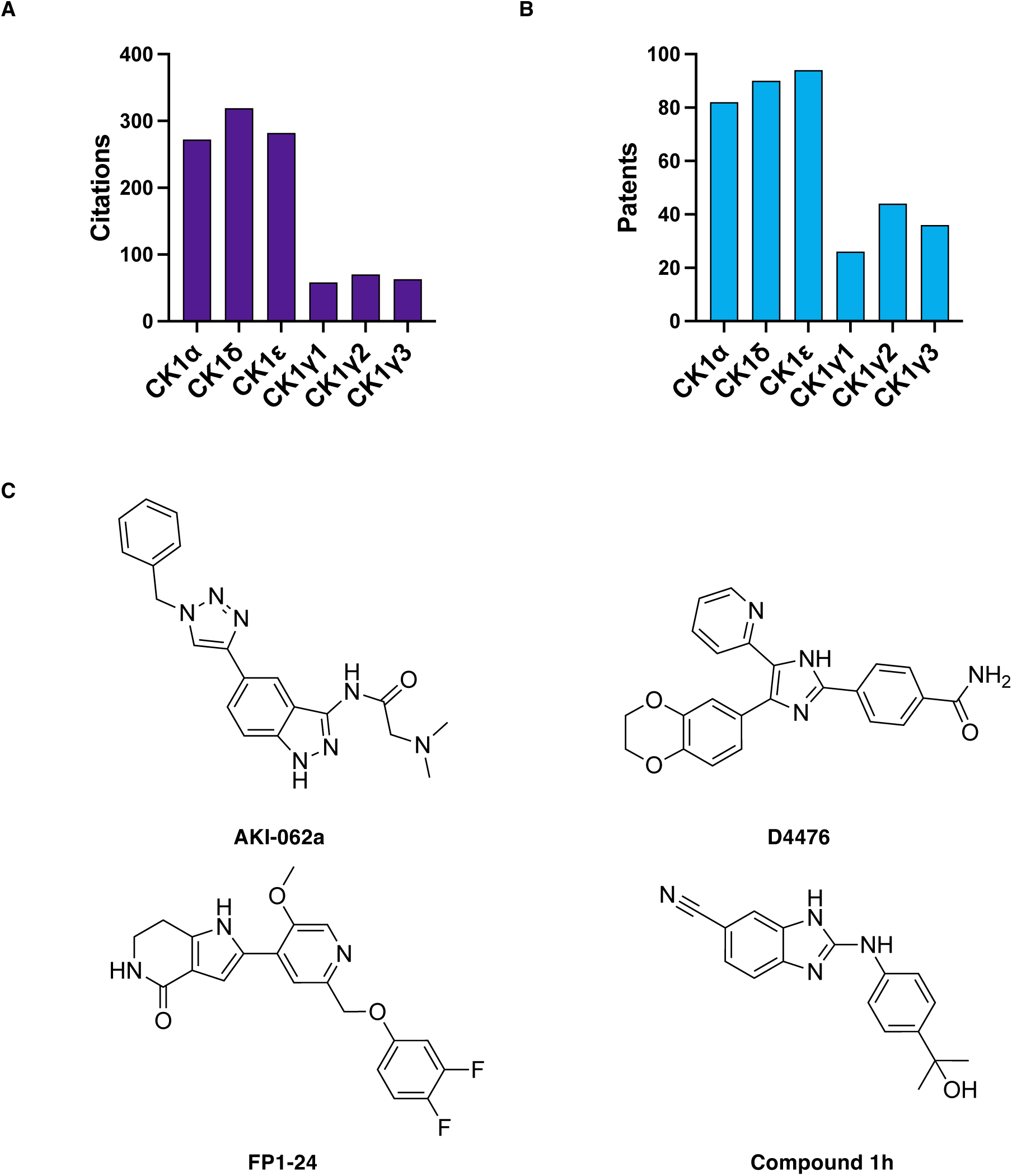
A summary of current literature references and inhibitors for the CK1γ isoforms. a) Count of citations that mention each CK1 isoform found through a Scifinder query using existing methodology.^4^ b) Count of patents that mention each CK1 isoform using the same querying methodology. c) Published inhibitors reported to inhibit CK1γ^1,15,16^.

While the CK1 family mediates numerous biological processes, the majority of its members play significant roles in the WNT signaling pathway^1,3,17^. The WNT signaling pathway is a fundamental biological process that governs cell fate, cell migration, cell polarity, and many other basic cellular functions^18,19^. Aberrant activation or inhibition of the WNT signaling pathway contributes to the pathogenesis of several forms of cancer, bone disease, and neurodegeneration^18^. Targeting various WNT pathway components, including other kinases, with small molecules has resulted in compounds that exhibit in vitro therapeutic efficacy, both when used as single agents and in combination with existing medications^19^. Recently, it was reported that the CK1γ kinases are necessary for maximal activation of WNT/β-catenin signaling^1,3^. Mechanistically, CK1γ phosphorylates the active form of the low-density lipoprotein receptor-related protein 6 (LRP6). LRP6 phosphorylation primes the receptor to form the WNT signalosome, a protein complex that suppresses β-catenin degradation, thereby driving the transcription of WNT-dependent genes^3,20,21^. Due to a scarcity of chemical tools, no studies have been done to demonstrate the effect of selective CK1γ inhibition on WNT-driven diseases. We aim to develop the requisite chemical tools to investigate the potential of CK1γ as a therapeutic target for treating diseases associated with aberrant WNT signaling.

Beyond WNT signaling, CK1γ isoforms play distinct roles in TNFα-induced necrotic cell death. CK1γ1 and CK1γ3 are reported to phosphorylate and enhance the kinase activity of receptor-interacting serine/threonine kinase 3 (RIPK3)^8^. Upon activation, RIPK3 phosphorylates mixed lineage kinase-like pseudokinase (MLKL), which then oligomerizes and develops pores in the plasma membrane of the cell. In contrast to CK1γ1 and CK1γ3, CK1γ2 was found to suppress necroptosis-promoted testicular aging by binding to and inhibiting RIPK3^22^. The involvement of necroptosis in dictating neuronal cell survival in neurodegenerative diseases suggests that CK1γ could be a promising therapeutic target for disrupting neuronal cell death. CK1γ subfamily members have also been implicated in human cytomegalovirus replication, sensitizing breast cancer cells to tamoxifen treatment, phosphorylation of p65, and in the production of sphingomyelin within cells^7,23,24,25,7^. However, many of these findings are reported in isolated publications with limited follow-up experimental validation to confirm that CK1γ kinase activity is solely responsible for the reported phenotype. Availability of a CK1γ subfamily chemical probe would enable dedicated studies of this nature.

Despite growing interest in the cellular roles of CK1γ, no selective inhibitors are currently available to pharmacologically investigate the functions outlined above. Current CK1γ inhibitors, such as FP1-24 and AKI-062a (Figure 1C), bind to and inhibit many kinases, including kinases essential for WNT signaling, which prevents them from being used to ask specific biological questions about CK1γ^1,26^. Specifically, when screened in the DiscoverX panel of 403 wild-type kinases at 1 µM, FP1-24 binds to 14 kinases with <10 percent of control (POC), including all CK1 isoforms. At the same concentration, AKI-062a binds 17 kinases with <10 POC, including GSK3β, an essential WNT pathway kinase. D4476, the inhibitor most frequently used to interrogate CK1γ biology, is a presumed pan-CK1 inhibitor with known potency against CK1α and CK1δ, but its cellular potency against the CK1γ isoforms has not been reported^15^. A scarcity of isoform-specific assays, especially those that assess cellular target engagement, has contributed to a paucity of high-quality chemical tools for the CK1γ subfamily. This emphasizes the urgent need for a potent, selective, and cell-permeable inhibitor to elucidate the cellular roles of CK1γ, as well as appropriate assays to characterize such an agent.

In the present work, we have addressed the lack of CK1γ chemical tools by developing the first potent and selective chemical probe, SGC-CK1γ-1. We identified this compound by optimizing for cellular potency and selectivity through a structure–activity relationship (SAR) study based on an underexplored scaffold found in literature^16^. To assess in-cell binding, we developed assays for each CK1γ isoform, including a NanoLuciferase (NLuc)-based thermal shift assay (NaLTSA) and a bioluminescence resonance energy transfer using NLuc (NanoBRET) cellular target engagement assay^27,28^. The availability of these assays addresses the current deficit in biological methods to evaluate CK1γ binding. Cumulatively, we have developed a suite of chemical tools and novel assays to enable research on this understudied set of kinases and demonstrated that these tools can be used to interrogate questions in diverse biological contexts.

## Results and Discussion

In 2012, Amgen optimized a 2-phenylamino-6-cyano-1*H*-benzimidazole-based hit into a promising CK1γ subfamily inhibitor (compound 1h, Figure 1C). We selected this molecule as a chemical starting point for our identification of CK1γ subfamily inhibitors because it exhibited good selectivity over CK1α, CK1δ, and GSK3β, decent biochemical potency for an unspecified CK1γ isoform (IC_50_ = 18 nM), and modest cellular potency in a substrate phosphorylation assay (IC_50_ = 700 nM)^16^. The initial selectivity screen for compound 1h revealed no off-targets, but only a small panel of 48 undisclosed kinases was evaluated^16^. While a seemingly promising starting point, the cellular potency, as well as the family and kinome-wide selectivity assessment, was insufficient to determine whether compound 1h satisfied the criteria established by the Structural Genomics Consortium (SGC) for a chemical probe^29,30^.

A crystal structure of a close analog of compound 1h that bears the cyanobenzimidazole structure revealed that the nitrile group engages in hydrogen bonding interactions mediated by a water network to Lys72, Glu86, and Tyr90 deep within the ATP binding pocket of CK1γ3. The 2-aminobenzimidazole makes a pair of hydrogen bonds with the backbone of Leu119^16^. We hypothesized that to enhance cellular potency, we would not modify the core to preserve these essential interactions and would instead explore various substituents around the distal phenyl ring of the molecule. This phenyl group is reported to engage in an edge-to-face hydrophobic interaction with Pro333, an amino acid within a region near the kinase domain that is unique to the CK1γ isoforms^16^. We investigated whether substituents around this ring system could enhance potency by forming new interactions with other amino acids near the solvent-exposed region of the compound, while not disrupting the interaction with Pro333. Additionally, at the solvent-exposed site of the compound, we added solubilizing groups that also served as chemical handles for later derivatization. A total of 19 analogs were synthesized through scheme 1, and 2 analogs were generated through alternate routes.

To assess in-cell affinity, we employed the existing CK1γ2 NanoBRET assay, which uses Promega tracer K10 to generate a medium signal window, but does not allow evaluation of CK1γ1 or CK1γ3 target engagement. These data are presented in Table 1. From our initial library, we found that solubilizing groups, such as piperazine (**3**) and morpholine (**4**), were tolerated at the *para* position in comparison to other chemical handles, such as a methyl ester (**1**). With a morpholine or piperazine fixed at the *para* position, we synthesized analogs bearing a diverse set of substituents at the *ortho* and *meta* positions, featuring varied steric and electronic properties. Compounds **2**, **4**, **7**, and **13** contained *ortho* substituents and all exhibited a dramatic loss in affinity compared to their molecular matched pairs with a hydrogen at that same position. Based on this observation, no further exploration was pursued at the *ortho* position.

**Table 1:**
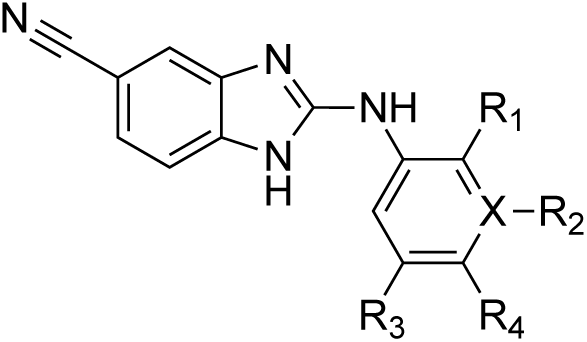
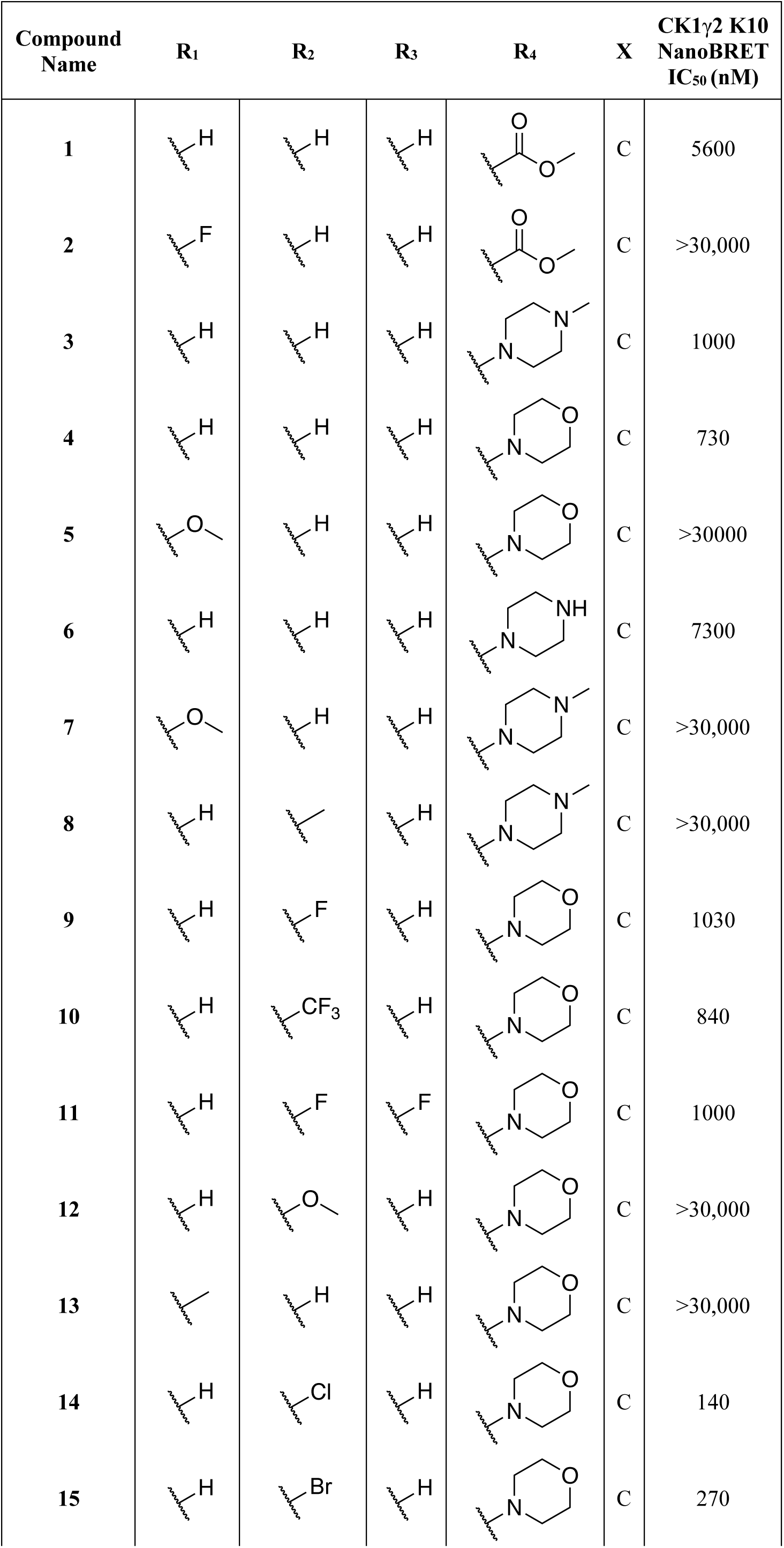
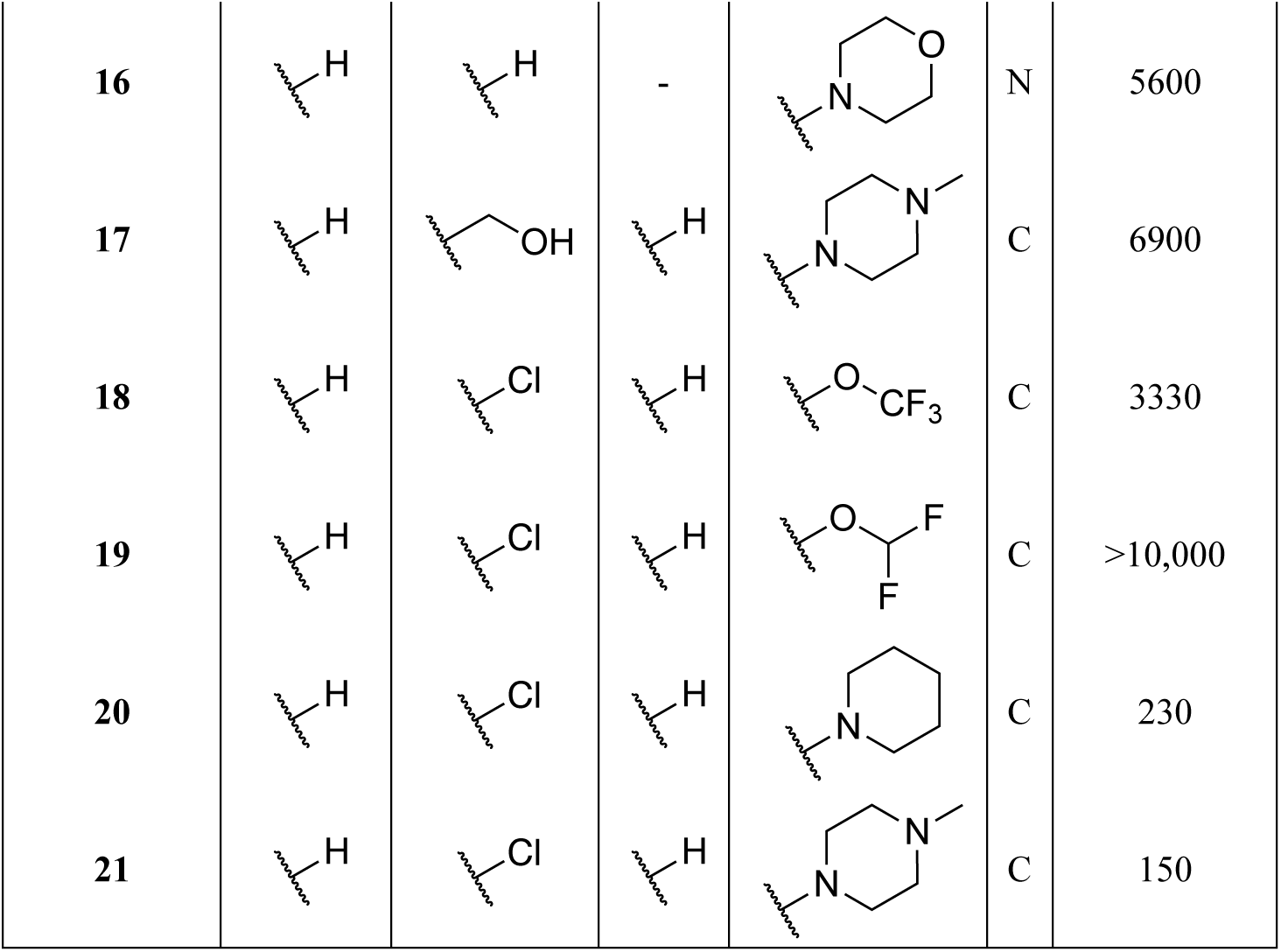
CK1γ NanoBRET data for analogs with a modified distal phenyl moiety.

### Analogs with Halogens at the *Meta* Position

At the *meta* position, electron-donating groups, such as a methyl (**8**) or methoxy group (**12**), were not tolerated (Table 1). In contrast, the addition of a strongly electron-withdrawing trifluoromethyl moiety (**10**) resulted in an analog with comparable cellular affinity to the corresponding hydrogen-containing analog. Furthermore, a phenyl to pyridine substitution by replacement of the 3-position carbon with a nitrogen in analog **16** resulted in reduced in-cell affinity. Next, to explore other electron-withdrawing groups of various sizes, we synthesized fluorinated analogs **9** and **12** with either a mono or di-substituted fluorine at the *meta* position. These substitutions were tolerated but did not increase CK1γ2 affinity. When larger halogens were introduced at the *meta* position in analogs **14** and **15**, we observed a significant reduction in cellular IC_50_ values (Table 1). We hypothesized that the substantial increase in affinity conferred by these larger halogens, which was absent in the fluorine-containing analogs, may be due to a halogen bond forming with a nitrogen or oxygen at the edge of the binding pocket. Consistent with this hypothesis, a co-folding experiment with compound **14** and CK1γ3 using the Boltz-2 model revealed that an oxygen on the backbone of Pro333 lies 3.3 Å from the chlorine of compound **14** (Supplementary Figure 1B)^31,32^. If a halogen bond forms, we would expect that **15** would create a stronger bond and, thus, have a significantly higher affinity to the protein due to the larger σ-hole potential of a bromine group. However, the chlorine in **14** may have an optimal geometric alignment for this specific putative halogen bond when compared to the larger bromine^33^. A co-crystal structure is needed to definitively validate any putative halogen bond occurring between **14** and CK1γ.

The high CK1γ2 affinity of compounds **14** and **15** inspired the design of analogs with further modifications at the *para* position. Specifically, incorporation of alternative solubilizing groups on compounds **17**–**21** resulted in compounds that were either equipotent with **14** or, in the case of compounds **18** and **19**, exhibited a significant loss of affinity. Compound **14** and its molecular matched pairs, bearing piperazine (**21**) and piperidine (**20**) moieties at the *para* position, demonstrated a significant increase in potency. These compounds were selected for further study.

### NaLTSA Assay Development

To begin to gather selectivity data, we first sought to assess the affinity of these compounds for the other CK1γ isoforms. To evaluate the cellular binding of **14** to all CK1γ isoforms, we performed a cellular thermal shift assay known as NaLTSA on cells transfected with a CK1γ isoform NLuc fusion plasmid. NaLTSA was chosen because there are currently no assays available to gauge binding to CK1γ1 or CK1γ3 in cells. In the presence of 30 μM of compound **14**, the melting temperatures shifted 3.5 °C for CK1γ1, 3.6 °C for CK1γ2, and 3.3 °C for CK1γ3 (Figure 2A). This temperature shift was not observed in the presence of cells expressing an unfused NLuc control vector (Supplementary Figure 2). To investigate whether the temperature shift was dose-dependent, we tested **14** in a nine-point dose–response format. The thermal shift decreased as the concentration of **14** was reduced (Figure 2B, C). The effect of the compound in NaLTSA for CK1γ2 was markedly less significant at lower concentrations compared to the corresponding CK1γ2 NanoBRET assay data, which was consistent with previous observations that NaLTSA requires a higher concentration of kinase inhibitors when compared to other assay formats to elicit the same effect^27^. The thermal shift data suggested that **14** demonstrated equal engagement with all CK1γ isoforms within cells. While the development of NaLTSA for the CK1γ subfamily represented a novel method to evaluate cellular binding, this assay required cell permeabilization with digitonin. Therefore, the thermal shift values recorded did not account for variables such as cellular ATP concentrations, permeation of the cell membrane, or protein localization within the cell.

**Figure 2.**
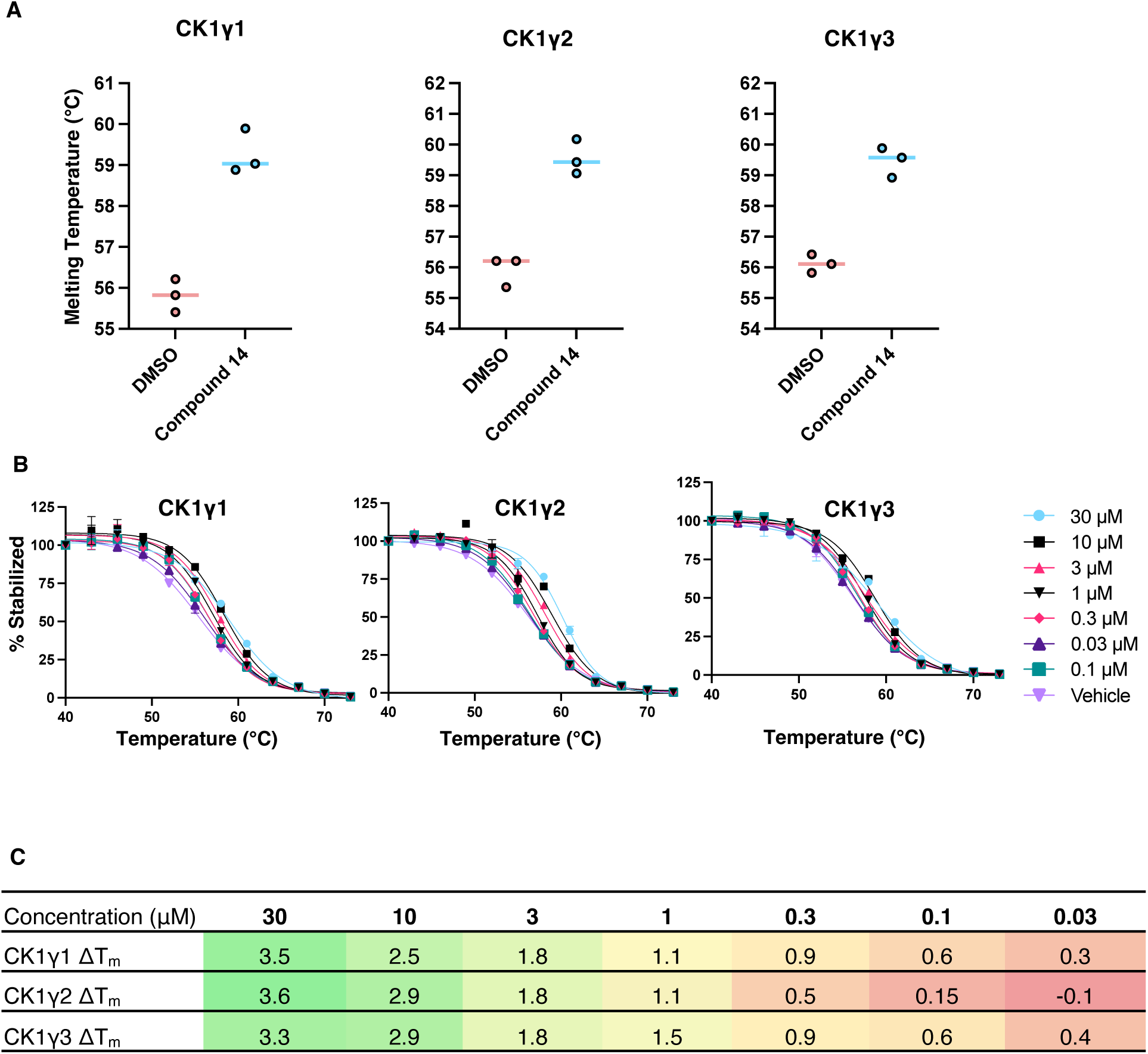
Development of a NaLTSA assay for the CK1γ isoforms. a) Melting temperatures of nanoluciferase-tagged CK1γ isoforms in the presence of either DMSO or 30 μM of compound **14** (N=3). All biological replicates were performed in technical duplicates. b) Validation of the thermal shift using a gradient dose range of compound **14,** starting at 30 µM and finishing at 0.1 µM. Data is from a single experiment run in duplicate. c) Melting temperatures determined for the nanoluciferase-tagged CK1γ isoforms in the dose range experiment with compound **14** in panel b. A color gradient was added to aid in data interpretation, with green indicating a significant shift, yellow an insignificant shift, and red no appreciable shift in melting temperature. All experiments were run in technical duplicates, and the color gradient represents data from three biological replicates.

### NanoBRET Assay Development

To better assess affinity in intact cells, we developed a NanoBRET tracer that enabled NanoBRET assays for the entire CK1γ subfamily. The development of NanoBRET tracers from promiscuous kinase inhibitors has enabled NanoBRET assays for a large fraction of the kinome^28^. However, to capture the portions of the kinome with limited chemical matter, selective or narrow-spectrum kinase inhibitors have been developed into NanoBRET tracers^34,35^. Consistent with this strategy, we utilized the piperazine on **21** as a chemical handle to link compound **21** to a NanoBRET 590 dye, thereby developing a CK1γ NanoBRET tracer (**23**) (Figure 3A). After synthesizing tracer **23,** we applied it to cells transfected with CK1γ1, 2, or 3, each with an N-terminal NLuc tag and obtained a significant BRET signal for all CK1γ kinases. To determine if the signal was caused by binding to the orthosteric site of the CK1γ kinases, we introduced 30 μM of **14** to compete with the signal caused by tracer binding (Figure 3B). Additionally, we determined tracer **23** EC_50_ values of 890 nM for CK1γ1, 370 nM for CK1γ2, and 590 nM for CK1γ3 through tracer dose–response experiments (Figure 3C). In tracer titration experiments carried out in dose–response format with compound **14**, the assays followed a linear trend between tracer concentration and IC_50_ values of **14** for all three kinases (Supplementary Figure 3). We then selected an optimal tracer concentration for each kinase that resulted in a signal window at least two-fold over background and below the EC_50_ value of the tracer for each kinase. Using these optimized assays, we screened all of our compounds as well as several CK1 inhibitors found in literature (structures in Supplementary Figure 5C) against all three kinases (Table 2)^15,36^.

**Figure 3.**
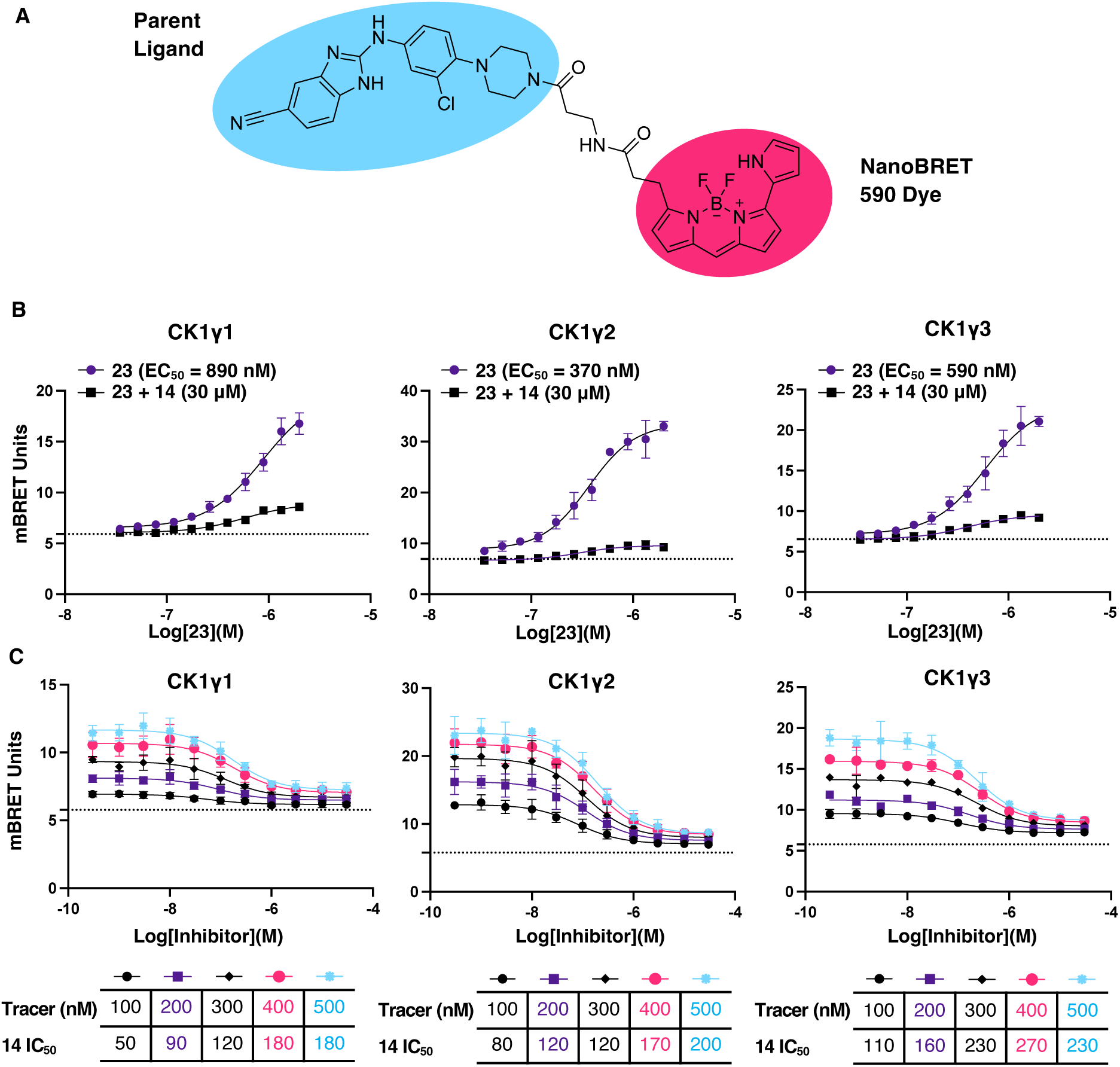
Development of NanoBRET assays for the CK1γ isoforms. a) Chemical structure of tracer **23** with the parent ligand in blue and NanoBRET 590 dye in magenta. b) 11-point dose–response experiments with tracer **23** in the presence and absence of compound **14**. The data is not background-subtracted, and a horizontal dotted line is present to indicate the background level. c) Tracer titrations with tracer **23** and compound **14**, with an IC_50_ displayed for each tracer concentration. All data are reported as N=3 ± SD. Data is not background-subtracted, and a horizontal dotted line is present to indicate the background.

**Table 2:** CK1γ1/2/3 NanoBRET data using tracer 23 for all analogs with potency against CK1γ2 in the K10 NanoBRET.

| Compound Name | NB<br>IC<sub>50</sub><br>CK1γ1 | NB<br>IC<sub>50</sub><br>CK1γ2 | NB<br>IC<sub>50</sub><br>CK1γ3 | Enzymatic<br>IC<sub>50</sub><br>CK1γ1 | Enzymatic<br>IC<sub>50</sub><br>CK1γ2 | Enzymatic<br>IC<sub>50</sub><br>CK1γ3 |
| --- | --- | --- | --- | --- | --- | --- |
| **Compound 1h** | 780 | 570 | 990 | 128 | 28 | 1661 |
| **4** | 910 | 920 | 2640 | 430 | 81 | 1298 |
| **5** | >30,000 | >30,000 | >30,000 | 5517 | 4171 | 2749 |
| **9** | 980 | 770 | 2760 | 212 | 34 | 656 |
| **10** | 1400 | 780 | 1350 | 120 | 34 | 656 |
| **11** | 920 | 1750 | 3300 | 121 | 25 | 976 |
| **14** | 180 | 180 | 230 | 28 | 7 | 958 |
| **15** | 580 | 110 | 110 | 40 | 10 | 700 |
| **20** | N.D. | N.D. | 300 | 30 | 9 | 475 |
| **21** | 170 | 220 | 180 | 36 | 14 | 431 |
| **D4476** | >30,000 | >30,000 | >30,000 | N.D. | N.D. | N.D. |
| **TAK-715** | >30,000 | >30,000 | >30,000 | N.D. | N.D. | N.D. |
| **PF-670462** | >30,000 | >30,000 | >30,000 | N.D. | N.D. | N.D. |
| **IWP-3** | >30,000 | >30,000 | >30,000 | N.D. | N.D. | N.D. |
| **CK1-IN-1** | >30,000 | >30,000 | N.D. | N.D. | N.D. | N.D. |

N.D. = Not determined

Given the homology of the active sites of these kinases, it is unsurprising that many of the SAR trends observed in the CK1γ2 NanoBRET assay were replicated when all three isoforms were evaluated using our NanoBRET assays^1^. Radiometric enzymatic assay data (Eurofins) for CK1γ1 and CK1γ2 replicated these observations (Table 2). However, the enzymatic assay for CK1γ3 appeared to yield significantly higher IC_50_ values when compared to the CK1γ3 NanoBRET assay (Table 2). We propose that this disparity may be due to the truncations and mutations, one of which is contained within the active site of the kinase, in the purified CK1γ3 used in the Eurofins enzymatic assay. For some biological contexts, it may be desirable to have an isoform-selective inhibitor^22,8,24^. However, in the context of WNT signaling, a pan CK1γ inhibitor would be more effective because these kinases act redundantly in phosphorylating LRP6^1,3,9^. Interestingly, none of the literature inhibitors of CK1 screened in the NanoBRET assay demonstrated affinity for the CK1γ isoforms, suggesting that distinct chemotypes may bind to CK1γ compared to the rest of the CK1 family. This is exemplified by inhibitors such as D4476, which has been used in several studies to specifically probe CK1γ function, yet does not show target engagement in our cellular assays^8,37,23,38^. From our screen, we identified several pan-CK1γ inhibitors that, if selective, could serve as more effective research tools for understanding CK1γ biology than existing inhibitors in the literature. To this end, we prioritized **14** and **21** as CK1γ subfamily chemical probe candidates, as they were the most potent compounds and demonstrated equal affinity for all three isoforms.

### Evaluation of Kinome Selectivity

To identify off-target kinases of compound **14**, we tested its binding to a panel of 192 kinases via a NanoBRET assay selectivity panel (K192) at a fixed concentration of 1 μM. We found that **14** demonstrated excellent selectivity across three biological replicates of K192 assays. The only kinase with greater than 30% fractional occupancy at 1 μM was CK1γ2 (74%) (Figure 4 A, C). It is important to note that CK1γ2 was the sole representative from the CK1γ subfamily in the K192 panel. This initial selectivity profile for compound **14** prompted us to screen compounds **14** and **21** in a larger NanoBRET panel of 240 kinases (K240). Compound **14** maintained strong in-cell selectivity with only two kinases demonstrating greater than 30% fractional occupancy: MOK (37%) and CK1γ2 (66%) (Figure 4A, B). Screening of **14** against MOK in a NanoBRET assay with an 11-point dose–response format revealed that the IC_50_ is greater than 5 μM; therefore, MOK is not a significant off-target of **14** (Supplementary Figure 5). The IC_50_ of **14** in the CK1γ2 NanoBRET assay with tracer K10, the same tracer used in the K192 and K240 panels, was previously determined to be 140 nM (Table 1).

**Figure 4.**
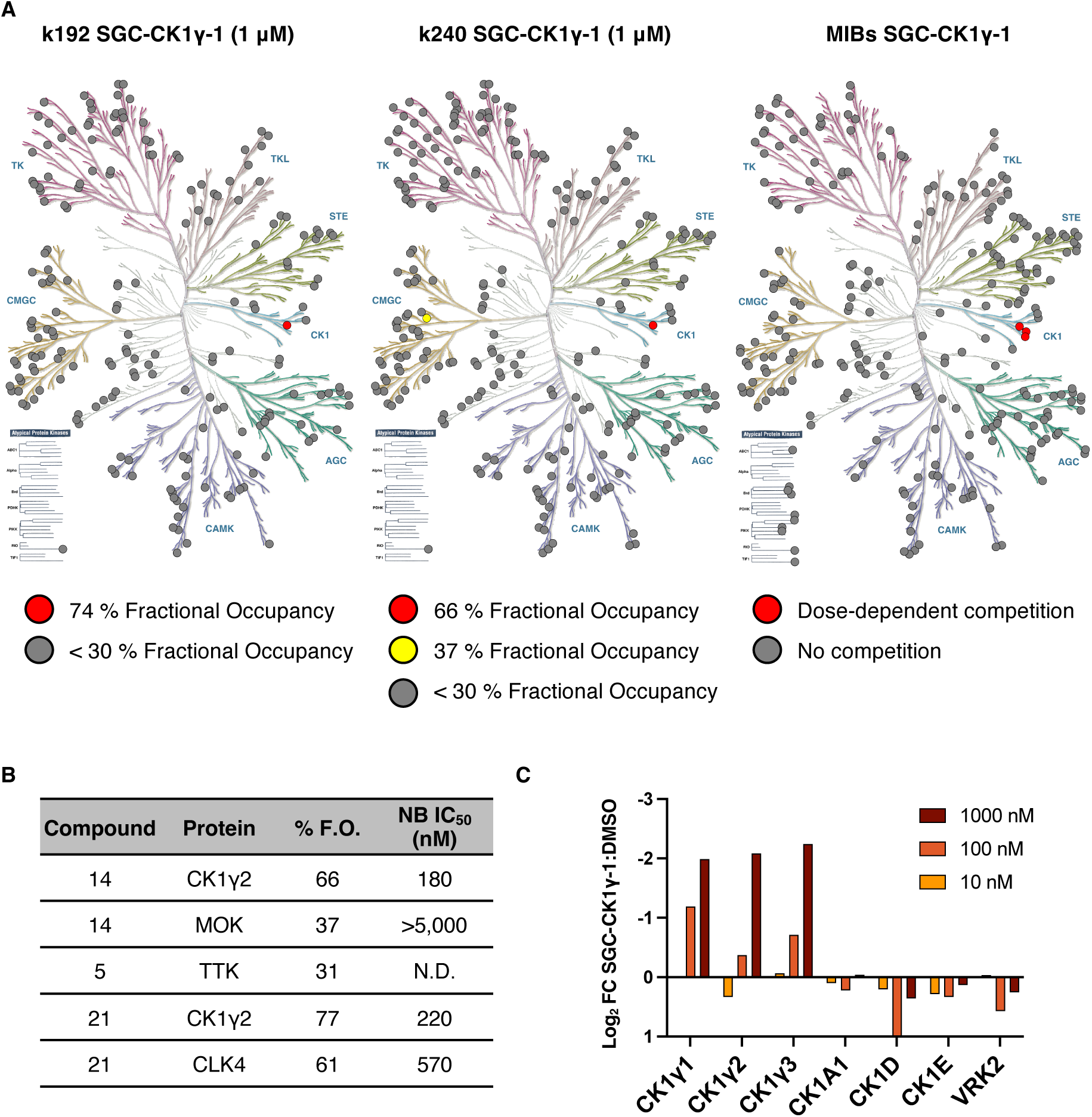
Demonstration of the in-cell selectivity of compound **14**. a) Dendrograms displaying all kinases inhibited in red, partially inhibited in yellow, and not inhibited but tested in grey for the K192 (N=3), K240 (N=1), and MIBs assays (N=1). b) All kinases identified across both selectivity screens and the follow-up IC_50_ determinations in the respective NanoBRET assays for compounds **14 (**SGC-CK1γ-1), **5 (**SGC-CK1γ-1N), and **21**. F.O. = fractional occupancy. NB = NanoBRET. c) Pulldown fold change data for all CK1γ isoforms treated with 10, 100, or 1000 nM of compound **14 (**SGC-CK1γ-1) normalized to DMSO treated lysate.

Selectivity screening of compound **21** confirmed it also had good in-cell kinome-wide selectivity, with two targets exceeding 30% fractional occupancy in the K192 assay: CLK4 (49%) and CK1γ2 (73%) (Supplementary Figure 6). In the subsequent K240, several kinases demonstrated fractional occupancy greater than 30%: CDK6 (33%), CDKL2 (39%), CDK9 (42%), CDK2 (45%), CLK4 (61%), and CK1γ2 (77%) (Supplementary Figure 6B). However, many of these kinases were present in both the K192 and the K240, CLK4 and CK1γ2 were the only kinases hit in both screens. In the dose–response follow-up NanoBRET assay, compound **21** bound CLK4 with an IC_50_ of 570 nM, making it a significant off-target of this compound (Figure 4B). Interestingly, **21** did not exhibit affinity for CLK1 or CLK2, which were included in the K192 and K240. Compound **21** may serve as a chemical starting point for the development of future isoform-specific CLK4 inhibitors. It can also be used in tandem with compound **14** to probe biology driven by CK1γ since the two compounds are potent CK1γ binders and have well annotated, yet largely non-overlapping kinome-wide profiles. Compound **14** demonstrated minimal affinity for CLK4 when tested in the corresponding NanoBRET assay in an 11-point dose–response format (Supplementary Figure 5B).

Having defined the cellular selectivity of compound **14** using large kinase-based panels, we sought to further interrogate its selectivity by performing an orthogonal multiplexed inhibitor bead and mass spectrometry (MIB/MS) screen. MIB/MS samples many human protein kinases, typically between 250 and 350 kinases, based on their expression in cell lysates, and thus includes kinases also in the K240 panel, as well as several not represented. Using a MIB matrix of six ATP competitive inhibitors, MIB/MS can pull down 75–80% of the kinome expressed in a human cell lysate^39,40^. Upon introducing compound **14** to the bead-lysate mixture, we noted remarkable selectivity, with only dose-dependent competition observed for CK1γ1, CK1γ2, and CK1γ3 (Figure 4A, C). Based on the MIB/MS analysis, we were able to pull down 278 human protein kinases. Cumulatively, across both the MIB/MS and k240, we were able to assess the binding of compound **14** to 372 kinases, 145 of which were screened in both assay formats. Using this dual profiling approach, we have identified compound **14** as our optimal candidate for designation as a CK1γ chemical probe for all three CK1γ proteins: CK1γ1, CK1γ2, and CK1γ3.

At this point in our study, we had confirmed that compound **14** demonstrates a cellular IC_50_ of <1 μM, a biochemical IC_50_ <100 nM, good aqueous solubility in DLS, and excellent kinome-wide selectivity (Supplementary Figure 9). The remaining requirement for a chemical probe, as defined by the Structural Genomics Consortium, is to identify a negative control compound with a similar chemical structure but significantly reduced affinity (100-fold reduced affinity when compared to the chemical probe) for the target protein^41^. To determine whether it met the criteria to serve as a negative control to pair with **14**, **5** was selected for K240 screening. Compound **5** bears structural similarity to **14** and exhibits no affinity for all CK1γ isoforms (Table 2). In the K240 screen, only one kinase was identified to manifest >30% fractional occupancy: TTK (37%) (Supplementary Figure 6). Given the excellent kinome-wide selectivity of **5**, we chose it as a negative control compound to be used alongside chemical probe **14**. Hereafter, compound **14** was termed SGC-CK1γ-1 and compound **5** was termed SGC-CK1γ-1N.

### Probe Applications in WNT Signaling and Human Cytomegalovirus Replication

With both the probe (SGC-CK1γ-1) and negative control (SGC-CK1γ-1N) in hand, we assayed CK1γ inhibition in functionally relevant biological assays: inhibition of WNT signaling and inhibition of human cytomegalovirus replication.

First, we sought to reproduce the reduction of LRP5/6 phosphorylation elicited by the less selective inhibitor FP1-24 and reported elsewhere^1^. Phosphorylation of LRP5/6 at T1479 by CK1γ upon WNT binding is a crucial step in recruiting AXIN and other proteins that mediate intracellular signaling that results in gene transcription (Figure 5A)^1^. Thus, CK1γ-driven phosphorylation is essential for WNT function and explains the agonistic effect that CK1γ has on WNT signaling^1,9^. We demonstrated a significant loss of LRP6 phosphorylation at T1479 when HEK293 cells were dosed with 5 μM of SGC-CK1γ-1 (Figure 5B, C). This reduction in LRP6 phosphorylation did not occur in the presence of 5 μM SGC-CK1γ-1N. When the dose was increased to 10 μM of SGC-CK1γ-1, a near-complete loss of phospho-LRP6 was observed. When tested in a CellTiter-Glo viability assay, SGC-CK1γ-1 did not impact cell viability when HEK293 cells were dosed at 10 μM for 24 hours or 48 hours (Supplementary Figures 7A and 7B). Treatment of a panel of 60 cancer cell lines with 10 µM of SGC-CK1γ-1 was found not to reduce cell viability, supporting the compound as non-toxic and that inhibition of CK1γ is not sufficient to elicit an anti-cancer phenotype (Supplementary Figure 7C). Since CK1γ is not the only reported kinase to influence phosphorylation of LRP6 at T1479^42,43^, an absolute loss of phosphorylated protein was not expected. Notably, similar results were obtained following treatment with 10 μM of FP1-24, an inhibitor with an enzymatic IC_50_ of <10 nM for all CK1γ isoforms^1,44^.

**Figure 5.**
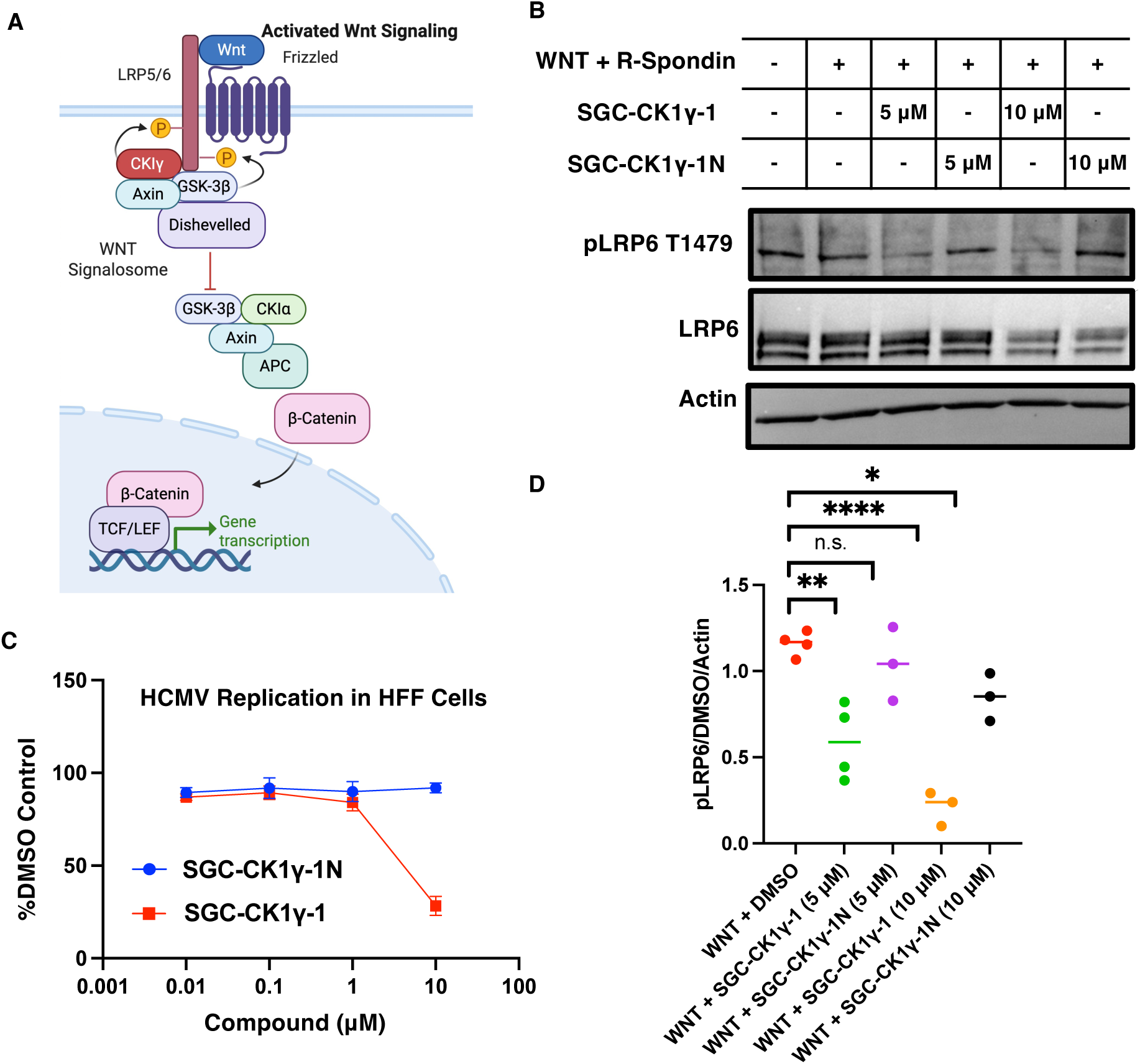
SGC-CK1γ-1 disrupts LRP6 phosphorylation. Schematic of the WNT signaling pathway. b) Western blot analysis of LRP6 phosphorylation at T1479 in HEK293 cells treated with WNT activating molecules WNT3A and R-Spondin-1 in the presence of DMSO, SGC-CK1γ-1, or SGC-CK1γ-1N for 2 hours prior to lysis. c) HFF cells were infected with HCMV strain Merlin (R1111) and treated with various concentrations of either SGC-CK1γ-1 or SGC-CK1γ-1N, or the corresponding volume of DMSO, at the time of infection (N=3). Error bars represent SD. d) Quantification of Western blots with all data normalized to cells treated with only DMSO and the actin control. Statistical significance was evaluated by Student’s *t*-test. For all quantifications: **** *p* < 0.0001, *** *p* < 0.005, ** *p* < 0.005, and * *p* < 0.05. n.s. = not significant.

Finally, it has been suggested that CK1γ proteins are required for replication of human cytomegalovirus (HCMV)^45,7^. In our hands, SGC-CK1γ-1 elicited anti-HCMV replication effects (Figure 5D) with no noticeable impact on cell viability when used at 10 µM (Supplemental Figure 8A, B). Notably, SGC-CK1γ-1N did not demonstrate any observable anti-HCMV effects, even when used at high concentrations (Figure 5D). The efficacy of these anti-HCMV effects supports that further optimization and/or co-treatment would be required to effectively inhibit HCMV replication. Regardless, these results support previous data suggesting that CK1γ proteins may play at least a minor role in promoting HCMV replication^7,45^.

This initial finding is intriguing but requires additional exploration to elucidate the mechanism driving this activity. It is possible that a CK1γ isoform phosphorylates any of the several HCMV proteins produced during replication that require phosphorylation for their function. These include the immediate early HCMV proteins IE1 and IE2, whose functions are essential for HCMV replication. This is supported by our observation that SGC-CK1γ-1 had no anti-HCMV activity when added to HCMV infected cells after expression of IE1 and IE2 (24 hours post infection) (Supplemental Figure 8C)^46,47^. Identification of CK1γ substrates, possibly mediated through the use of SGC-CK1γ-1 in phosphoproteomics-based studies, would help identify protein motifs that these underexplored kinases may phosphorylate in virus infected cells^47^.

## Conclusions

Our SAR campaign yielded a potent and selective compound, SGC-CK1γ-1, that satisfies the criteria for a chemical probe. Through our dual selectivity profiling approach, including both MIB/MS and a large panel of kinase NanoBRET assays, we have provided an unprecedented interrogation of the kinome-wide selectivity of SGC-CK1γ-1 in both intact and lysed cells. Furthermore, we developed the first NanoBRET assays for CK1γ1 and CK1γ3 and significantly improved the signal window of the existing CK1γ2 NanoBRET assay through the use of a single tracer. NanoBRET, alongside NaLTSA, represents cellular assays that are amenable to high-throughput formats for future drug discovery efforts for CK1γ. Alongside its inactive control derivative, SGC-CK1γ-1N, SGC-CK1γ-1 can be used to explore the various putative functions of CK1γ, including its role in LRP6 phosphorylation in the WNT signaling pathway and potential contribution to HCMV replication. It can further be used to refine the substrate scope of this family of poorly characterized kinases, which may illuminate previously unknown functions of the CK1γ subfamily and initiate research in poorly understood areas of CK1γ research, such as necroptosis.

## Materials and Methods

### Chemistry

SGC-CK1γ-1 and its analogs were synthesized through a nucleophilic aromatic substitution (SNAr) reaction between 2-chloro-1*H*-benzo[*d*]imidazole-5-carbonitrile, representing the conserved left terminal of the compound, and an aniline with a varying structure.

The chemical synthesis of compounds **1-5, 7-21** (**14** is SGC-CK1γ-1 and **5** is SGC-CK1γ-1N) is outlined in Scheme 1, the synthesis of the literature compound 1h is outlined in Scheme 2, and the synthesis of compound **23** is outlined in Scheme 3.

#### Scheme 1

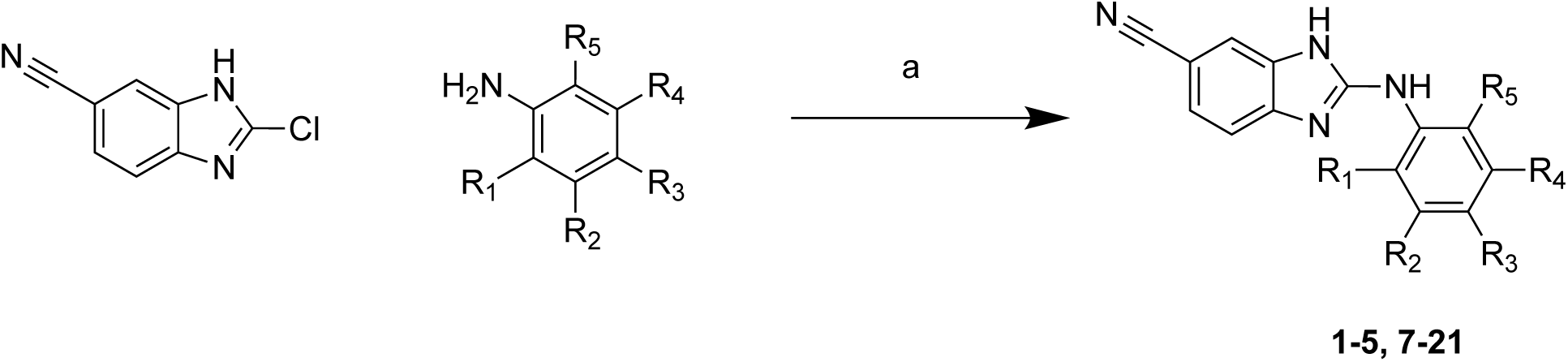

Scheme 1: Synthesis of analogs **1-5, 7-21**

*^a^*Reagents and conditions: (a) methanesulfonic acid, ACN, 140 °C, 30 min, 2-60%.

To synthesize compound 1h, we used an alternate route starting with compound **1**, synthesized using Scheme 1, and methyl lithium to achieve the *tert*-butoxy group on the right terminal of compound 1h.

#### Scheme 2

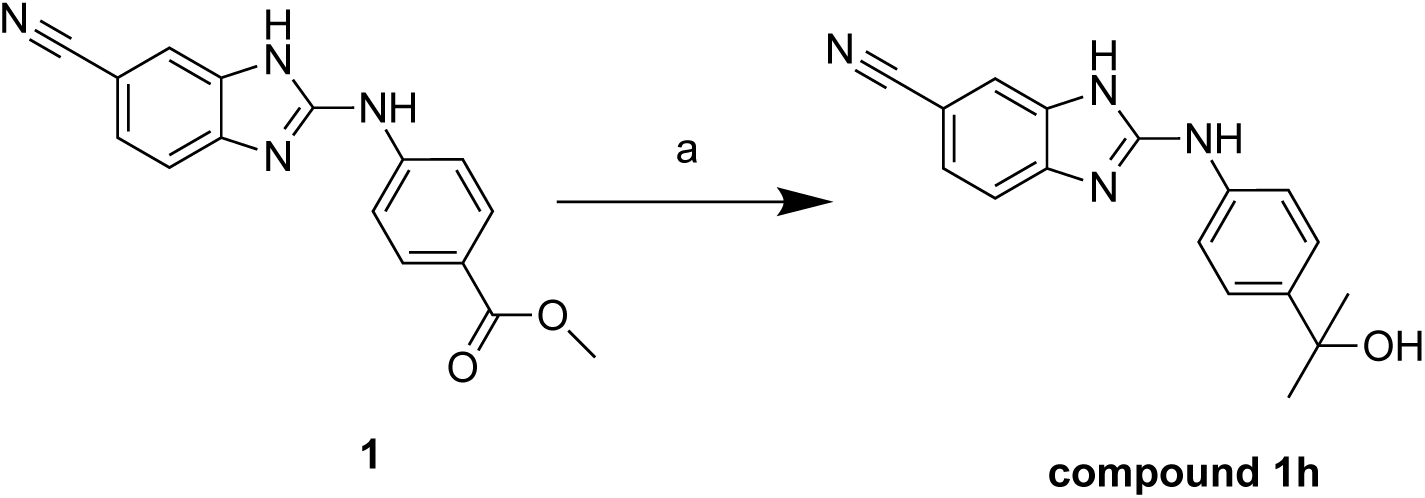

Scheme 2: Synthesis of analogs **compound 1h**

*^a^*Reagents and conditions: (a) methyl lithium in ether, THF, -78 °C, 2 h, 10%.

#### Scheme 3

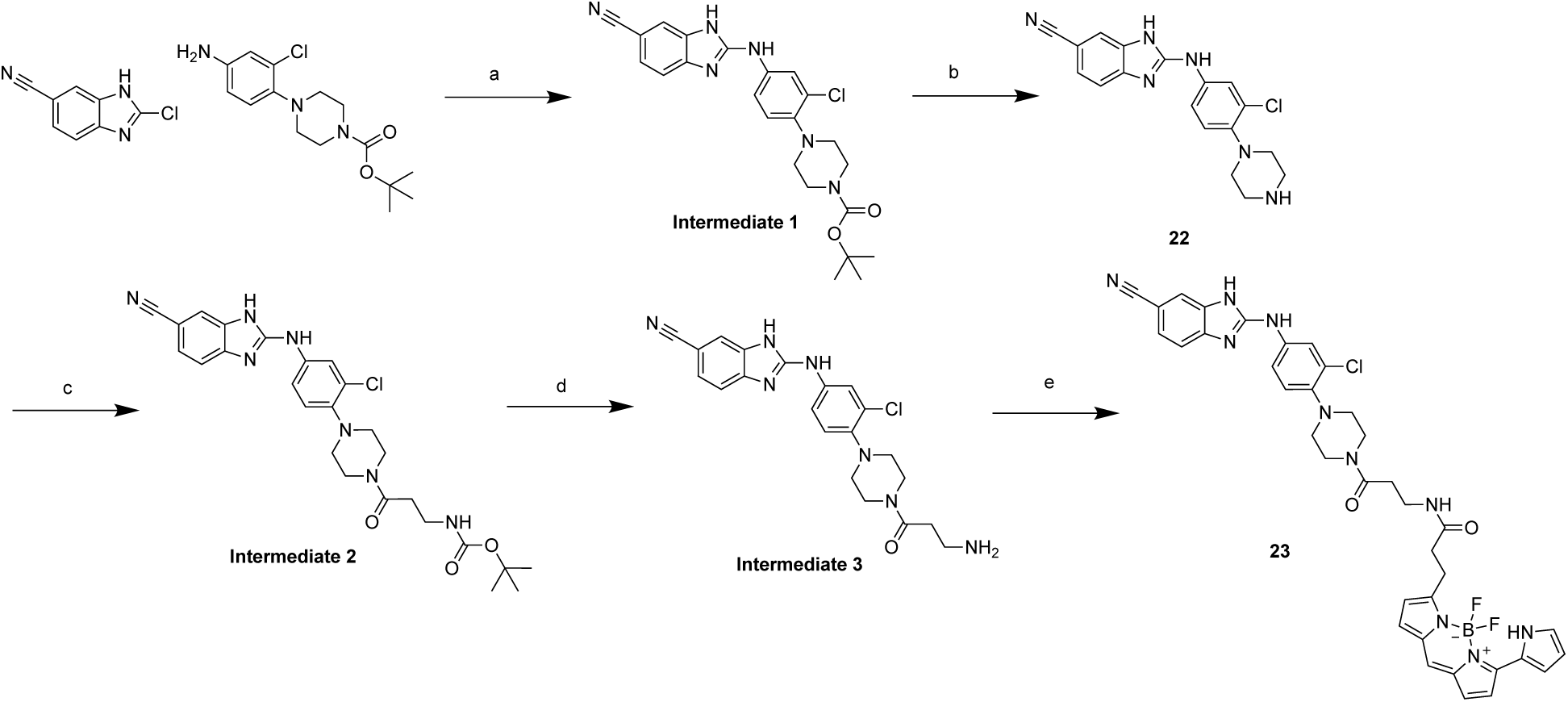

Scheme 3: Synthesis of tracer **23**

*^a^*Reagents and conditions: (a) methanesulfonic acid, ACN, 140 °C, 30 min. (b) DCM, TFA, rt, 2 hours, 16.7 % over two steps. (c) HATU, DIPEA, 3-((*tert*-butoxycarbonyl)amino)propanoic acid, DMF, rt, o.n.. (d) DCM, TFA, rt, 2 hours. (e) 2,5-dioxopyrrolidin-1-yl 3-(5,5-difluoro-7-(1*H*-pyrrol-2-yl)-5*H*-5λ^4^,6λ^4^-dipyrrolo[1,2-*c*:2’,1’-*f*][1,3,2]diazaborinin-3-yl)propanoate, DIPEA, DMF, rt, 2 h, 7% over three steps.

### Experimental Section

#### Chemistry: General Information

All purchased reagents and solvents were used without further purification or characterization. Reaction temperatures are reported in degrees Celsius (°C). All reactions without a listed temperature occurred at room temperature (25°C). All solvent removal was accomplished using a rotary evaporator under reduced pressure. The following abbreviations are used in schemes and/or experimental procedures: μmol (micromoles), mmol (millimoles), mg (milligrams), equiv (equivalent(s)), r.t. (room temperature), min (minutes), sec (seconds), and h (hours). Compound purity and identity were confirmed using ^1^H NMR and/or additional microanalytical data for intermediates and final compounds. ^1^H NMR and ^13^C NMR spectra were obtained in either DMSO-*d*_6_ or CD_3_OD and recorded using Bruker instruments. The magnet’s strength for the NMR spectra is indicated in each line listing. The peak positions, listed in parts per million (ppm), are calibrated to the indicated deuterated solvent. Coupling constants (*J* values) are listed as follows: singlet (s), doublet (d), doublet of doublets/triplets/quartets (dd/dt/dq), doublet of doublet of doublets (ddd), triplet (t), triplet of doublets/triplets (td/tt), quartet (q), quartet of doublets (qd), pentet (p), and multiplet (m). Preparative HPLC was performed using an Agilent 1100 Series System equipped with a Phenomenex Luna Phenyl-Hexyl column (5 μm particle size, 100 Å pore size, 75 × 30 mm) or an Agilent 1260 Infinity II LC System equipped with a Phenomenex C18 Phenyl-Hexyl column (30 °C, 5 μm particle size, 75 x 30 mm). For LCMS analysis, an Agilent 1290 Infinity II LC System equipped with an Agilent Infinity Lab PoroShell 120 EC-C18 column (30°C, 2.7 μm particle size, 2.1 × 50 mm), eluent 10−90% CH_3_CN in water with 0.2% formic acid (v/v), and flow rate of 1 mL/min, was used. HPLC analysis determined that all reported final compounds were greater than 95% pure.

#### General Procedure A

To a microwave vial was added a mixture of 2-chloro-1*H*-benzo[*d*]imidazole-5-carbonitrile (2.0 equiv), amine (1.1 equiv), and methanesulfonic acid (1.2 equiv) in ACN. The reaction mixture was then heated in a microwave to 140 °C for 30 min. The reaction mixture was cooled to r.t. and concentrated *in vacuo*. The crude residue was purified via reverse phase chromatography on a C18 column (5-100% acetonitrile (ACN) in H_2_O + 0.1% TFA) to yield the title compounds.

#### General Procedure B

To a flask was added a mixture of 2-chloro-1*H*-benzo[*d*]imidazole-5-carbonitrile (2.0 equiv), amine (1.1 equiv), and potassium dihydrogen phosphate (1.0 equiv) in 1-butanol. The reaction mixture was heated to 120 °C for 12 h. The reaction mixture was cooled to r.t. and concentrated *in vacuo*. The crude residue was added to a vial with DCM and TFA (5.0 equiv). The crude residue was concentrated *in vacuo* and purified via reverse phase chromatography on a C18 column (5-100% ACN in H_2_O + 0.5% TFA) to yield the title compounds.

#### Methyl 4-((6-cyano-1*H*-benzo[*d*]imidazol-2-yl)amino)benzoate (1)

To a microwave vial was added 2-chloro-1*H*-benzo[*d*]imidazole-5-carbonitrile (50 mg, 0.28 mmol), methyl 4-aminobenzoate (47 mg, 0.31 mmol), and methanesulfonic acid (32 mg, 0.34 mmol) in ACN (2 mL). The reaction mixture was heated in a microwave to 140 °C for 30 min. The reaction was cooled to r.t. and filtered to yield the title compound as a white powder (50.4 mg, 62% yield). ^1^H NMR (400 MHz, DMSO-*d_6_*) δ 11.13 (s, 1H), 8.00 (d, *J* = 8.9 Hz, 2H), 7.84 (s, 1H), 7.80 (dd, *J* = 8.9, 2.1 Hz, 2H), 7.56 (s, 2H), 3.84 (s, 3H). ^13^C NMR (100 MHz, DMSO-*d_6_*) δ 166.25, 150.82, 143.18, 138.15, 134.74, 131.14 (2C), 126.75, 124.47, 120.21 (2C), 119.48, 116.91, 113.94, 104.12, 52.42. HPLC Purity >96%. LCMS calculated for C_16_H_13_N_4_O_2_ [M + H]^+^: 293.1. Found: 292.9.

#### Methyl 4-((5-cyano-1*H*-benzo[*d*]imidazol-2-yl)amino)-3-fluorobenzoate (2)

The reaction was carried out according to general procedure A with 2-chloro-1*H*-benzo[*d*]imidazole-5-carbonitrile (50 mg, 0.28 mmol), methyl 4-amino-3-fluorobenzoate (52 mg, 0.31 mmol), and methanesulfonic acid (32 mg, 0.34 mmol) in ACN (2 mL). The mixture was microwaved at 140 °C for 30 min to yield the TFA salt form of the compound as a light brown powder (34 mg, 39.0 % yield). ^1^H NMR (400 MHz, DMSO-*d_6_*) δ 8.56 (t, *J* = 8.2 Hz, 1H), 7.92 – 7.86 (m, 2H), 7.81 (dd, *J* = 11.8, 1.9 Hz, 1H), 7.60 (dd, *J* = 8.2, 0.7 Hz, 1H), 7.55 (dd, *J* = 8.2, 1.5 Hz, 1H), 3.87 (s, 3H). ^13^C NMR (214 MHz, DMSO-*d_6_*) δ 168.27, 161.25, 161.09, 154.51, 153.37, 135.72 (2C), 128.15, 125.71, 123.48 (2C), 121.73, 118.72, 118.62, 55.28. HPLC Purity >99%. LCMS calculated for C_16_H_12_FN_4_O_2_ [M + H]^+^: 311.9. Found: 311.1.

#### 2-((4-(4-methylpiperazin-1-yl)phenyl)amino)-1H-benzo[d]imidazole-6-carbonitrile (3)

The reaction was carried out according to general procedure A with 2-chloro-1*H*-benzo[*d*]imidazole-5-carbonitrile (150 mg, 0.845 mmol), 4-(4-methylpiperazin-1-yl)aniline (178 mg, 0.929 mmol), and methanesulfonic acid (97.4 mg, 1.01 mmol) in ACN (4 mL). The mixture was microwaved at 140 °C for 30 min to yield the TFA salt form of the title compound as a dark brown amorphous solid (25 mg, 8.9% yield). ^1^H NMR (400 MHz, CD_3_OD) δ 7.53 (t, *J* = 1.1 Hz, 1H), 7.41 – 7.29 (m, 4H), 7.05 – 6.96 (m, 2H), 3.21 – 3.14 (m, 4H), 2.67 – 2.60 (m, 4H), 2.36 (s, 3H).^13^C NMR (214 MHz, CD_3_OD) δ 169.70, 157.23, 149.38, 135.45, 127.46, 124.07 (2C), 122.67, 120.28 (2C), 118.05, 115.41, 104.99, 56.49 (2C), 45.71 (2C), 41.70. HPLC purity > 99%. LCMS calculated for C_16_H_21_N_6_ [M + H]^+^: 333.2. Found: 333.1.

#### 2-((4-morpholinophenyl)amino)-1H-benzo[d]imidazole-6-carbonitrile (4)

The reaction was carried out according to general procedure A with 2-chloro-1H-benzo[*d*]imidazole-5-carbonitrile (50 mg, 0.28 mmol), 4-morpholinoaniline (55 mg, 0.31 mmol), and methanesulfonic acid (32 mg, 0.34 mmol) in ACN (2 mL). The mixture was microwaved at 140 °C for 30 min to yield the double TFA salt form of the title compound as a brown amorphous solid (10.7 mg, 6.9% yield). ^1^H NMR (400 MHz, CD_3_OD) δ 7.70 (dd, *J* = 1.5, 0.7 Hz, 1H), 7.64 (dd, *J* = 8.3, 1.5 Hz, 1H), 7.50 (dd, *J* = 8.3, 0.7 Hz, 1H), 7.40 – 7.32 (m, 2H), 7.18 – 7.10 (m, 2H), 3.89 – 3.82 (m, 4H), 3.25 – 3.20 (m, 4H). ^13^C NMR (100 MHz, CD_3_OD) δ 151.11, 150.96, 133.44, 130.23, 127.89, 126.55, 125.59 (2C), 118.16, 116.61 (2C), 115.00, 112.15, 106.58, 66.37 (2C), 48.95 (2C). HPLC Purity >95%. LCMS calculated for C_18_H_18_N_5_O [M + H]^+^: 320.1. Found: 319.9.

#### 2-((2-methoxy-4-morpholinophenyl)amino)-1H-benzo[d]imidazole-6-carbonitrile (5)

The reaction was carried out according to general procedure A with 2-chloro-1H-benzo[*d*]imidazole-5-carbonitrile (50 mg, 0.28 mmol), 2-methoxy-4-morpholinoaniline (64 mg, 0.31 mmol), and methanesulfonic acid (32 mg, 0.34 mmol) in ACN (2 mL). The mixture was microwaved at 140 °C for 30 min to yield the double TFA salt form of the title compound as a light brown amorphous solid (6.9 mg, 4.2% yield). ^1^H NMR (400 MHz, CD_3_OD) δ 7.69 (dd, *J* = 1.5, 0.7 Hz, 1H), 7.64 (dd, *J* = 8.3, 1.5 Hz, 1H), 7.49 (dd, *J* = 8.3, 0.7 Hz, 1H), 7.28 (d, *J* = 8.7 Hz, 1H), 6.76 (d, *J* = 2.5 Hz, 1H), 6.67 (dd, *J* = 8.7, 2.6 Hz, 1H), 3.85 (m, 7H), 3.27 – 3.21 (m, 4H). ^13^C NMR (100 MHz, CD_3_OD) δ 155.30, 153.30, 151.52, 133.36, 130.15, 127.85, 127.69, 118.15, 114.90, 114.24, 112.06, 107.43, 106.55, 99.75, 66.44, 54.89 (2C), 48.81 (2C). HPLC Purity >99%. LCMS calculated for C_19_H_20_N_5_O_2_ [M + H]^+^: 350.2. Found: 350.1.

#### 2-((4-(piperazin-1-yl)phenyl)amino)-1H-benzo[d]imidazole-6-carbonitrile (6)

The reaction was carried out according to general procedure B with 2-chloro-1H-benzo[*d*]imidazole-5-carbonitrile (200 mg, 1.13 mmol), *tert*-butyl 4-(4-aminophenyl)piperazine-1-carboxylate (344 mg, 1.24 mmol), and potassium dihydrogen phosphate (153 mg, 1.13 mmol) in 1-butanol (4 mL). After concentrating the crude reaction mixture, dichloromethane (3 mL) was added with trifluoracetic acid (642 mg, 5.63 mmol) to yield the double TFA salt form of the title compound as a dark purple amorphous solid (42 mg, 11.7 % yield over two steps) after reverse phase purification. ^1^H NMR (400 MHz, CD_3_OD) δ 7.72 (dd, *J* = 1.5, 0.6 Hz, 1H), 7.65 (dd, *J* = 8.3, 1.5 Hz, 1H), 7.51 (dd, *J* = 8.3, 0.7 Hz, 1H), 7.45 – 7.36 (m, 2H), 7.24 – 7.16 (m, 2H), 3.49 (dd, *J* = 6.6, 3.6 Hz, 4H), 3.41 (dd, *J* = 6.6, 3.5 Hz, 4H). ^13^C NMR (100 MHz, CD_3_OD) δ 151.06, 149.90, 133.64, 130.40, 127.87, 127.78, 125.57 (2C), 118.16, 117.73 (2C), 115.05, 112.20, 106.59, 46.13 (2C), 43.27 (2C). HPLC Purity >99%. LCMS calculated for C_18_H_19_N_6_ [M + H]^+^: 319.2. Found: 318.8.

#### 2-((2-methoxy-4-(4-methylpiperazin-1-yl)phenyl)amino)-1H-benzo[d]imidazole-6-carbonitrile (7)

The reaction was carried out according to general procedure A with 2-chloro-1H-benzo[*d*]imidazole-5-carbonitrile (50 mg, 0.28 mmol), 2-methoxy-4-(-methylpiperazin-1-yl)aniline (69 mg, 0.31 mmol), and methanesulfonic acid (32 mg, 0.34 mmol) in ACN (2 mL). The mixture was microwaved at 140 °C for 30 min to yield the double TFA salt form of the title compound as a light purple amorphous solid (19.2 mg, 11.3% yield). ^1^H NMR (400 MHz, DMSO-*d*_6_) δ 10.60 (s, 1H), 10.26 (s, 1H), 7.77 (d, *J* = 1.5 Hz, 1H), 7.63 (dd, *J* = 8.3, 1.6 Hz, 1H), 7.48 (d, *J* = 8.2 Hz, 2H), 6.80 (d, J = 2.5 Hz, 1H), 6.67 (dd, J = 8.8, 2.5 Hz, 1H), 3.95 (m, 2H), 3.83 (s, 3H), 3.56 (s, 2H), 3.17 (s, 2H), 3.03 (s, 2H), 2.89 (s, 3H). ^13^C NMR (214 MHz, DMSO-*d*_6_) δ 156.97, 154.60, 152.97, 130.34, 122.44, 121.51, 120.13, 118.75, 118.50, 117.37, 115.79, 110.86, 107.61, 103.98, 58.98, 55.44 (2C), 48.77 (2C), 45.23, 43.56. HPLC Purity >99%. LCMS calculated for C_20_H_23_N_6_O [M + H]^+^: 363.2. Found: 362.9.

#### 2-((3-methyl-4-(4-methylpiperazin-1-yl)phenyl)amino)-1H-benzo[d]imidazole-6-carbonitrile (8)

The reaction was carried out according to general procedure A with 2-chloro-1H-benzo[*d*]imidazole-5-carbonitrile (50 mg, 0.28 mmol), 3-methyl-4-(-methylpiperazin-1-yl)aniline 64 mg, 0.31 mmol), and methanesulfonic acid (32 mg, 0.34 mmol) in ACN (2 mL). The mixture was microwaved at 140 °C for 30 min to yield the double TFA salt form of the title compound as a mixture of rotamers that is dark brown amorphous solid (7.5 mg, 7.5 % yield). ^1^H NMR (400 MHz, DMSO-*d*_6_) δ 10.86 (s, 1H), 7.77 (d, *J* = 1.5 Hz, 1H), 7.66 (dd, *J* = 8.3, 1.5 Hz, 1H), 7.49 (d, *J* = 8.2 Hz, 1H), 7.37 (d, *J* = 8.7 Hz, 1H), 7.05 (d, *J* = 2.8 Hz, 1H), 6.98 (dd, *J* = 8.8, 2.8 Hz, 1H), 3.91 (d, *J* = 13.1 Hz, 2H), 3.55 (d, *J* = 11.9 Hz, 2H), 3.16 (s, 2H), 3.02 (d, *J* = 13.2 Hz, 2H), 2.88 (s, 3H), 2.54 (s, 5H), 2.23 (s, 3H).^13^C NMR (100 MHz, DMSO-*d*_6_) δ 158.66, 158.31, 151.52, 149.14, 135.44, 127.56, 126.03, 119.15, 118.07, 117.55, 115.22, 114.59, 112.55, 104.83, 52.28 (2C), 45.44 (2C), 42.10, 17.71. HPLC Purity > 99%. LCMS calculated for C_20_H_23_N_6_ [M + H]^+^: 347.2. Found: 347.3.

#### 2-((3-fluoro-4-morpholinophenyl)amino)-1H-benzo[d]imidazole-6-carbonitrile (9)

The reaction was carried out according to general procedure A with 2-chloro-1H-benzo[*d*]imidazole-5-carbonitrile (25 mg, 0.14 mmol), 3-fluoro-4-morphoaniline (30 mg, 0.15 mmol), and methanesulfonic acid (16 mg, 0.17 mmol) in ACN (2 mL). The mixture was microwaved at 140 °C for 30 min to yield the double TFA salt form with the title compound as a light brown amorphous solid (9.3 mg, 11.6% yield). ^1^H NMR (400 MHz, CD_3_OD) δ 7.73 (dd, *J* = 1.5, 0.7 Hz, 1H), 7.66 (dd, *J* = 8.3, 1.5 Hz, 1H), 7.53 (dd, *J* = 8.3, 0.6 Hz, 1H), 7.31 – 7.15 (m, 3H), 3.89 – 3.82 (m, 4H), 3.16 – 3.09 (m, 4H). ^13^C NMR (100 MHz, CD_3_OD) δ 156.84, 154.37, 150.72, 139.80, 139.72, 133.49, 130.29, 129.31, 129.21, 127.98, 120.72, 120.69, 119.76, 119.72, 118.11, 115.21, 112.79, 112.55, 112.32, 106.76, 66.51 (2C), 50.65, 50.62. HPLC Purity > 99%. LCMS calculated for C_18_H_17_FN_5_O [M + H]^+^: 338.1. Found: 338.4.

#### 2-((4-morpholino-3-(trifluoromethyl)phenyl)amino)-1H-benzo[d]imidazole-6-carbonitrile (10)

The reaction was carried out according to general procedure A with 2-chloro-1H-benzo[*d*]imidazole-5-carbonitrile (25 mg, 0.14 mmol), 4-morpholino-3-(trifluoromethyl)aniline (38 mg, 0.15 mmol), and methanesulfonic acid (16 mg, 0.17 mmol) in ACN (2 mL). The mixture was microwaved at 140 °C for 30 min to yield double TFA salt of the title compound as a purple amorphous solid (8.5 mg, 8.7 %)^1^H NMR (400 MHz, CD_3_OD) δ 7.82 (d, *J* = 2.6 Hz, 1H), 7.80 – 7.72 (m, 2H), 7.71 – 7.61 (m, 2H), 7.53 (dd, *J* = 8.3, 0.7 Hz, 1H), 3.88 – 3.78 (m, 4H), 3.00 – 2.94 (m, 4H). ^13^C NMR (214 MHz, CD_3_OD) δ 154.33, 154.29, 138.90, 136.08, 132.87, 130.63, 130.39, 127.62, 120.80, 120.58, 117.65, 117.12, 114.80, 109.35, 69.05 (2C), 51.61 (2C), 19.26. HPLC Purity > 95%. LCMS calculated for C_19_H_17_F_3_N_5_O [M + H]^+^: 388.1. Found: 388.1.

#### 2-((3,5-difluoro-4-morpholinophenyl)amino)-1H-benzo[d]imidazole-6-carbonitrile (11)

The reaction was carried out according to general procedure A with 2-chloro-1H-benzo[*d*]imidazole-5-carbonitrile (25 mg, 0.14 mmol), 3,5-difluoro-4-morphoaniline (30 mg, 0.15 mmol), and methanesulfonic acid (16 mg, 0.17 mmol) in ACN (2 mL). The mixture was microwaved at 140 °C for 30 min to yield the double TFA salt form of the title compound as a mixture of rotamers that is a light brown amorphous solid (10.2 mg, 15.5 % yield). ^1^H NMR (400 MHz, CD_3_OD) δ 7.75 (dd, *J* = 1.5, 0.7 Hz, 1H), 7.64 (dd, *J* = 8.3, 1.5 Hz, 1H), 7.55 (dd, *J* = 8.3, 0.6 Hz, 1H), 7.22 – 7.10 (m, 2H), 3.83 – 3.73 (m, 4H), 3.24 – 3.17 (m, 4H).^13^C NMR (100 MHz, CD_3_OD) δ 160.24, 160.15, 157.77, 157.68, 150.46, 134.39, 131.95, 131.82, 131.68, 131.17, 127.68, 126.38, 126.25, 118.29, 115.56, 112.60, 107.52, 107.43, 107.33, 107.24, 106.42, 67.16 (2C), 51.24, 51.21, 51.18. HPLC Purity > 99%. LCMS calculated for C_18_H_16_F_2_N_5_O [M + H]^+^: 356.1. Found: 356.1.

#### 2-((3-methoxy-4-morpholinophenyl)amino)-1H-benzo[d]imidazole-6-carbonitrile (12)

The reaction was carried out according to general procedure A with 2-chloro-1H-benzo[*d*]imidazole-5-carbonitrile (25 mg, 0.14 mmol), 3-methoxy-4-morphoaniline (32 mg, 0.15 mmol), and methanesulfonic acid (16 mg, 0.17 mmol) in ACN (2 mL). The mixture was microwaved at 140 °C for 30 min to yield the double TFA salt form of the title compound as a light purple amorphous solid (20.5 mg, 25.3% yield). ^1^H NMR (400 MHz, CD_3_OD) δ 7.74 (dd, *J* = 1.5, 0.6 Hz, 1H), 7.65 (dd, *J* = 8.3, 1.5 Hz, 1H), 7.53 (dd, J = 8.3, 0.7 Hz, 1H), 7.30 – 7.18 (m, 2H), 7.09 (dd, *J* = 8.5, 2.4 Hz, 1H), 3.95 (s, 3H), 3.94 – 3.89 (m, 4H), 3.27 – 3.20 (m, 4H). ^13^C NMR (126 MHz, CD_3_OD) δ 153.37, 150.95, 137.83, 134.14, 132.08, 130.92, 127.71, 119.51, 118.30, 116.10, 115.29, 112.37, 107.84, 106.40, 66.09, 55.12 (2C), 51.48 (2C). HPLC Purity >95%. LCMS calculated for C_19_H_20_N_5_O_2_ [M + H]^+^: 350.2. Found: 350.3.

#### 2-((2-methyl-4-morpholinophenyl)amino)-1H-benzo[d]imidazole-6-carbonitrile (13)

The reaction was carried out according to general procedure A with 2-chloro-1H-benzo[*d*]imidazole-5-carbonitrile (25 mg, 0.14 mmol), 2-methyl-4-morphoaniline (27 mg, 0.15 mmol), and methanesulfonic acid (16 mg, 0.17 mmol) in ACN (2 mL). The mixture was microwaved at 140 °C for 30 min to yield the double TFA salt form of the title compound as a light purple amorphous solid (14.6 mg, 18.4 % yield). ^1^H NMR (400 MHz, CD_3_OD) δ 7.70 (d, *J* = 1.5 Hz, 1H), 7.64 (dd, *J* = 8.4, 1.5 Hz, 1H), 7.50 (d, *J* = 8.3 Hz, 1H), 7.32 (d, *J* = 8.7 Hz, 1H), 7.06 (d, *J* = 2.9 Hz, 1H), 7.00 (dd, *J* = 8.7, 2.9 Hz, 1H), 3.90 – 3.83 (m, 4H), 3.28 – 3.21 (m, 4H), 2.31 (s, 3H). ^13^C NMR (100 MHz, CD_3_OD) δ 151.58, 151.51, 136.24, 133.38, 130.18, 127.94, 127.71, 125.06, 118.10, 117.97, 114.96, 114.51, 112.11, 106.66, 66.33 (2C), 48.99 (2C), 16.56. HPLC Purity >99% LCMS calculated for C_19_H_20_N_5_O [M + H]^+^: 334.2. Found: 334.1.

#### 2-((3-chloro-4-morpholinophenyl)amino)-1H-benzo[d]imidazole-6-carbonitrile (14)

The reaction was carried out according to general procedure A with 2-chloro-1H-benzo[*d*]imidazole-5-carbonitrile (25 mg, 0.14 mmol), 3-chloro-4-morphoaniline (33 mg, 0.15 mmol), and methanesulfonic acid (16 mg, 0.17 mmol) in ACN (2 mL). The mixture was microwaved at 140 °C for 30 min to yield the double TFA salt form of the title compound as a clear amorphous solid (3.6 mg, 4.4% yield). ^1^H NMR (400 MHz, CD_3_OD) δ 7.74 (dd, *J* = 1.5, 0.6 Hz, 1H), 7.67 (dd, *J* = 8.3, 1.5 Hz, 1H), 7.60 – 7.50 (m, 2H), 7.40 (dd, *J* = 8.6, 2.5 Hz, 1H), 7.30 (d, *J* = 8.6 Hz, 1H), 3.90 – 3.84 (m, 4H), 3.13 – 3.06 (m, 4H). ^13^C NMR (100 MHz, CD_3_OD) δ 150.66, 148.83, 133.41, 130.54, 130.23, 129.49, 128.02, 126.57, 123.92, 121.50, 118.10, 115.23, 112.34, 106.80, 66.67 (2C), 51.40 (2C). HPLC Purity > 99%. LCMS calculated for C_18_H_17_ClN_5_O [M + H]^+^: 354.2. Found: 354.3.

#### 2-((3-bromo-4-morpholinophenyl)amino)-1H-benzo[d]imidazole-6-carbonitrile (15)

The reaction was carried out according to general procedure A with 2-chloro-1H-benzo[*d*]imidazole-5-carbonitrile (25 mg, 0.14 mmol), 3-bromo-4-morphoaniline (40 mg, 0.15 mmol), and methanesulfonic acid (16 mg, 0.17 mmol) in ACN (2 mL). The mixture was microwaved at 140 °C for 30 min to yield the double TFA salt form of the title compound as a white powder (1.7 mg, 1.9% yield). ^1^H NMR (850 MHz, CD_3_OD) δ 7.76 (d, J = 2.5 Hz, 1H), 7.74 (dd, *J* = 1.5, 0.7 Hz, 1H), 7.66 (dd, *J* = 8.3, 1.5 Hz, 1H), 7.53 (dd, *J* = 8.3, 0.7 Hz, 1H), 7.46 (dd, *J* = 8.5, 2.6 Hz, 1H), 7.30 (d, *J* = 8.6 Hz, 1H), 3.89 – 3.86 (m, 4H), 3.10 – 3.07 (m, 4H). ^13^C NMR (214 MHz, CD_3_OD) δ 152.13, 151.54, 134.98, 132.52, 131.78, 131.02, 129.38, 125.86, 123.29, 121.47, 119.55, 116.67, 113.78, 108.15, 68.11 (2C), 53.24 (2C). HPLC Purity > 99%. LCMS calculated for C_18_H_17_BrN_5_O [M + H]^+^: 398.1. Found: 398.2.

#### 2-((6-morpholinopyridin-3-yl)amino)-1H-benzo[d]imidazole-5-carbonitrile (16)

The reaction was carried out according to general procedure A with 2-chloro-1H-benzo[*d*]imidazole-5-carbonitrile (50 mg, 0.28 mmol), 6-morpholinopyridin-3-amine (56 mg, 0.31 mmol), and methanesulfonic acid (32 mg, 0.34 mmol) in ACN (4 mL). The mixture was microwaved at 140 °C for 30 min to yield the double TFA salt form of the title compound as a dark purple amorphous solid (15 mg, 9.7 % yield). ^1^H NMR (500 MHz, CD_3_OD) δ 8.37 (d, *J* = 2.7 Hz, 1H), 7.79 (dd, *J* = 9.2, 2.7 Hz, 1H), 7.71 (d, *J* = 1.4 Hz, 1H), 7.61 (dd, *J* = 8.2, 1.4 Hz, 1H), 7.50 (d, *J* = 8.3 Hz, 1H), 7.09 (d, *J* = 9.2 Hz, 1H), 3.85 – 3.80 (m, 4H), 3.60 (dd, *J* = 5.7, 4.1 Hz, 4H). ^13^C NMR (126 MHz, CD_3_OD) δ 156.87, 151.86, 140.54, 136.17, 134.95, 131.72, 127.43, 123.36, 118.45, 115.42, 112.47, 109.10, 106.04, 66.08 (2C), 45.50 (2C). HPLC Purity > 99%. LCMS calculated for C_17_H_17_N_6_O [M + H]^+^: 321.1. Found: 321.2.

#### 2-((3-(hydroxymethyl)-4-(4-methylpiperazin-1-yl)phenyl)amino)-1H-benzo[d]imidazole-6-carbonitrile (17)

The reaction was carried out according to general procedure A with 2-chloro-1H-benzo[*d*]imidazole-5-carbonitrile (25 mg, 0.14 mmol), (5-amino-2-(4-methylpiperazin-1-yl)phenyl)methanol (34 mg, 0.15 mmol), and methanesulfonic acid (16 mg, 0.17 mmol) in ACN (2 mL). The mixture was microwaved at 140 °C for 30 min to yield the double TFA salt form of the title compound as a clear amorphous solid (2.5 mg, 3.0% yield). ^1^H NMR (500 MHz, CD_3_OD) δ 7.74 (d, *J* = 1.4 Hz, 1H), 7.66 (dd, *J* = 8.3, 1.5 Hz, 1H), 7.61 (d, *J* = 2.7 Hz, 1H), 7.54 (d, *J* = 8.3 Hz, 1H), 7.41 (dd, *J* = 8.5, 2.7 Hz, 1H), 7.35 (d, *J* = 8.5 Hz, 1H), 4.78 (s, 2H), 3.62 (d, *J* = 11.8 Hz, 2H), 3.36 (d, *J* = 11.1 Hz, 4H), 3.18 (t, *J* = 13.2 Hz, 2H), 3.00 (s, 3H). ^13^C NMR (126 MHz, CD_3_OD) δ 152.10, 149.71, 139.99, 134.97, 133.31, 131.76, 129.35, 125.83, 125.09, 122.80, 119.58, 116.61, 113.74, 108.09, 60.55 (2C), 55.26, 51.01 (2C), 43.65. HPLC Purity > 99% LCMS calculated for C_20_H_23_N_6_O [M + H]^+^: 363.2. Found: 363.0.

#### 2-((3-chloro-4-(trifluoromethyl)phenyl)amino)-1H-benzo[d]imidazole-5-carbonitrile (18)

The reaction was carried out according to general procedure A with 2-chloro-1H-benzo[*d*]imidazole-5-carbonitrile (25 mg, 0.14 mmol), 3-chloro-4-(trifluoromethoxy)aniline (30 mg, 0.14 mmol), and methanesulfonic acid (16 mg, 0.17 mmol) in ACN (2 mL). The mixture was microwaved at 140 °C for 30 min to yield the TFA salt form of the title compound as a clear amorphous solid (11.8 mg, 24% yield). ^1^H NMR (500 MHz, DMSO-*d_6_*) δ 10.29 (s, 1H), 8.24 (d, *J* = 2.6 Hz, 1H), 7.79 (s, 1H), 7.71 (dd, *J* = 9.0, 2.7 Hz, 1H), 7.55 (dd, *J* = 9.0, 1.4 Hz, 1H), 7.50 (d, *J* = 8.2 Hz, 1H), 7.46 (dd, *J* = 8.1, 1.5 Hz, 1H). ^13^C NMR (126 MHz, DMSO-*d_6_*) δ 140.56, 137.54, 126.29, 124.78, 123.83, 121.26, 120.46, 119.22, 118.42, 117.32. HPLC Purity > 99%. LCMS calculated for C_15_H_9_ClF_3_N_4_O [M + H]^+^: 353.1. Found: 353.0.

#### 2-((3-chloro-4-(difluoromethyl)phenyl)amino)-1H-benzo[d]imidazole-5-carbonitrile (19)

The reaction was carried out according to general procedure A with 2-chloro-1H-benzo[*d*]imidazole-5-carbonitrile (25 mg, 0.14 mmol), 3-chloro-4-(difluoromethoxy)aniline (27 mg, 0.14 mmol), and methanesulfonic acid (16 mg, 0.17 mmol) in ACN (2 mL). The mixture was microwaved at 140 °C for 30 min to yield the TFA salt form of the title compound as a white powder (19.3 mg, 18% yield). ^1^H NMR (500 MHz, DMSO-*d_6_*) δ 11.74 (s, 1H), 10.16 (s, 1H), 8.21 (s, 1H), 7.80 (d, *J* = 2.7 Hz, 1H), 7.65 (d, *J* = 11.6 Hz, 1H), 7.48 (s, 1H), 7.43 (d, *J* = 9.8 Hz, 1H), 7.36 (d, *J* = 9.0 Hz, 1H), 7.17 (t, *J* = 73.6 Hz, 1H). ^13^C NMR (126 MHz, DMSO-*d_6_*) δ 165.03, 140.69, 139.25, 125.59, 122.65, 120.96, 119.23, 119.06, 117.80, 117.15, 115.09. HPLC Purity > 99%. LCMS calculated for C_15_H_10_ClF_2_N_4_O [M + H]^+^: 335.0. Found: 335.0.

#### 2-((3-chloro-4-(piperidin-1-yl)phenyl)amino)-1H-benzo[d]imidazole-5-carbonitrile (20)

The reaction was carried out according to general procedure A with 2-chloro-1H-benzo[*d*]imidazole-5-carbonitrile (25 mg, 0.14 mmol), 3-chloro-4-(piperidin-1-yl)aniline (33 mg, 0.15 mmol), and methanesulfonic acid (16 mg, 0.17 mmol) in ACN (2 mL). The mixture was microwaved at 140 °C for 30 min to yield the formic acid salt form of the title compound as a clear amorphous solid (1.1 mg, 1.7% yield). ^1^H NMR (400 MHz, CD_3_OD) δ 7.74 (dd, *J* = 1.5, 0.6 Hz, 1H), 7.66 (dd, *J* = 8.3, 1.5 Hz, 1H), 7.59 (d, *J* = 2.5 Hz, 1H), 7.53 (dd, *J* = 8.3, 0.7 Hz, 1H), 7.39 (dd, *J* = 8.6, 2.5 Hz, 1H), 7.32 (d, *J* = 8.7 Hz, 1H), 3.13 – 3.06 (m, 4H), 1.81 (dq, *J* = 11.0, 5.2 Hz, 4H), 1.70 – 1.60 (m, 2H). ^13^C NMR (214 MHz, CD_3_OD) δ 155.19, 147.48, 136.51, 130.57, 126.28, 122.86, 122.25, 121.35, 120.23, 54.35 (2C), 27.37 (2C), 25.30. HPLC Purity > 99%. LCMS calculated for C_19_H_19_ClN_5_ [M + H]^+^: 352.1. Found: 353.3.

#### 2-((3-chloro-4-(4-methylpiperazin-1-yl)phenyl)amino)-1H-benzo[d]imidazole-6-carbonitrile (21)

The reaction was carried out according to general procedure A with 2-chloro-1H-benzo[d]imidazole-5-carbonitrile (25 mg, 0.14 mmol), 3-chloro-4-(4-methylpiperazin-1-yl) aniline (35 mg, 0.15 mmol), and methanesulfonic acid (16 mg, 0.17 mmol) in ACN (2 mL). The mixture was microwaved at 140 °C for 30 min to yield the double TFA salt form of the title compound as a yellow amorphous solid (9.1 mg, 11% yield). ^1^H NMR (400 MHz, CD_3_OD) δ 7.78 – 7.73 (m, 1H), 7.70 – 7.61 (m, 2H), 7.55 (dd, *J* = 8.3, 0.7 Hz, 1H), 7.44 (dd, *J* = 8.6, 2.5 Hz, 1H), 7.36 (d, *J* = 8.6 Hz, 1H), 3.66 (d, *J* = 12.0 Hz, 2H), 3.58 (d, *J* = 13.3 Hz, 2H), 3.38 (t, *J* = 11.9 Hz, 2H), 3.18 (t, *J* = 11.7 Hz, 2H), 3.00 (s, 3H). ^13^C NMR (100 MHz, CD_3_OD) δ 150.61, 146.66, 133.69, 132.02, 130.49, 129.80, 127.96, 126.17, 123.63, 122.00, 118.18, 115.36, 112.45, 106.73, 53.64 (2C), 42.27 (2C). HPLC Purity > 97%. LCMS calculated for C_19_H_20_ClN_6_ [M + H]^+^: 367.1. Found: 367.4.

#### 2-((3-chloro-4-(piperazin-1-yl)phenyl)amino)-1H-benzo[d]imidazole-5-carbonitrile (22)

The reaction was carried out according to general procedure B with 2-chloro-1H-benzo[*d*]imidazole-5-carbonitrile (200 mg, 1.13 mmol), tert-butyl 4-(4-amino-2-chlorophenyl)piperazine-1-carboxylate (386 mg, 1.24 mmol), and potassium dihydrogen phosphate (153 mg, 1.13 mmol) in 1-butanol (4 mL). After concentrating the crude reaction mixture, dichloromethane (3 mL) was added with trifluoracetic acid (642 mg, 5.63 mmol) to yield the double TFA salt form of the title compound as a clear amorphous solid (66.3 mg, 16.7% yield over two steps). ^1^H NMR (500 MHz, CD_3_OD) δ 7.80 (d, *J* = 2.6 Hz, 1H), 7.63 (d, *J* = 1.3 Hz, 1H), 7.49 (dd, *J* = 8.7, 2.6 Hz, 1H), 7.45 – 7.37 (m, 2H), 7.21 (d, *J* = 8.7 Hz, 1H), 3.43 – 3.37 (m, 4H), 3.28 – 3.23 (m, 4H). ^13^C NMR (126 MHz, CD_3_OD) δ 150.39, 147.51, 133.32, 131.62, 130.15, 129.85, 128.17, 126.59, 124.08, 122.09, 118.07, 115.33, 112.44, 107.01, 43.75 (2C), 43.69 (2C). HPLC Purity > 99%. LCMS calculated for C_18_H_18_ClN_6_ [M + H]^+^: 353.1. Found: 353.0.

#### N-(3-(4-(2-chloro-4-((5-cyano-1H-benzo[d]imidazol-2-yl)amino)phenyl)piperazin-1-yl)-3-oxopropyl)-3-(5,5-difluoro-7-(1H-pyrrol-2-yl)-5H-5λ4,6λ4-dipyrrolo[1,2-c:2’,1’-f][1,3,2]diazaborinin-3-yl)propenamide (23)

A reaction flask with 3-((*tert*-butoxycarbonyl)amino)propanoic acid (32 mg, 0.17 mmol), HATU (65 mg, 0.17 mmol), and DIPEA (66 mg, 0.51 mmol) dissolved in DMF (2.0 mL) was allowed to stir at r.t.. After 15 minutes of stirring, 2-((3-chloro-4-(piperazin-1-yl)phenyl)amino)-1*H*-benzo[*d*]imidazole-5-carbonitrile (**22**) (60 mg, 0.17 mmol) was added, and the reaction was allowed to stir overnight. The reaction mixture was diluted with H_2_O and extracted with EtOAc, and the combined organic layers were washed with brine and then dried over Na_2_SO_4_. The dried organic layer was concentrated *in vacuo*. 50 mg of the crude reaction product *tert*-butyl (3-(4-(2-chloro-4-((5-cyano-1*H*-benzo[*d*]imidazol-2-yl)amino)phenyl)piperazin-1-yl)-3-oxopropyl)carbamate was added to a vial with DCM (1 mL) and TFA (296 mg, 2.6 mmol). The reaction was stirred for 2 h and dried *in vacuo*. A portion of the dried crude reaction containing 2-((4-(4-(3-aminopropanoyl)piperazin-1-yl)-3-chlorophenyl)amino)-1*H*-benzo[*d*]imidazole-5-carbonitrile was added to a vial containing *N*,*N*-Diisopropylethylamine (15 mg, 0.11 mmol), 2,5-dioxopyrrolidin-1-yl 3-(5,5-difluoro-7-(1*H*-pyrrol-2-yl)-5*H*-5λ^4^,6λ^4^-dipyrrolo[1,2-*c*:2’,1’-*f*][1,3,2]diazaborinin-3-yl)propanoate (5 mg, 12 μmol), and DMF (1 mL). The reaction was stirred o.n., concentrated *in vacuo*, and purified using preparative HPLC (10–100% MeOH in H_2_O + 0.05%TFA) to yield the double TFA form of the pure final product as a purple amorphous solid (3.2 mg, 6.7 % yield over three steps). ^1^H NMR (500 MHz, CD_3_OD) δ 7.67 (d, *J* = 1.5 Hz, 1H), 7.63 (d, *J* = 2.6 Hz, 1H), 7.54 (dd, *J* = 8.4, 1.5 Hz, 1H), 7.47 (d, *J* = 8.3 Hz, 1H), 7.34 (dd, *J* = 8.6, 2.6 Hz, 1H), 7.25 – 7.20 (m, 2H), 7.20 – 7.14 (m, 3H), 6.99 (d, *J* = 4.6 Hz, 1H), 6.93 (d, *J* = 4.0 Hz, 1H), 6.37 – 6.31 (m, 2H), 3.77 – 3.70 (m, 2H), 3.69 – 3.64 (m, 2H), 3.49 (t, *J* = 6.7 Hz, 2H), 3.29 (d, *J* = 7.4 Hz, 2H), 3.00 (dt, *J* = 10.1, 4.9 Hz, 4H), 2.64 (q, *J* = 6.9 Hz, 4H). HPLC purity > 99%. LCMS calculated for C_37_H_35_BClF_2_N_10_O_2_ [M + H]^+^: 735.3. Found: 735.1.

#### 2-((4-(2-hydroxypropan-2-yl)phenyl)amino)-1H-benzo[d]imidazole-5-carbonitrile (Compound 1h)

To a vial containing methyl 4-((6-cyano-1H-benzo[*d*]imidazol-2-yl)amino)benzoate (**1**) (136 mg, 465 μmol) in THF (5 mL), stirred at -78°C, methyl lithium in ether (1.6 M, 1.45 mL, 2.33 mmol) was added dropwise. After 1.5 h, the reaction was allowed to warm to r.t. The reaction mixture was diluted with H_2_O and then extracted with EtOAc, and the combined organic layers were washed with brine and then dried over Na_2_SO_4_. The dried organic layers were then concentrated *in vacuo* and purified using preparative HPLC (10–100% MeOH in H_2_O + 0.05% NH_4_OH) to yield the pure final product as a clear amorphous solid (13.2 mg, 9.7 % yield). ^1^H NMR (500 MHz, CD_3_OD) δ 7.59 (s, 1H), 7.54 – 7.44 (m, 4H), 7.38 (d, *J* = 1.2 Hz, 2H), 1.55 (s, 6H). HPLC Purity > 99%. LCMS calculated for C_17_H_16_N_4_O [M + H]^+^: 293.2. Found: 293.4.

### Biological Evaluation

#### NaLTSA

All NaLTSA assays were performed using an altered version of existing published protocols^27^. A transfection complex was formed with 9 μg/mL carrier DNA and 1 μg/mL of a DNA construct containing either CSNK1G1-NL, CSNK1G2-NL, or CSNK1G3-NL in Opti-MEM without phenol red (Gibco). Once the DNA solution was made, for every mL of DNA solution, 30 μL of FuGENE HD was added. The transfection solution was immediately vortexed and incubated at r.t. for at least 20 minutes. The transfection complex solution was mixed with a 20x volume of HEK293 cells in DMEM supplemented with 10% FBS for a final concentration of 2x10^5^ cells/mL. The transfected cell solution was then plated onto a T75 plate and incubated at 37°C in 5% CO_2_. After a 24-hour incubation, the cells were washed with PBS and harvested with Trypsin (GIBCO). The harvested cells were resuspended in DMEM supplemented with 10% FBS and added to a 15 mL centrifuge tube (Falcon). The resuspended cells were then centrifuged at 1200 rpm for 5 minutes. The trypsin-containing media was aspirated, and the pelleted cells were resuspended and then diluted to a concentration of 225,000 cells/mL in Opti-MEM.

To prepare the compound plate, a protease inhibitor cocktail (reconstituted in DMSO) was used to make a stock solution with compound **14** at a 100x concentration. One μL of the 100x protease inhibitor/compound **14** solution was added to all 96 wells in a 3 × 32-well PCR Reaction Plate (Thermo Fisher). A digitonin solution was then prepared at 500 μg/mL (10x) in Opti-MEM, and 10 μL of this solution was added to each well of the PCR reaction plate containing the protease inhibitor/compound **14** solution. After digitonin addition, 89 μL of the resuspended cells in Opti-MEM was added to every well to bring the final volume to 100 μL per well. The solution was then pipetted up and down to allow for thorough mixing. Each section of the 3x32 well PCR compound plates was then covered with an adhesive seal (Thermo Fisher) and incubated at r.t. for 40 min. After the r.t. incubation, the cells were then incubated at a temperature range from 40-73 °C with 3-degree increments for 3 min in a thermal cycler (Applied Biosystems ProFlex PCR System, Thermo Fisher). The cells were then removed from the thermal cycler, cooled at r.t. for three minutes, and transferred to a white non-binding surface (NBS) 96-well plate (Corning, 3917). 25 μL of a 5x solution of NanoGlo substrate (Promega) was added to each well, and the plates’ total luminescence was then read using a GloMax Discover luminometer (Promega). To generate melting temperature values, the raw luminescence data were normalized to the 40 °C luminescence values. The resulting values, indicated as percent stabilized, were then fitted to gather apparent melting temperature values using the Boltzmann Sigmoid Equation. All NaLTSA experiments were run in technical duplicates and repeated independently three times.

#### General information for NanoBRET Assays

All NanoBRET assays were performed with a modified version of previously published protocols^28,34,48,49^. Human Embryonic Kidney (HEK293) cells from ATCC were cultured at 37 °C in 5% CO_2_ in Dulbecco’s modified Eagle medium (DMEM; Gibco) supplemented with 10% fetal bovine serum (FBS; Avantor). To form the transfection complex of DNA at 10 μg/mL, carrier DNA (Promega) was added at 9 μg/mL and 1 μg/mL NLuc fusion protein (NL-CSNK1G1, NL-CSNK1G2, NL-CSNK1G3, NL-GSK3β, NL-CSNK1D, NL-CSNK1E, NL-MOK, or CLK4-NL) in Opti-MEM without serum or phenol red (Gibco) with FuGENE HD (Promega) added at 30 μL/mL. The transfection complex solution was then vortexed and incubated at r.t. for a period of at least 20 minutes. The transfection complex solution was mixed with a 20x volume of HEK293 cells in DMEM supplemented with 10% FBS to allow for a final concentration of 2x10^5^ cells/mL. 100 μL of the resulting solution was then added to a 96-well tissue culture-treated plate (Corning, 3917).

The following day, the media was aspirated and replaced with 90 μL of Opti-MEM in no tracer wells, 85 μL in wells with tracer, and 75 μL in wells with tracer and digitonin. A total of 5 μL per well of a 20x tracer dilution solution, made with tracer dilution buffer, was added to each well except the “no tracer” control wells. Either 10 μL of 10x compound stock solution made up in Opti-MEM or, for control wells, 10 μL of Opti-MEM and DMSO was added to every control well for a final concentration of DMSO of 1.1% in all wells. For permeabilized cell analysis, 10 μL of a 10x digitonin made in Opti-MEM was added to all wells (50 μg/mL). For intact cell analysis, the NanoBRET substrate solution was made using the NanoBRET NanoGlo substrate (Promega) at a ratio of 1:166 in Opti-MEM with an extracellular NLuc inhibitor (Promega) diluted at 1:500. After incubating the 96-well plate with tracer and test compounds for 2 hours at 37 °C in 5% CO_2_, 50 μL of the NanoBRET substrate solution was added to each well. For permeabilized cell analysis, an adjusted NanoBRET substrate solution was made using NanoBRET NanoGlo substrate at a ratio of 1:166 in Opti-MEM with no extracellular NLuc inhibitor. After the plate was incubated at r.t. for no longer than 25 min after digitonin addition, 50 μL of the adjusted NanoBRET substrate solution was added to each well. After substrate addition, the 96-well plates were then read within 10 minutes of substrate addition with a GloMax Discover luminometer (Promega) using a 450 nm BP filter (donor) and a 600 nm LP filter (acceptor) with an integration time of 0.3 seconds. To analyze the data, Raw milliBRET units (mBU) values were generated by dividing the acceptor emission (600 nm) by the donor emissions (450 nm) and multiplying the resulting values by 1000. All NanoBRET studies’ mBU values were background corrected by subtracting the average of the “no tracer” control wells.

#### Tracer EC_50_ Determination

HEK293 were transfected with NL-CSNK1G1, NL-CSNK1G2, or NL-CSNK1G3. **23** was tested in an 11-point dose-response format with a top concentration of 1.5 μM (intact) or 1 μM (permeabilized). Three biological replicate(s) were plotted using GraphPad Prism software and fit using a Sigmoidal three-parameter dose-response logistical curve to determine EC_50_ values. Error bars indicate the standard deviation.

#### NanoBRET Tracer titration

HEK293 cells were transfected with NL-CSNK1G1, NL-CSNK1G2, or NL-CSNK1G3. 20x stocks of tracer **23** were prepared in tracer dilution buffer containing 20% DMSO, such that the final concentrations were 100, 200, 300, 400, and 500 nM after adding 5 μL of the tracer stocks to the 96-well plate. Compound **14** was tested for each tracer concentration in an 11-point dose-response format with a top concentration of 10 μM by adding 10 μL of a 10x dilution series of Compound **14** prepared in Opti-MEM. 3 biological replicate(s) were plotted using GraphPad Prism software and fit using a Sigmoidal three-parameter dose-response logistical curve to determine IC_50_ values. Error bars indicate the standard deviation.

#### NanoBRET Inhibitor Screening Assays

HEK293 cells were transfected with NL-CSNK1G1, NL-CSNK1G2, NL-CSNK1G3, NL-GSK3β, NL-CSNK1D, NL-CSNK1E, NL-MOK or CLK4-NL. Based on tracer titration results, inhibitor screening assays were conducted using 250 nM of compound **14** for CSNK1G2, 500 nM of compound **14** for CSNK1G1 and CSNK1G3. Based on the manufacturer’s recommendations, assays were performed with tracer K8 (Promega) at 130 nM for CSNK1D, 500 nM for CSNK1E, 63 nM for GSK3B, and 170 nM of tracer K9 was used for CLK4. Each compound was tested in an 11-point dose-response format with a starting concentration of either 10 μM or 30 μM by adding 10 μL of a 10x dilution series of compound solution prepared in Opti-MEM without phenol red. NanoBRET data was plotted using GraphPad Prism software and fit using a Sigmoidal three-parameter dose-response logistical curve to determine IC_50_ values. Error bars indicate the standard deviation.

#### K192 Screening

The K192 kinome screen was executed with a modified version of previously published protocols^28,48,50^. For all experiments, HEK293 cells were cultured in DMEM with 10% FBS at 37 °C in 5% CO_2_. DNA solutions were made through the reconstitution of the NanoBRET TE K192 plates. To create the transfection solution, 10 μL of the 10x DNA solutions were aliquoted with 30 μL of a Fugene solution (30 μL/mL in Opti-MEM) in 96-well plates (Corning, 3917). The transfection lipid: DNA complexes were allowed to form during a 20-minute incubation at r.t. For the negative control, the pNL1.1.CMV [NLuc/CMV] Vector (Promega) was used. HEK293 cells were grown to 70-90% confluency in DMEM with 10% FBS at 37 °C in 5% CO_2_. The cells were harvested and reconstituted in DMEM with 10% FBS at 2.5x10^5^ cells per mL. To the transfection solution, 60 μL of the cell suspension was added to the 40 μL of transfection solution in the 96-well plates. The plates containing cells and lipid: DNA complexes were incubated overnight at 37 °C in 5% CO_2_.

The following day, the DMEM was aspirated and replenished with 85 μL of Opti-MEM. 5 μL of a 20x K10 tracer (Promega) solution was prepared and added at the manufacturer’s recommended concentrations. A 10x solution of compound **14** or DMSO vehicle was then prepared by diluting 4 µL of a 10 mM stock of compound **14** or DMSO, respectively, into Opti-MEM. For the top control wells, 10 µL of the DMSO vehicle solution was added, and for the sample wells, 10 µL of the compound **14** solution was added. The plates, now containing tracer and compound/vehicle, were then shaken at 300 rpm and incubated for 2 h at 37 °C in 5% CO_2_. After incubating for 2 h, the plates were cooled by a 15 minute incubation at r.t. Then, a 3x NLuc substrate solution was prepared using NanoBRET NanoGlo substrate (Promega) and the extracellular NLuc inhibitor (Promega) at the manufacturer’s recommended volumes in Opti-MEM without phenol red. To each well, 50 µL of the 3x NLuc substrate solution was added. All plates were shaken at 300 rpm on an orbital shaker for 15 seconds. Within 15 minutes of substrate additions, the plates were analyzed using a GloMax Discover luminometer (Promega) equipped with a 450 nm BP filter (donor) and a 600 nm LP filter (acceptor) with an integration time of 0.3 seconds. The raw BRET values were then calculated by dividing the acceptor emission values (600 nm) by the donor emission values (450 nm). To determine the fractional occupancy, the following equation was used:

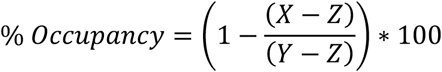

X = The mean BRET value across all wells containing the tracer and the test compound (sample wells) for a single kinase.

Y = The mean BRET value for all wells containing the tracer and the vehicle (top control wells) for a single kinase.

Z = Mean BRET value of the NLuc control wells (negative control wells)

#### Multiplexed Inhibitor Bead Assay

The MIBs screen was executed using a modified version of previously published protocols ^39, 40^. A protein lysate solution, derived from HEK293 cells, containing ≥ 5 mg total protein in 4 mL of lysis buffer (50 mM HEPES buffer, 150 mM NaCl, 0.5 % Triton X-100, 1 mM EDTA, 1 mM EGTA, 10 mM NaF, 2.5 mM NaVO_4_, 2 tablets per 50 mL of Complete Protease Inhibitor Cocktail (Roche), 1% Phosphatase Inhibitor Cocktail 2 (Sigma), and 1% Phosphatase Inhibitor Cocktail 3 (Sigma)) was mixed with 4 µL of DMSO or a compound **14** stock solution followed by a 1 hour incubation on ice. For each sample, a Bio-Rad Poly-Prep Column was prepared with 350 µL of a 50% slurry of a total MIB matrix mix consisting of the following inhibitors Shokat, Purvalanol B, PP58, UNC-21474 (14% each by volume), VI-16832, and Ctx-0294885 (22% each by volume) attached to ECH Sepharose 4B in 20% aq. ethanol. To each column, 2 mL of a high salt buffer was added (50 mM HEPES, 0.5% Triton X-100, 1 M NaCl, 1 mM EDTA, 1 mM EGTA, pH 7.5) for equilibration. The cell lysate samples were then brought to a concentration of 1 M NaCl. The cell lysates were then added to the columns containing the MIB slurry. After the cell lysate flow through had passed, columns were washed 5 mL of the high salt MIB wash buffer, 5 mL of the low salt MIB wash buffer (50 mM HEPES, 150 mM NaCl, 0.5% Triton X-100, 1 mM EGTA, 1 mM EDTA, and 500 pH 7.5), and 500 µL of low salt MIB wash buffer with 0.1% SDS. To elute, the columns were capped, and the MIB slurry was resuspended with 500 µL of MIB elution buffer (0.1 M Tris-HCl, 0.5% w/v SDS, pH 7.25). The mixture was then transferred to an Eppendorf tube and heated at 95°C for 10 minutes. This step was then repeated. The elution buffer, now containing the proteins that the MIBs beads pulled down, was then isolated from the MIBs slurry. The proteins in the elution solution were then reduced using 5 mM DL-Dithiothreitol while the solution was incubated at 60 °C on a heating block for 25 min. To alkylate all free cysteines, 19 mM iodoacetamide was added, and the solution was incubated in a dark chamber at r.t. for 30 min. After the incubation, to quench the alkylation reaction, additional DL-Dithiotheitol was added. To achieve a final concentration of 10 mM protein, the elution solution was concentrated using Amicon Millipore Ultra-4 10K cutoff spin columns. The samples were centrifuged with the 10 kDa filter for 30 min at 3000 rpm, 4 °C. The concentrated samples were then subjected to a methanol/chloroform extraction. For the extraction, 100 µL of chloroform, 300 µL LC-MS grade water, and 400 µL of methanol was pipetted into each sample. Samples were vortexed thoroughly and then centrifuged at 1500 rpm for 10 minutes at 4 °C. The aqueous layer was removed and samples were washed four times with methanol (500 µL). All samples were then dried using a Labconco Acid-Resistant CentriVap Concentrator for 30 minutes. The samples were then reconstituted in a 100 µL solution of LC-MS grade water containing 50 mM HEPES, pH 8.0 buffer. To digest the sample, 3 µL of 0.4 µg/µL sequencing-grade Modified Trypsin (Promega) was added to all samples. The samples were then mixed and incubated at 37°C overnight for digestion.

The digested samples were washed with water-saturated ethyl acetate, and the organic layer was discarded after the samples were centrifuged at 15000 rpm for 5 minutes. This step was repeated three times. The washed samples were dried using a Labconco Acid-Resistant CentriVap Concentrator for 2 hours. The trypsin-digested dried samples were then resuspended in 200 μL of equilibration buffer (5% ACN and 0.5% TFA in LC-MS grade water) and loaded onto pre-equilibrated C-18 PepClean Spin Columns (Pierce). 200 µL equilibration buffer was then added to the columns, which were centrifuged for 1.5 min at 4400 rpm. The wash step was repeated. The samples were then eluted from the column by running 50 µL of 50% ACN in LC-MS grade water over the column and centrifuging each column at 4000 rpm for 1 minute.

The LC-MS/MS analysis was performed by the UNC metabolomics and proteomics core using the following previously published protocol. The dried tryptic peptides were resuspended in 15 μL 2% ACN, 0.1% Formic Acid to prepare the samples for LC-MS/MS analysis. A Thermo Scientific Easy nLC 1200 coupled to a Thermo Scientific Biopharma QExactive HF Orbitrap Mass Spectrometer equipped with an Easy-Spray Nano Source was used for the LC-MS/MS analysis. The dried tryptic peptides were separated using an Easy-Spray PepMap C18 column (75 µm ID X 25cm, 2 µm particle size; Thermo Scientific) and they were then eluted using a 90 min method. Separation of peptides was achieved with a gradient of 5– 45% mobile phase B (80% ACN, 0.1% formic acid, v/v) at a 250 nL/min flow rate with mobile phase A (water with 0.1% formic acid, v/v). The 15 most intense precursors were picked for further fragmentation by using QExactive HF operated in data-dependent mode. The resolution for the precursor scan (m/z 375– 1700) was set to 60,000 with an AGC target of 3×106 ions, 100 ms max IT. The normalized collision energy was set to 27% for HCD. MS/MS scans resolution was set to 15,000 with an AGC target value of 1×105 ions, 100 ms max IT. Dynamic exclusion was set to 30 sec, and precursors with an unknown charge or a charge state of 1 and ≥ 7 were excluded. MaxQuant (version 1.6.15.0) was then used to screen the data against a reviewed human database and a contaminants database using Andromeda within MaxQuant. Enzyme specificity was set to trypsin, allowing for two missed cleavages. All human kinase sequences were imported into Perseus (version 1.6.14.0) for additional processing. Decoy proteins, contaminants, single hits, and proteins within Perseus with 50% missing values were removed from the analysis. Then, the Log_2_ transformation of LFQ intensity was performed. The Log_2_ fold change (FC) ratios were calculated using the Log_2_ LFQ intensities of the compound **14** treated sample compared to the DMSO control.

#### Cell-Titer Glo HEK293 cell viability assay

Cell-Titer Glo viability assays were performed using an altered version of existing published protocols^51^. HEK293 cells, cultured in DMEM with 10% FBS at 37 °C in 5% CO_2_, were plated at 12,000 cells/well in a 96-well plate (Corning) and incubated overnight (37 °C, 5% CO_2_). Compound **14** was added to wells in a seven-point dose response in quadruplicate and incubated for either 24 or 48 hours. DMSO percentage was constant across each compound concentration. 70 μL of CellTiter-Glo 2.0 (Promega) was added to every well. The plate was then shaken for 2 min at 300 rpm and incubated for 10 min at r.t. The luminescence signal was then read on a GloMax plate reader (Promega). The analysis of the dose-response was done on GraphPad Prism. Cell viability values were calculated through the following equation:

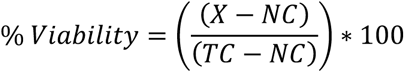

X = The mean RLU value for a concentration

NC = The mean RLU value of the 10% DMSO negative control

TC = The mean RLU value of the 1% DMSO top control

#### Eurofins Enzymatic Assay

Enzymatic radiometric enzyme assays performed by Eurofins were performed using the ATP values at the K_m_ value for each kinase. A 9-point dose-response curve was generated to gather IC50 values for each kinase tested. Further details on the assay procedure, technical controls, protein constructs, or dosing are available through Eurofins.

#### Dynamic Light Scattering (DLS)

The solubility of compound SGC-CK1γ-1 was estimated using dynamic light scattering (DLS), which measures the scattering intensity associated with compound aggregation in solution. SGC-CK1γ-1 was serially diluted from DMSO stock solutions (up to 100 µM final concentration) and then further diluted 50-fold into HBS buffer (final DMSO concentration 2%) for DLS analysis, following the protocol described by Aleandri et al.^52^ Both scattering intensity (in counts per second, Cnt/s) and the instrument-adjusted laser power (%) were monitored as indirect indicators of compound solubility. Scattering intensities above 1x10^6^ were considered to indicate the presence of aggregated compounds. All measurements were performed in triplicate at a fixed temperature of 25 °C.

#### Immunoblotting

Standard Western blot techniques were performed as previously described protocols^1^,^52^. HEK293 cells were seeded at 2x10^5^ cells/mL in 6-well plates in DMEM supplemented with 10% FBS. After a 24-h incubation (37 °C, 5% CO_2_), the cells were treated with serum-free DMEM without phenol red containing test compound or DMSO vehicle and incubated at 37 °C in 5% CO_2_ for 2 h. After the incubation, the cells were harvested and lysed on ice using RIPA buffer containing protease and phosphatase inhibitor (Thermo Scientific) and benzonase inhibitor (4 ug/mL). After the cells had incubated on ice for 10 minutes in the RIPA buffer, each sample was sonicated. Following sonication, cell lysates were centrifuged for 10 min at 14,000 rpm at 4 °C. Protein gels were performed with 30 µg of protein per lane in Novex Tris-glycine 4-12% gels. The membrane transfer was performed on the iBlot3 onto a polyvinylidene fluoride (PVDF) membrane. Membranes were blocked and then incubated overnight with primary antibody at r.t. with rotation. After overnight exposure to antibody, the blots were washed with Tris-buffered saline with Tween-20 (TBST), incubated for 1h at r.t with secondary antibody, washed with TBST, and imaged after exposure to SuperSignal West Femto reagent (Thermo Fisher) for chemiluminescence. All primary antibodies used are as follows: LRP6 (1:1000; Cell Signaling Technology; catalog no.: 3395), LRP6 T1479 (1:500; Life Technologies; catalog no.: PA564736), and actin (1:5000; Sigma, catalog no.: A2228). All secondary antibodies were used as follows: Rb-HRP (1:5,000) and IR Dye 680RD (1:10,000, LiCOR, #926-68070).

#### Human cytomegalovirus infection of HFF cells

Human foreskin fibroblast (HFF) cells (clone Hs27) were obtained from American Type Culture Collection, no. CRL-1634 (ATCC, Manassas, VA). All cells were maintained in complete media: Dulbecco’s Modified Eagle’s Medium (DMEM) (Gibco) containing 10% (v/v) fetal bovine serum (FBS) (Gibco). In all experiments, HFF cells were infected with human cytomegalovirus (HCMV) strain Merlin(R1111), a kind gift from Richard Stanton (Cardiff University Medical School)^53^, in the presence or absence of compounds at a multiplicity of infection (MOI) of 1 for 96 h. Viral titers were determined by serial dilution of HCMV supernatant onto HFF monolayers, which were then covered in DMEM containing 5% (v/v) FBS, antibiotics, and 0.6% (w/v) methylcellulose. After incubation for 14 days, cells were stained with crystal violet, and plaques in the infected cell monolayers were counted.

#### Cell viability experiments in HFF cells

HFF cells were seeded at high (1x10^4^ cells per well) or low (1x10^3^ cells per well) numbers cells per well into 96 well plates without infection. High numbers of uninfected cells (1 x 10^4^ cells per well) were to assess cell viability, whereas low numbers of uninfected cells (1 x 10^3^ cells per well) were to assess both cell viability and cell proliferation. After overnight incubation to allow cell attachment, cells were treated for 96 hours with DMSO or compounds as indicated in the figures and text. MTT assays were carried out on cells in the wells of 96 well plates using a CyQUANT MTT cell viability Assay Kit (Invitrogen) according to the manufacturer’s instructions. In these assays, the ability of cellular NAD(P)H-dependent cellular oxidoreductase enzymes to reduce the tetrazolium dye 3-(4,5-dimethythiazo-2-yl)-2,5-diphenyltetrazolium bromide (MTT) to formazan was measured in a colorimetric assay, read on a GloMax Discover Microplate Reader (Promega).

## Author Contributions

All authors have given approval to the final version of the manuscript.

## Notes

The authors declare no competing financial interest.

## Acknowledgment

The SGC is a registered charity (number 1097737) that receives funds from Bayer AG, Boehringer Ingelheim, the Canada Foundation for Innovation, Eshelman Institute, Genentech, Genome Canada through Ontario Genomics Institute, EU/EFPIA/OICR/McGill/KTH/Diamond, Innovative Medicines Initiative 2 Joint Undertaking, Janssen, Merck KGaA (aka EMD in Canada and USA), Pfizer, the São Paulo Research Foundation-FAPESP, and Takeda. Research reported in this publication was supported in part by NIH U24DK116204, U54AG065187, and the NC Biotechnology Center Institutional Support Grant 2018-IDG-1030. J.L.C. was supported by the PhRMA Foundation (Crossref Funder ID: 100001797) Predoctoral Fellowship in Drug Discovery. Research reported in this publication was supported by the Office of the Director, NIH under award number S10OD032476 for upgrading the 500 MHz NMR spectrometer in the UNC Eshelman School of Pharmacy NMR Facility. B.L.S. was supported by New Investigator funds from St. George’s, University of London, a St. George’s Impact and Innovation Award and a PARK/WestFocus Award. The funders had no role in study design, data collection and analysis, decision to publish, or preparation of the manuscript. The analysis of the MIBs experiment was in part performed by the UNC Proteomics Core Facility, which is supported by the NCI Center Core Support Grant (2P30CA016086-45) to the UNC Lineberger Comprehensive Cancer Center. We wish to thank the National Cancer Institute Developmental Therapeutics Program (NCI/DTP) for the screening data of compounds **14** and **5** against their panel of 60 cancer cell lines.

The NanoBRET constructs used in this study were generously provided by Promega Corporation (Madison, WI). In addition to the work done at UNC, Promega performed some biological replicates of the K192 and all replicates of the K240 NanoBRET assay panels on **14**, **21**, and **5**. Dendrograms in Figure 5 and Supplemental Figure 6 were made using Kinmap. The depiction of the WNT signaling pathway in Figure 5 was made using Biorender. We thank Hassan Al-Ali for his support of this project and Richard Stanton for providing human cytomegalovirus strain Merlin.

## Abbreviations

ACN: acetonitrile
BSA: bovine serum albumin
CK1: Casein Kinase 1
DCM: dichloromethane
DIPA: *N, N*-diisopropylamine
DMEM: Dulbecco’s modified Eagle medium
DMF: *N, N*-dimethylformamide
DMSO: dimethyl sulfoxide
EtOAc: ethyl acetate
FBS: fetal bovine serum
FO: fractional occupancy
HCMV: human cytomegalovirus
HPLC: high-performance liquid chromatography
IC_50_: half maximal inhibitory concentration
K_m_: Michaelis constant
k192: panel of 192 kinases via NanoBRET assays
k240: panel of 240 kinases via NanoBRET assays
LC–MS: liquid chromatography–mass spectrometry
LRP6: low-density lipoprotein receptor-related protein 6
MLKL: mixed lineage kinase-like pseudokinase
NaLTSA: nanoluciferase-based thermal shift assay
NanoBRET: bioluminescence resonance energy transfer using NanoLuciferase
NB: NanoBRET
NLuc: NanoLuciferase
NMR: nuclear magnetic resonance
POC: percent of control
RIPK3: receptor-interacting serine/threonine kinase 3
SAR: structure–activity relationships
SD: standard deviation
SGC: Structural Genomics Consortium
SNAr: nucleophilic aromatic substitution
TEA: triethylamine
TFA: trifluoroacetic acid
v/v: volume for volume.

