## Supplemental Information for "Development of a chemical probe to enable characterization of the casein kinase 1γ subfamily"

### Table of Contents

| | |
|---|---|
| Figure S1 | S1 |
| Figure S2 | S2 |
| Figure S3 | S3 |
| Figure S4 | S4 |
| Figure S5 | S5-S6 |
| Figure S6 | S7-S8 |
| Figure S7 | S9 |
| Figure S8 | S10 |
| Figure S9 | S11 |
| Figure S9 | S12 |
| Figure S10 | S13-S14 |
| Purity traces and spectra for all compounds | S15–S59 |

**Figure S1:**

A

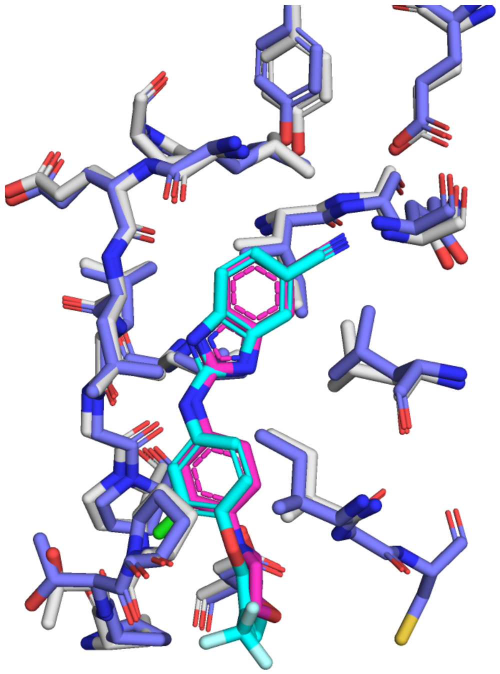

B

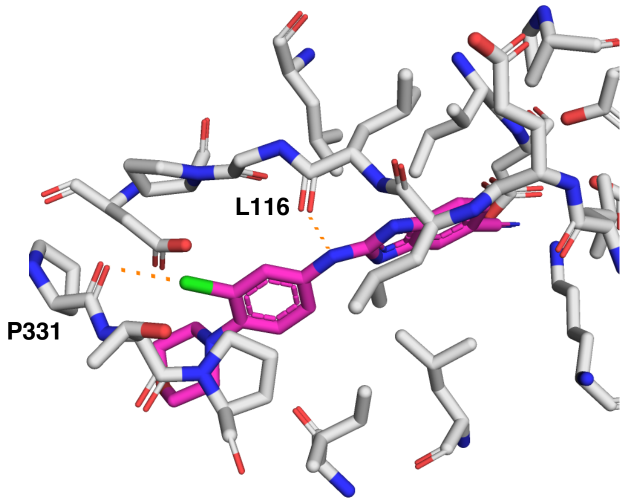


C

**SGC-CK1γ-1 with CK1γ3**

| Property | Values |
|---|---|
| Predicted TM-Score | 0.836 |
| Interfaced Predicted TM-Score | 0.99 |
| Confidence Score | 0.8 |
| Average pLDDT | 0.753 |
| Affinity Probability | 0.843 |
| Predicted $IC_{50}$ | 36.8 nM |

**SGC-CK1γ-1N with CK1γ3**

| Property | Values |
|---|---|
| Predicted TM-Score | 0.773 |
| Interfaced Predicted TM-Score | 0.975 |
| Confidence Score | 0.794 |
| Average pLDDT | 0.749 |
| Affinity Probability | 0.478 |
| Predicted $IC_{50}$ | 1220 nM |

**Figure S1**. Co-folding of SGC-CK1γ-1 with CK1γ3. a) Overlay of the Boltz-2 co-folding of SGC-CK1γ-1 (magenta) with CK1γ3 (blue) and a literature crystal structure of CK1γ3 (grey) with an analog of compound 1h bearing a cyanobenzimidazole moiety (cyan, Figure 1D). RMSD between crystal structure (PDB code: 4G16) and the Boltz-2 model: 1.16 Å over 293 shared residues.[16,31] b) Co-folding model of CK1γ3 predicted with SGC-CK1γ-1 with projected interactions (orange), such as a putative halogen bond Pro331 and a putative hydrogen bonding with Leu116. c) Properties of the Boltz-2 model for co-folding of SGC-CK1γ-1 and SGC-CK1γ-1N with CK1γ3. Green represents high confidence, yellow represents medium confidence, and red represents low confidence. These colors correspond with the reliability of this model as determined by the associated software (Rowan Scientific).

#### Figure S2:

A

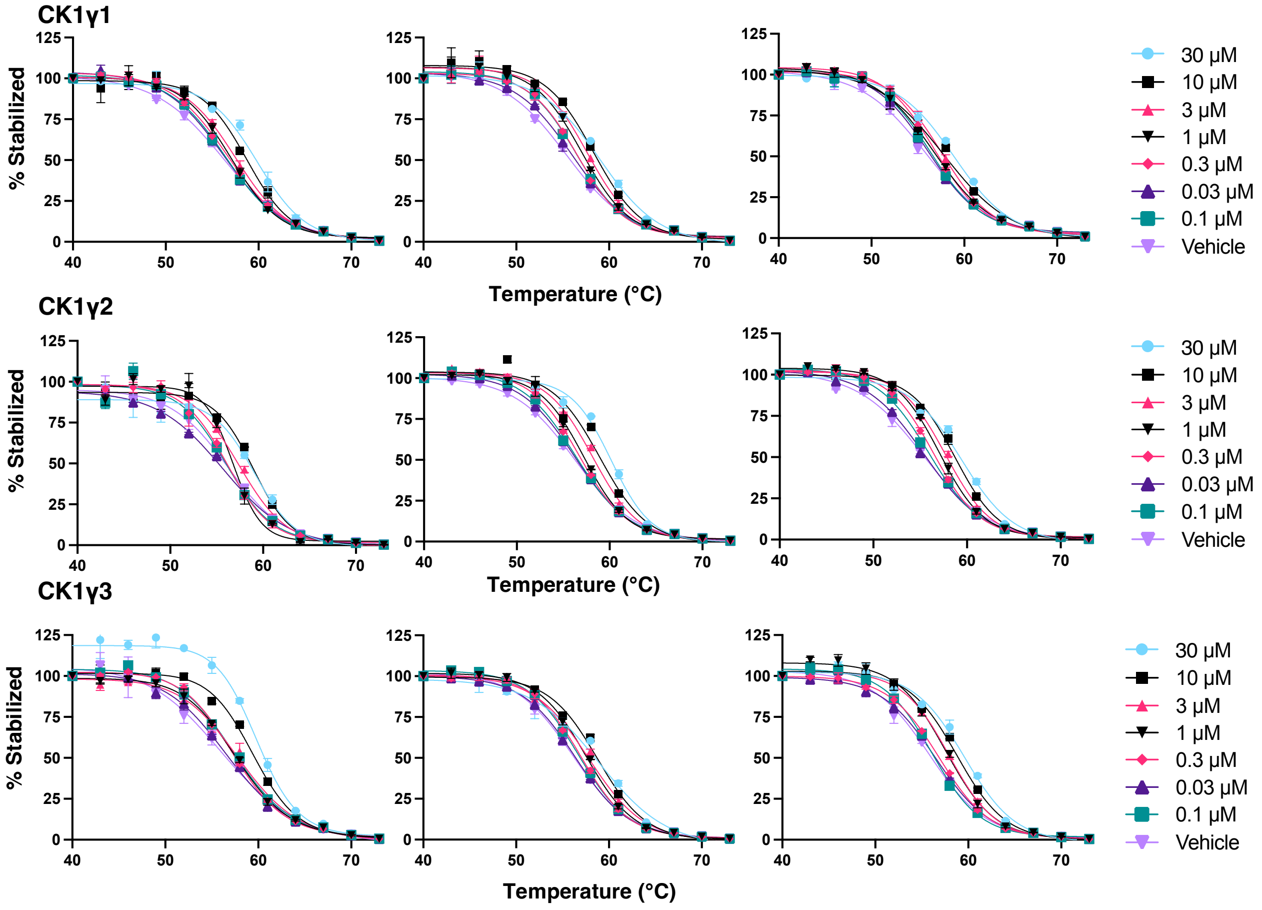


B

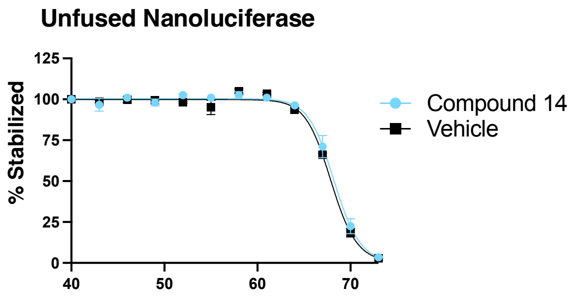


**Figure S2**. Curves from NaLTSA replicate and control experiments. a) Biological replicates of the gradient dose range for compound **14** versus all CK1γ isoforms. b) Data generated when cells transfected with unfused nanoluciferase were treated with either 30 μM of compound **14** or DMSO.

**Figure S3:**

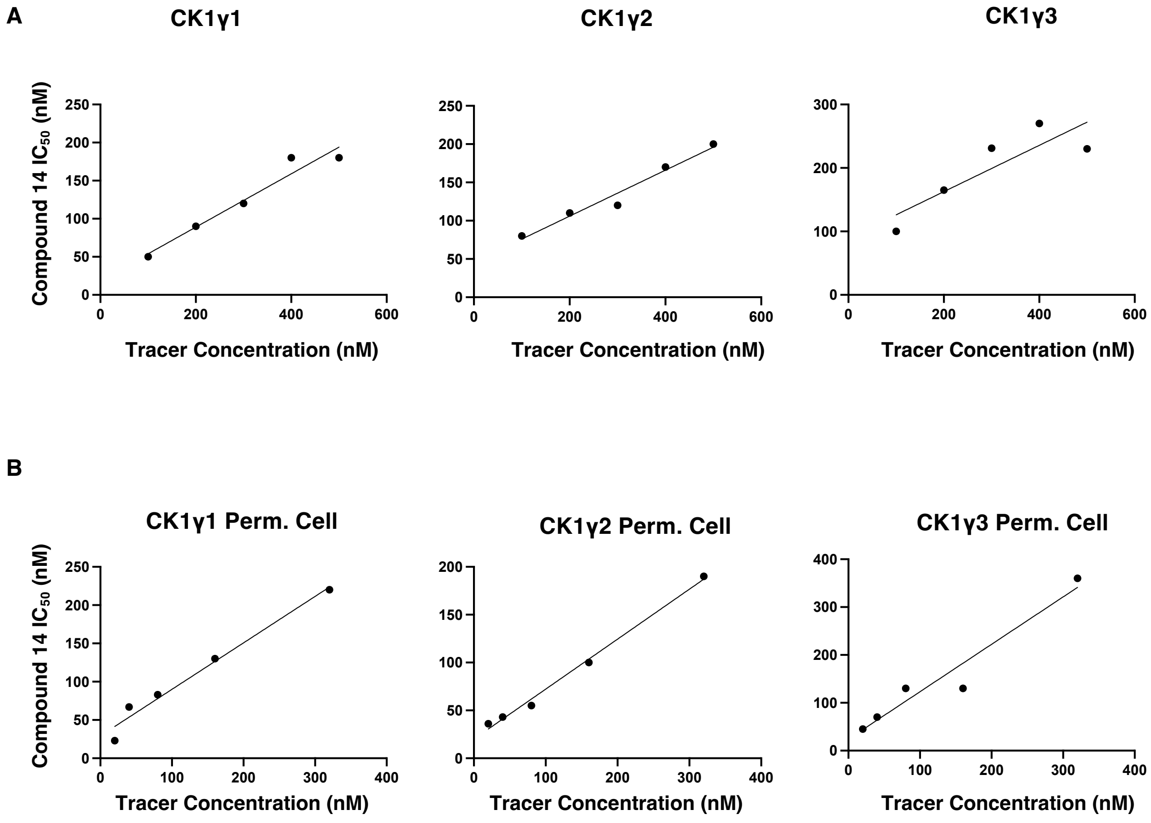


**Figure S3**. Confirmation that all assays adhere to the linear decline is $IC_{50}$ with reduced tracer concentration described by the Cheng-Prusoff equation. a) Linear regression between tracer **23** concentration and $IC_{50}$ of compound **14** in the intact cell NanoBRET assay (N=3). b) Linear regression between tracer **23** concentration and $IC_{50}$ of compound **14** in the permeabilized cell NanoBRET assay (N=2). Perm = permeabilized.

**Figure S4:**

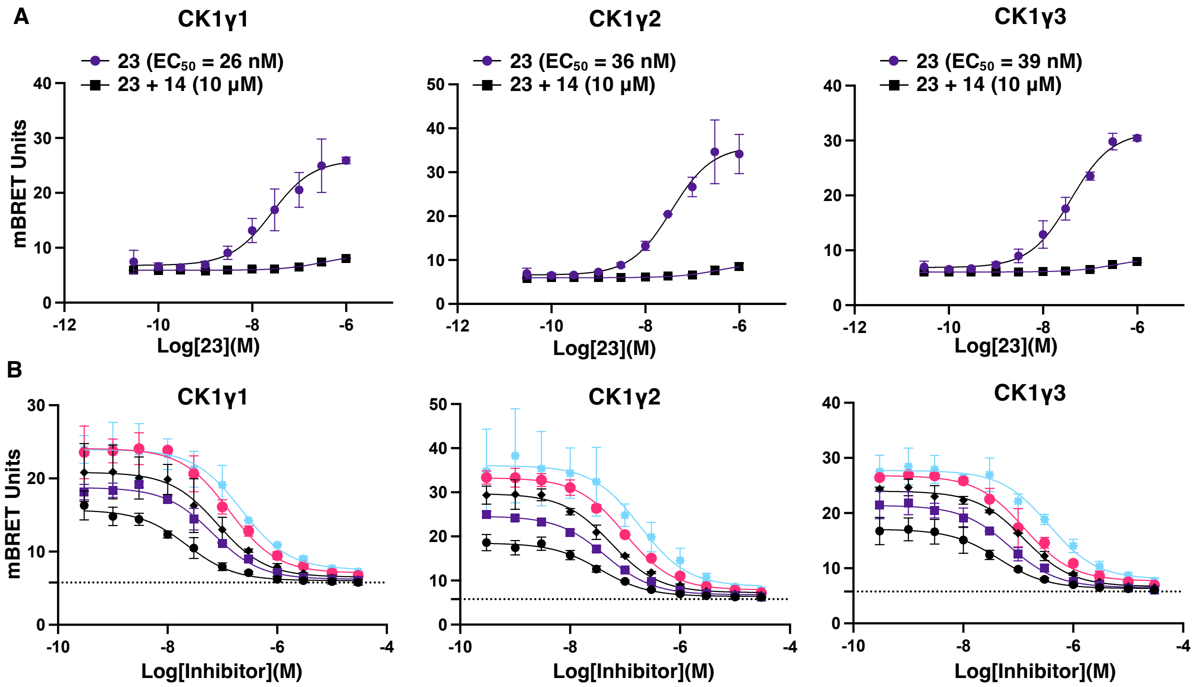


**CK1γ1**

| | | | | | |
|---|---|---|---|---|---|
| Tracer (nM) | 20 | 40 | 80 | 160 | 320 |
| 14 $IC_{50}$ (nM) | 23 | 67 | 83 | 130 | 220 |

**CK1γ2**

| | | | | | |
|---|---|---|---|---|---|
| Tracer (nM) | 20 | 40 | 80 | 160 | 320 |
| 14 $IC_{50}$ (nM) | 36 | 43 | 55 | 100 | 190 |

**CK1γ3**

| | | | | | |
|---|---|---|---|---|---|
| Tracer (nM) | 20 | 40 | 80 | 160 | 320 |
| 14 $IC_{50}$ (nM) | 45 | 70 | 130 | 130 | 360 |

**Figure S4**. Development of the NanoBRET assay in permeabilized cells. a) In digitonin-treated cells, 11-point dose–response experiments with tracer **23** in the presence or absence of 10 µM compound **14** (N =2). Error bars correlate with SD. b) Tracer titrations for tracer **23** and compound **14** in permeabilized cells with an $IC_{50}$ displayed for each tracer concentration. All data are reported as N=2. Error bars correlate with SD. The data is not background-subtracted, and a horizontal dotted line is present to indicate the background level.

#### Figure S5

A

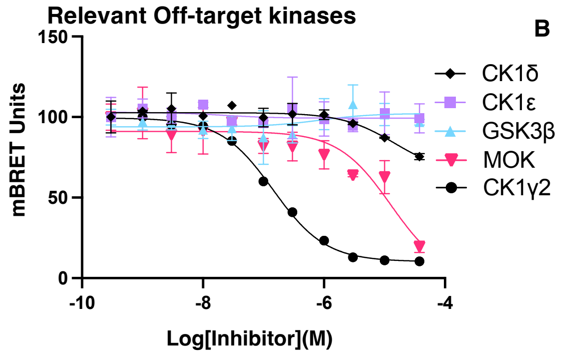


B

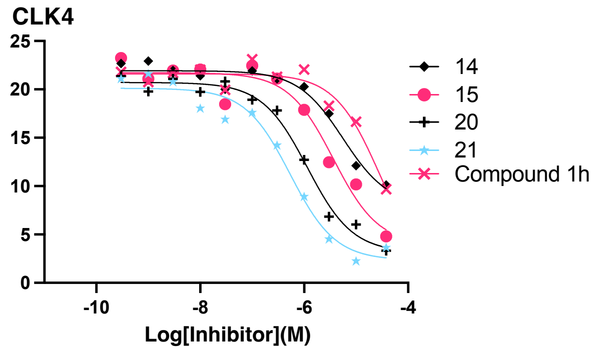


| Kinase | NB $IC_{50}$ (nM) |
|---|---|
| MOK | >5000 |
| CK1δ | >30000 |
| CK1ε | >30000 |
| GSK3β | >30000 |
| CK1γ2 | 140 |

| Compound | NB $IC_{50}$ (nM) |
|---|---|
| 14 | >10000 |
| 15 | 3600 |
| 20 | 1200 |
| 21 | 530 |
| Compound 1h | >10000 |

C

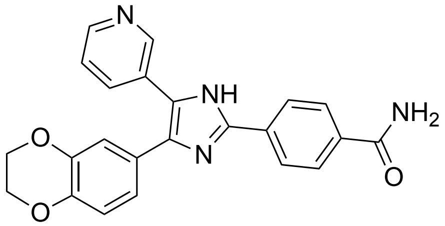

D4476
Pan-CK1 inhibitor

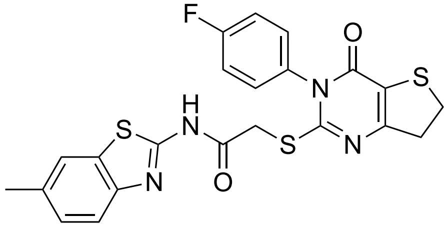

IWP-3
Porcupine inhibitor that weakly inhibits CK1γ3

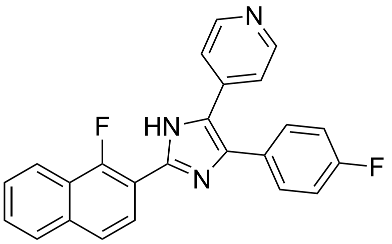

CK1-IN-1
CK1δ and CK1ε inhibitor

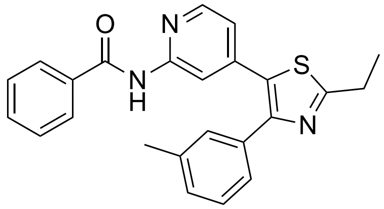

TAK-715
P38, CK1δ, and CK1ε inhibitor

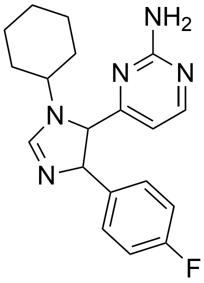

PF-670462
CK1δ, and CK1ε inhibitor

**Figure S5**. NanoBRET curves generated for off-target kinases of SGC-CK1γ-1 and **21** based on in-cell selectivity screening. a) 11-point dose–response NanoBRET assay screen for compound **14** against WNT signaling kinases enabled for NanoBRET and MOK (N=1). Error bars correlate with SD. b) 11-point dose–response CLK4 NanoBRET assay screens for compounds **14**, **15**, **20**, **21**, and the Amgen parent compound **1** (N=1). Error bars correlate with SD. NB = NanoBRET. c) Chemical structures of all tested CK1 literature inhibitors with descriptions of each compound.

**Figure S6:**

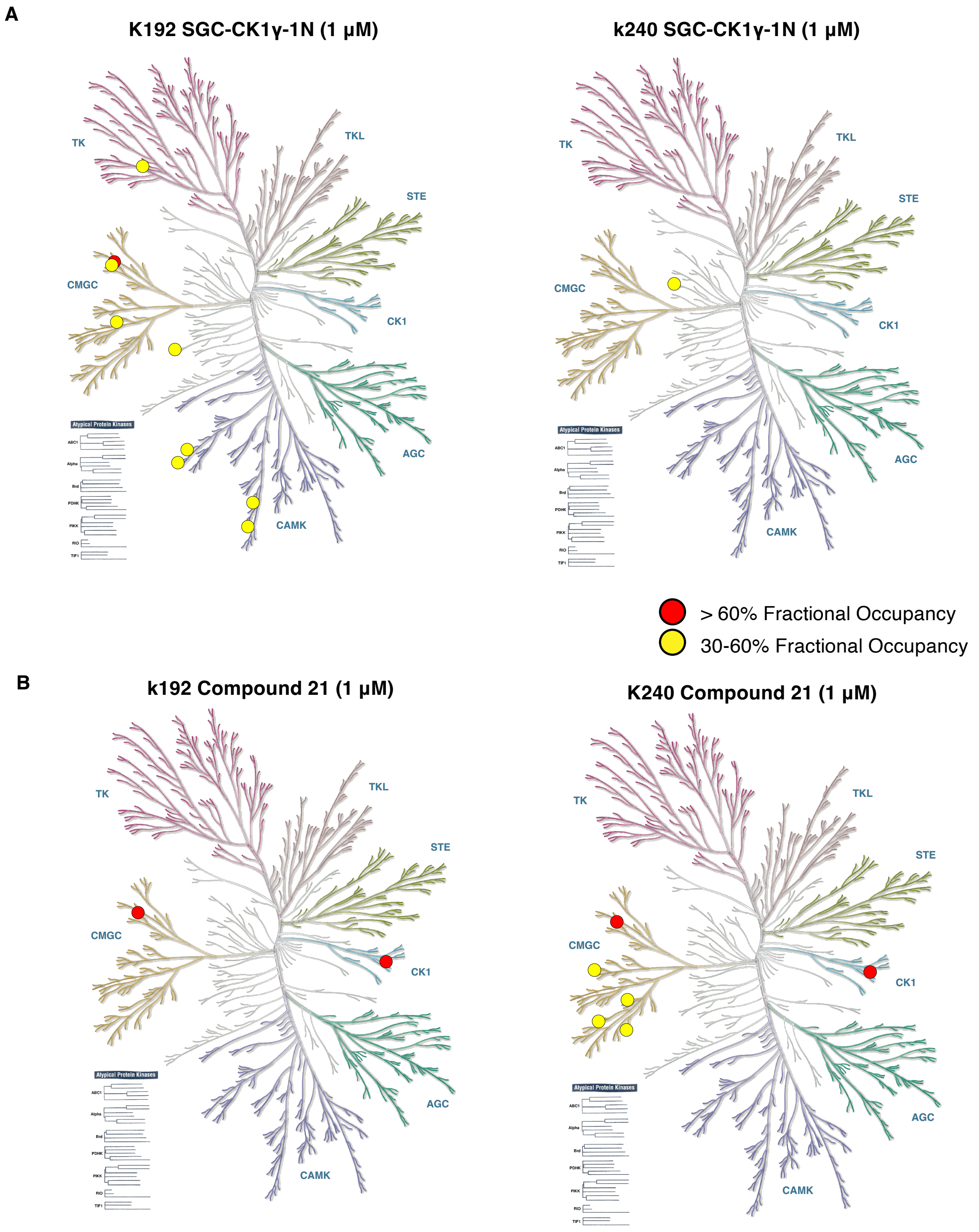
A
K192 SGC-CK1γ-1N (1 µM)
k240 SGC-CK1γ-1N (1 µM)
TK
TKL
STE
CMGC
CK1
AGC
CAMK
Atypical Protein Kinases
> 60% Fractional Occupancy
30-60% Fractional Occupancy
B
k192 Compound 21 (1 µM)
K240 Compound 21 (1 µM)

**Figure S6**. In-cell selectivity of SGC-CK1γ-1N and compound **21** in the K192 and K240 assays. a) Dendrogram displaying all kinases that bound to SGC-CK1γ-1N (1 μM) with greater than 30% F.O. in the K192 (left) and K240 (right), with kinases demonstrating between 30% and 60% F.O. in yellow and kinases demonstrating greater than 60% F.O. depicted in red. b) Dendrogram displaying all kinases that bound to compound **21** (1 μM) with greater than 30% F.O. in the K192 (left) and K240 (right), with kinases demonstrating between 30% and 60% F.O. in yellow and kinases demonstrating greater than 60% F.O. depicted in red.

**Figure S7:**

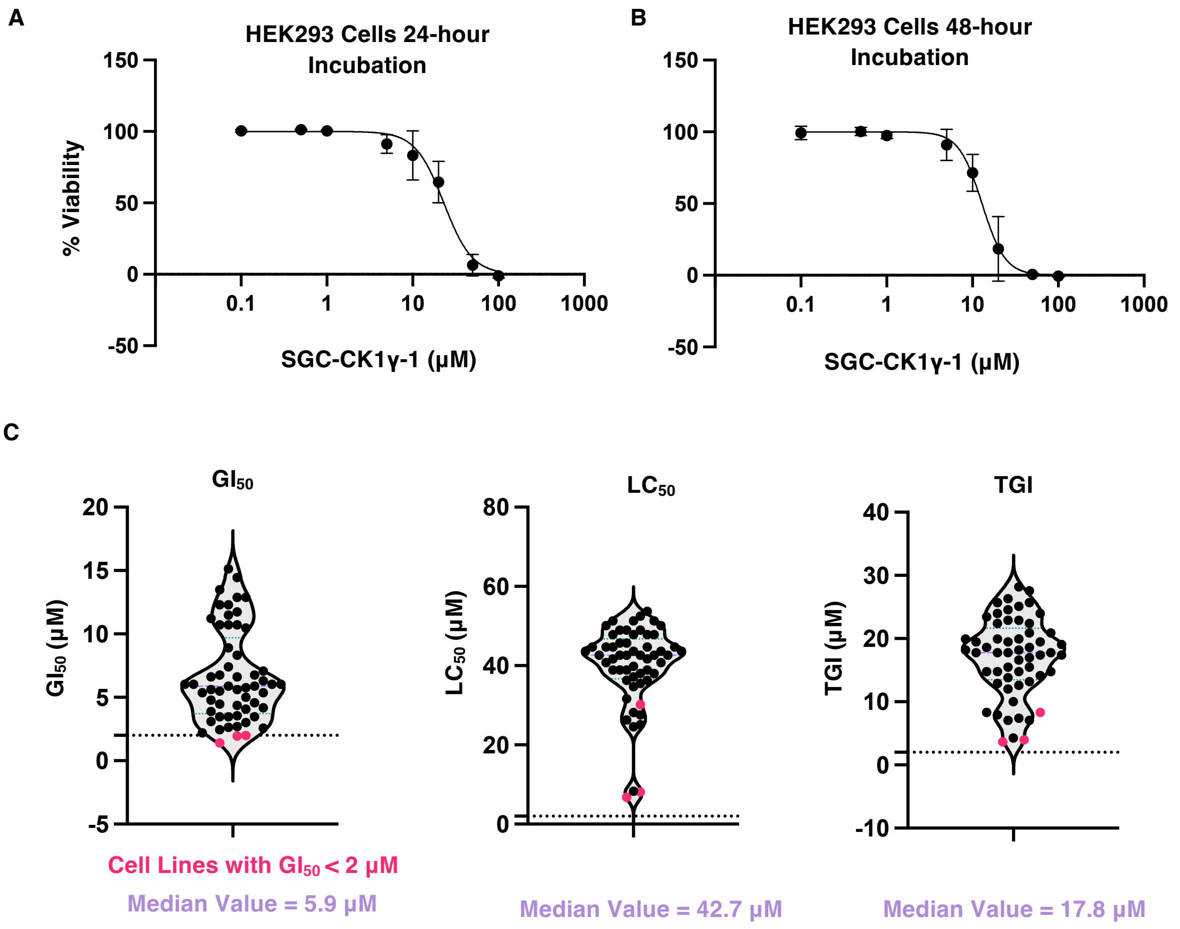


**Figure S7**. Toxicity assessment of SGC-CK1γ-1 against a broad panel of cell lines. a) Cell Titer Glo results from an experiment with 24-hour exposure of SGC-CK1γ-1 to HEK293 cells at varied doses (N=2). Error bars represent SD. Cell Titer Glo results from an experiment with 48-hour exposure of SGC-CK1γ-1 to HEK293 cells at varied doses (N=1). Error bars represent SD. c) Violin plots of growth inhibition 50% ($GI_{50}$), lethal concentration 50% ($LC_{50}$), and total growth inhibition (TGI) across a panel of 60 cancer cell lines performed by the National Cancer Institute (NCI-60). Three leukemia cell lines with a $GI_{50}$ < 2 µM are highlighted in pink: MOLT-4, HL-60(TB), and SR. A dotted line marking the 2 µM threshold is present in all graphs. Median values for $GI_{50}$, $LC_{50}$, and TGI are indicated beneath each violin plot in purple.

**Figure S8:**

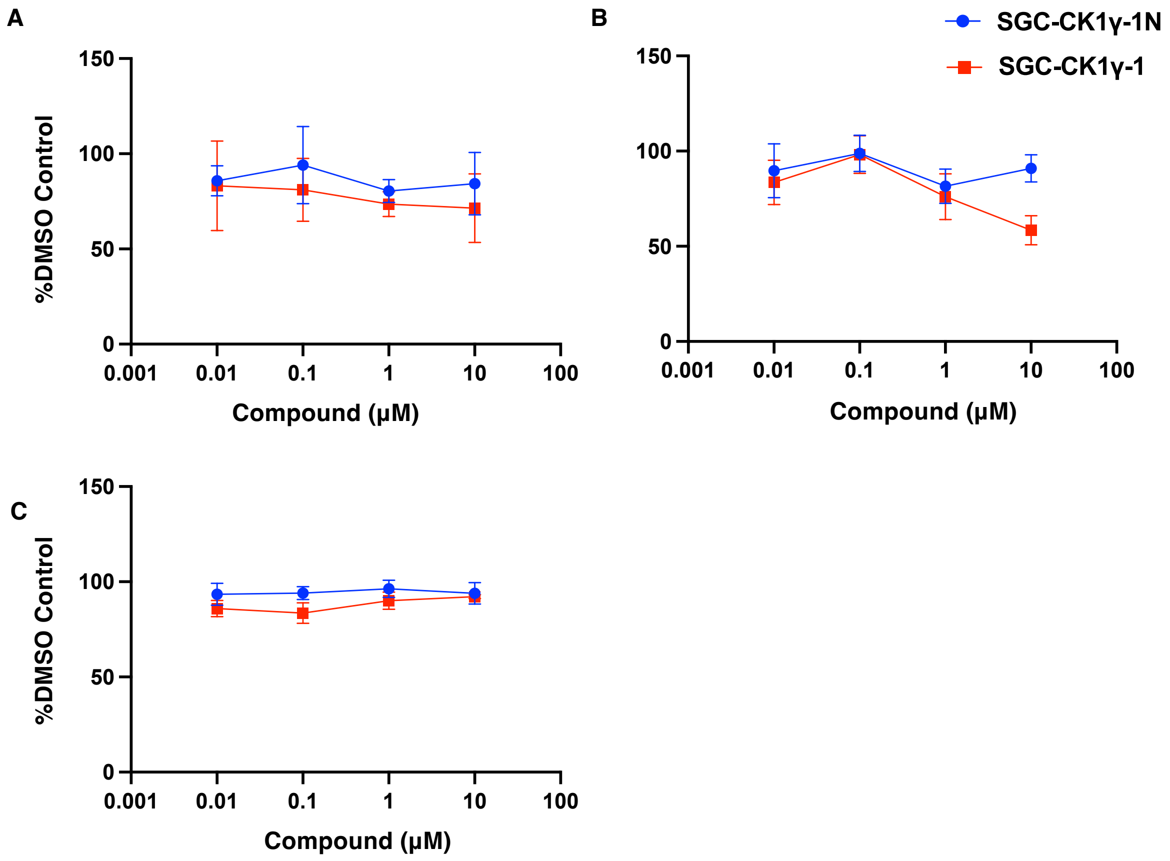


**Figure S8**. Plots of HCMV replication data in the presence of SGC-CK1γ-1. a,b) A high (a) or low (b) concentration of uninfected HFF cells were treated for 96 hours with a 10-fold dilution series of either SGC-CK1γ-1 or SGC-CK1γ-1N, or the corresponding volume of DMSO, and then examined using an MTT assay. Cell viability in the presence of either compound is shown as the percentage viability compared to the appropriate DMSO control (N=3). Error bars represent SD. c) HFF cells were infected with HCMV strain Merlin (R1111) and treated with various concentrations of either SGC-CK1γ-1 or SGC-CK1γ-1N, or the corresponding volume of DMSO, 24 hours post-infection (d) (N=3). Error bars represent SD.

**Figure S9:**

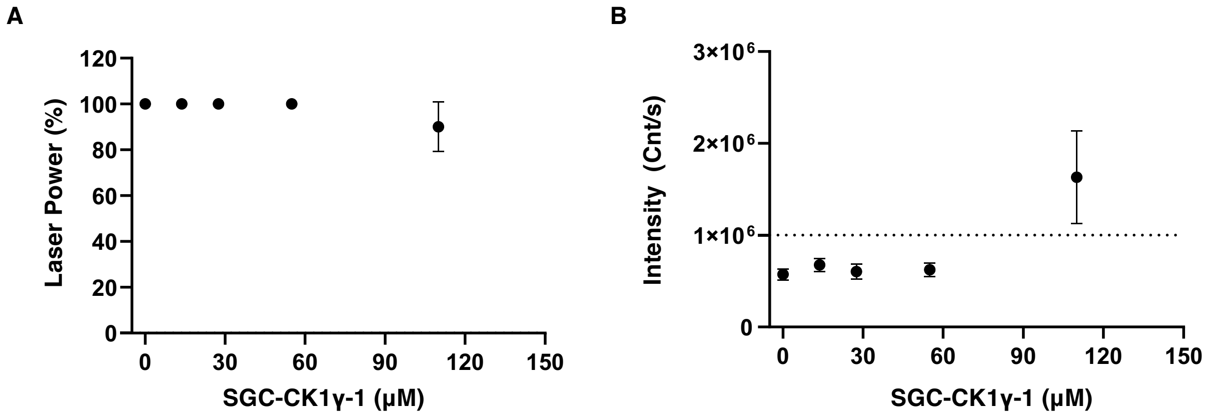


**Figure S9**. Dynamic light scatter data for SGC-CK1γ-1 confirms it has good aqueous solubility, even at high micromolar concentrations. (a) Instrument-adjusted laser power (%). (b) Scattering intensity (counts per second, Cnt/s). Scattering intensities exceeding $1 \times 10^6$ Cnt/s (dashed line in panel b) were considered indicative of compound aggregation. All measurements were performed in triplicate.

#### Figure S10

**A**

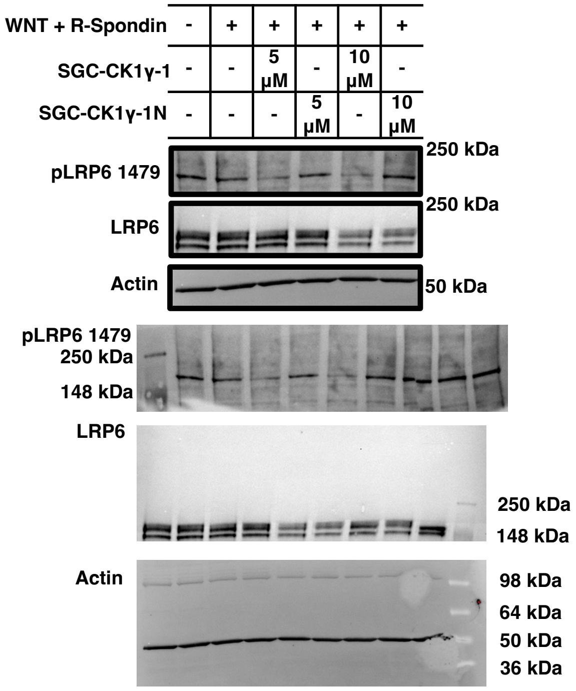


**Lanes: 1. DMSO, 2. WNT+DMSO, 3. WNT+ SGC-CK1γ-1 (5 μM), 4. SGC-CK1γ-1N (5 μM), 5. SGC-CK1γ-1 (10 μM), 6. SGC-CK1γ-1N (10 μM), 7-9. Untested lysate (Ladder on left or Right indicated by molecular weight markers)**

**B**

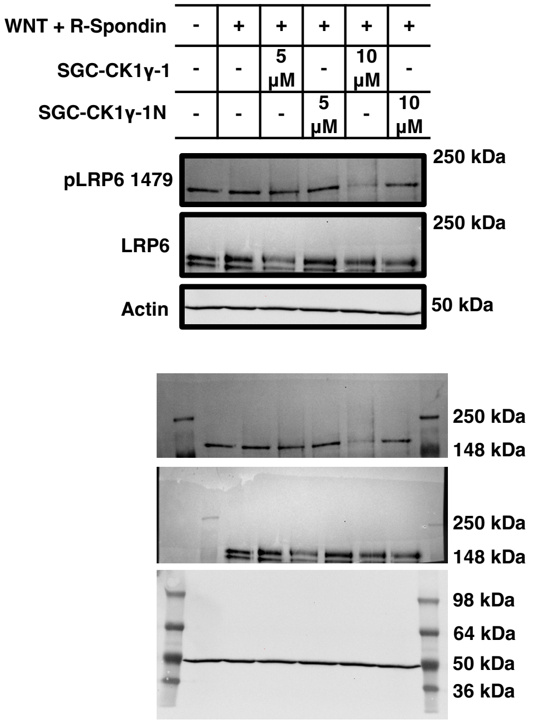


**Lanes: 1. DMSO, 2. WNT+DMSO, 3. WNT+ SGC-CK1γ-1 (5 μM), 4. SGC-CK1γ-1N (5 μM), 5. SGC-CK1γ-1 (10 μM), 6. SGC-CK1γ-1N (10 μM)**

**Figure S10 Continued:**

C

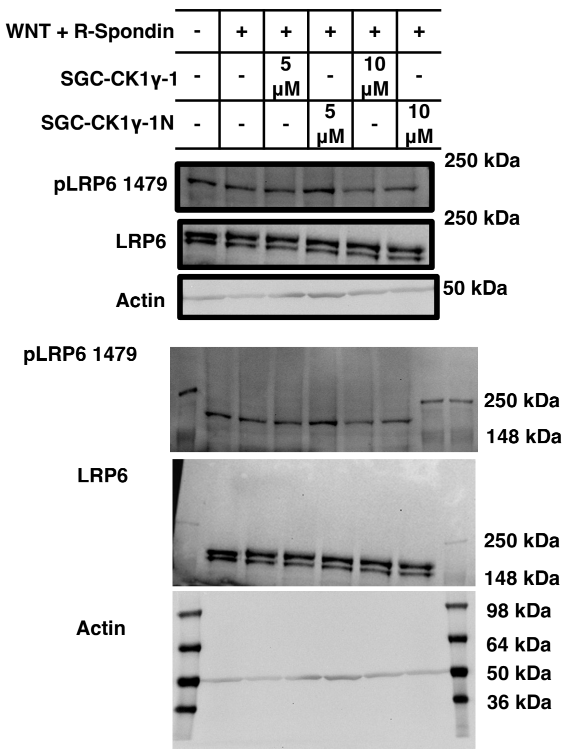


Lanes: 1. DMSO, 2. WNT + DMSO, 3. WNT + SGC-CK1γ-1 (5 μM), 4. SGC-CK1γ-1N (5 μM), 5. SGC-CK1γ-1 (10 μM), 6. SGC-CK1γ-1N (10 μM)

D

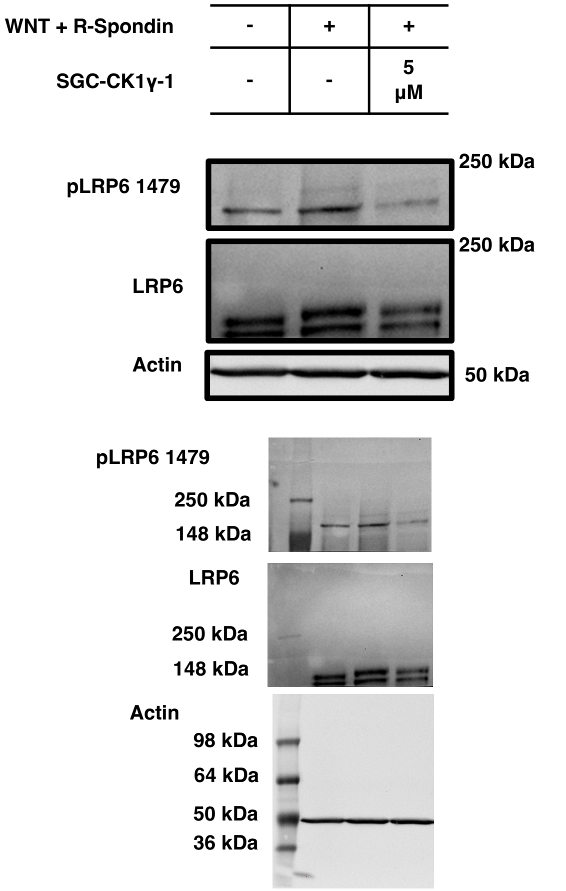


Lanes: 1. DMSO, 2. WNT+DMSO, 3. WNT + SGC-CK1γ-1 (5 μM)

**Figure S10**. Uncropped western blot images. a-d) Cropped (above) and full (below) blots corresponding to Figure 5B, and for all biological replicates depicted in the quantification in Figure 5C.

$^{1}H$ NMR (400 MHz, DMSO-$d_6$) for Compound **1**

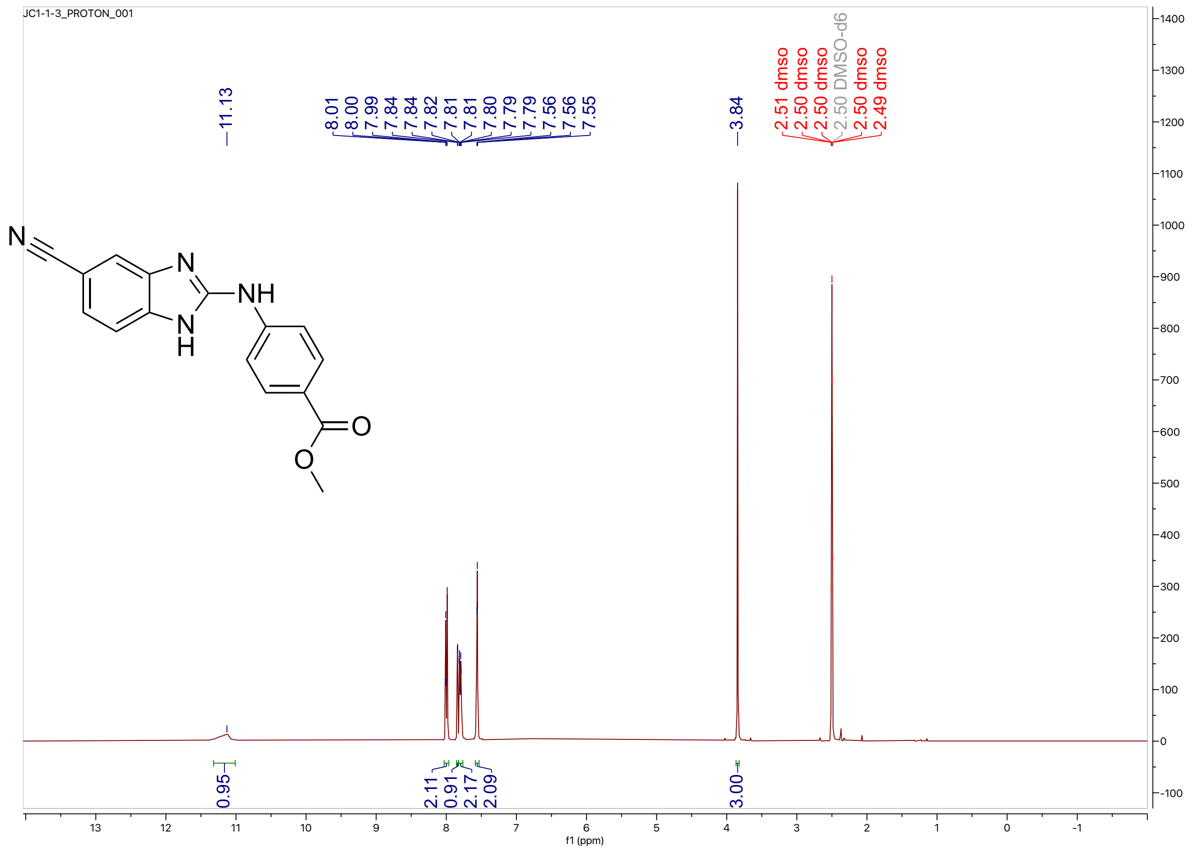


$^{13}C$ NMR (100 MHz, DMSO-$d_6$) for Compound **1**

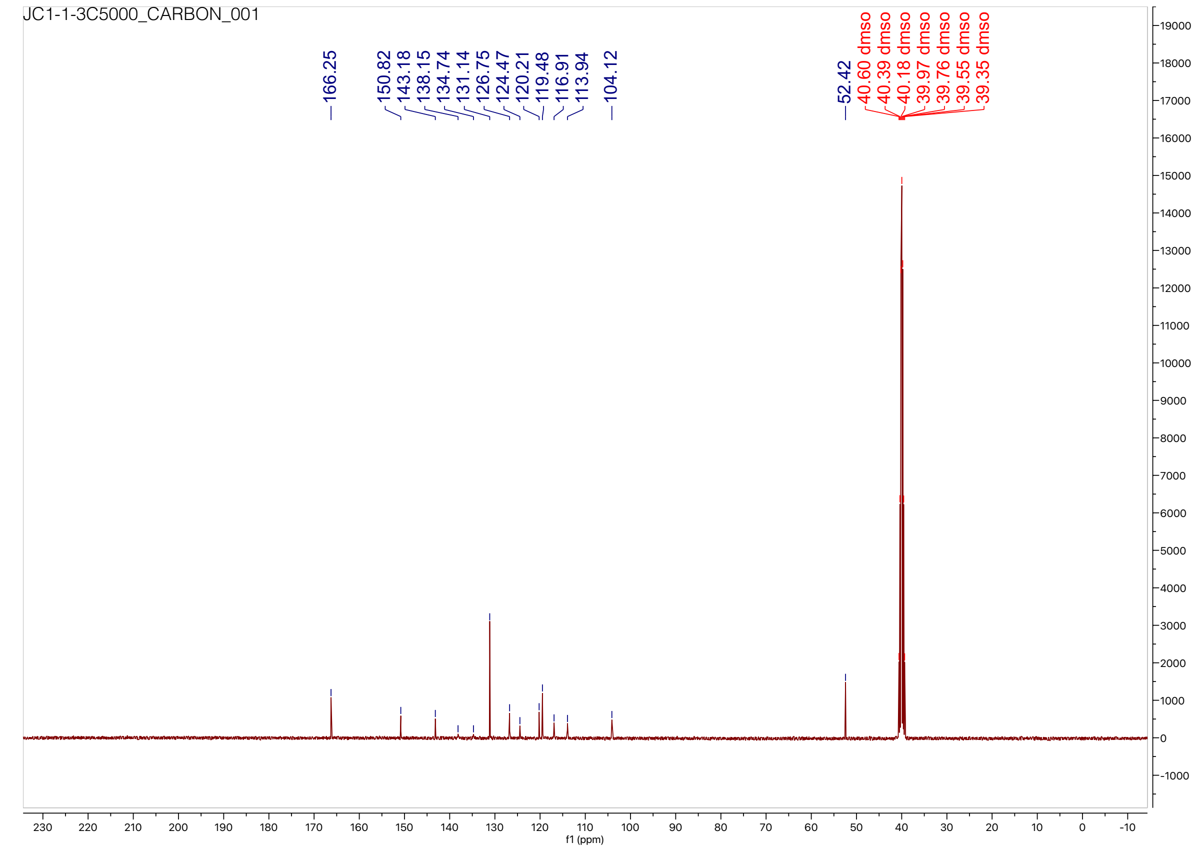

HPLC trace for compound **1**

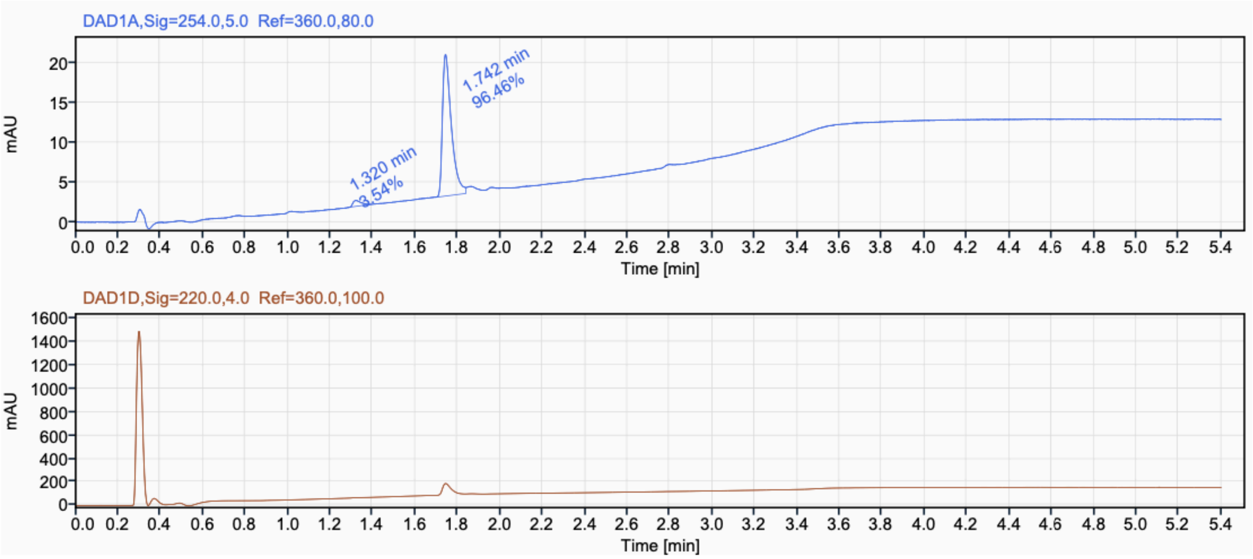

$^{1}H$ NMR (400 MHz, DMSO-$d_6$) for Compound **2**

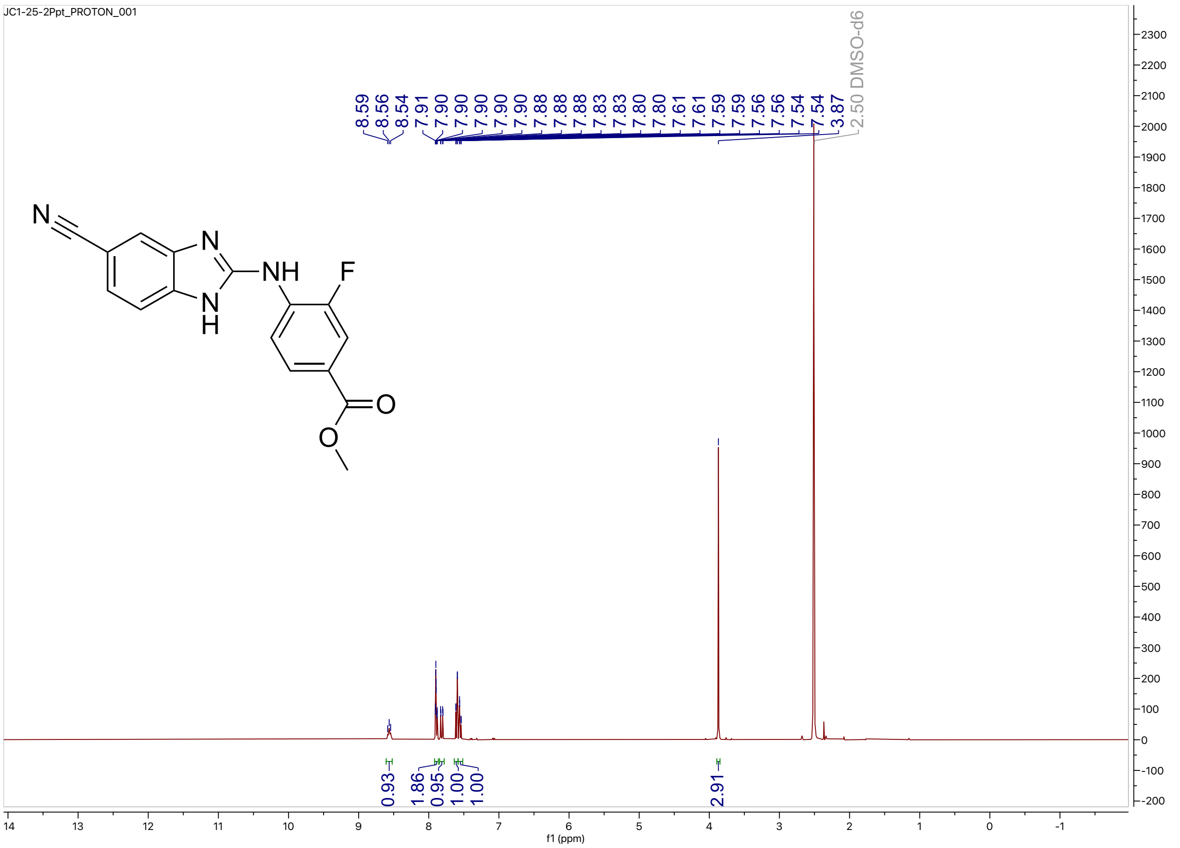


$^{13}C$ NMR (100 MHz, DMSO-$d_6$) for Compound **2**

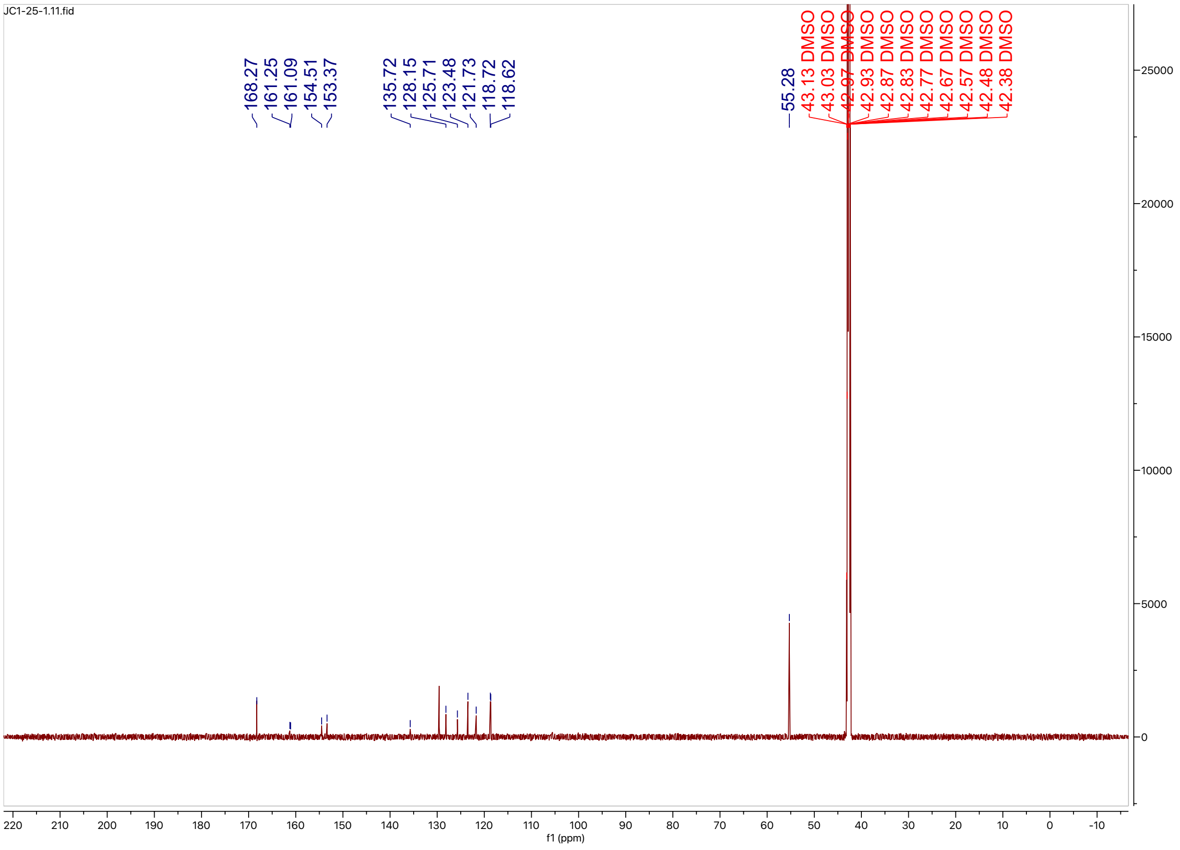

HPLC trace for compound **2**

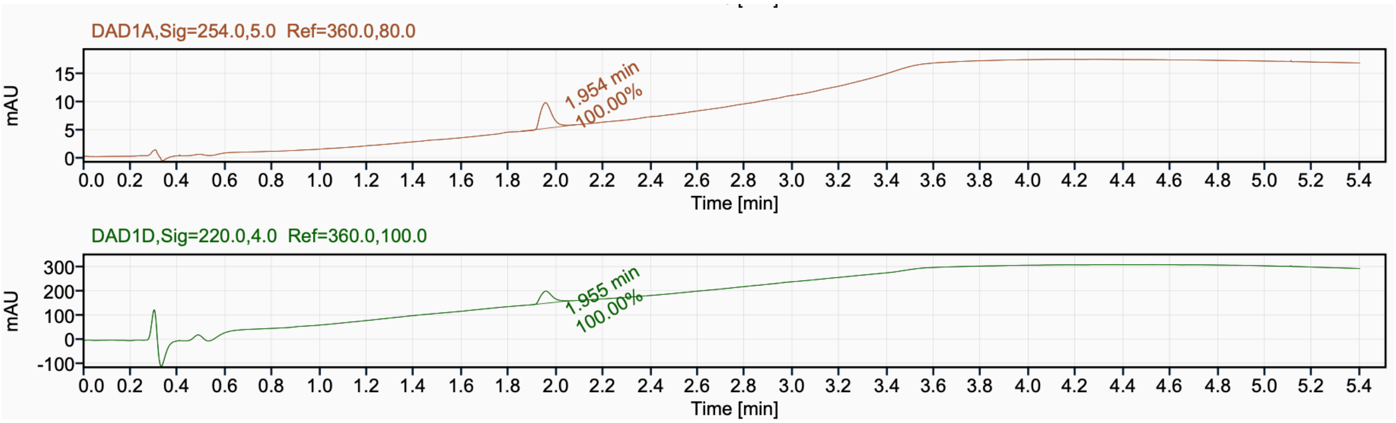

$^{1}H$ NMR (400 MHz, $CD_3OD$) for Compound **3**

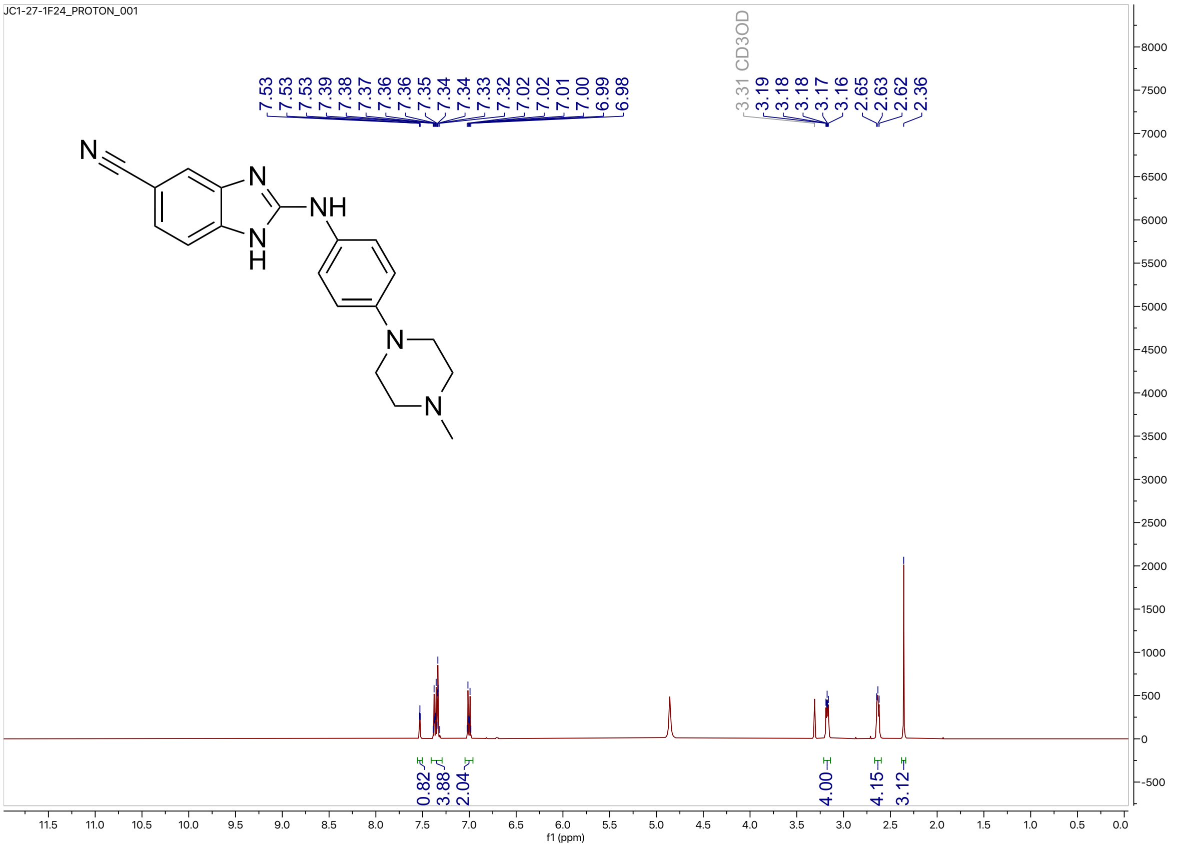


$^{13}C$ NMR (214 MHz, $CD_3OD$) for compound **3**

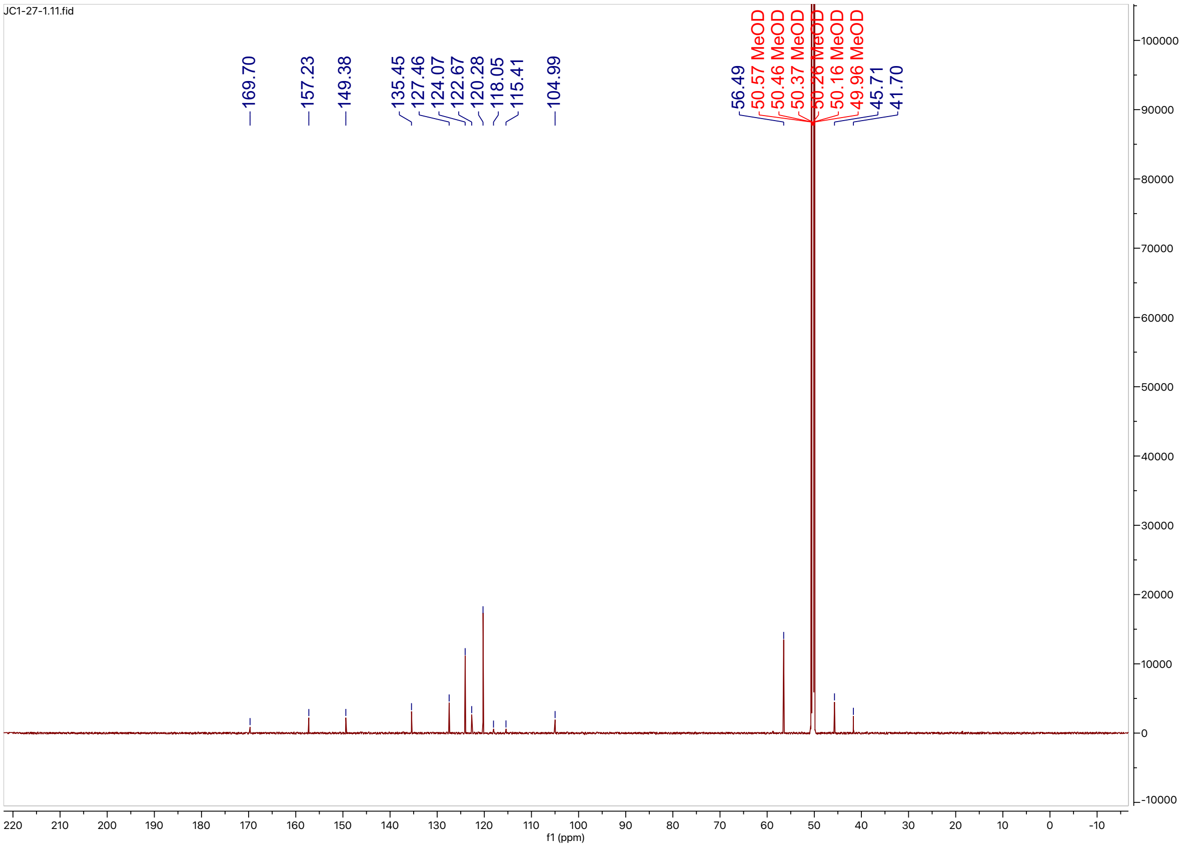

HPLC trace for compound **3**

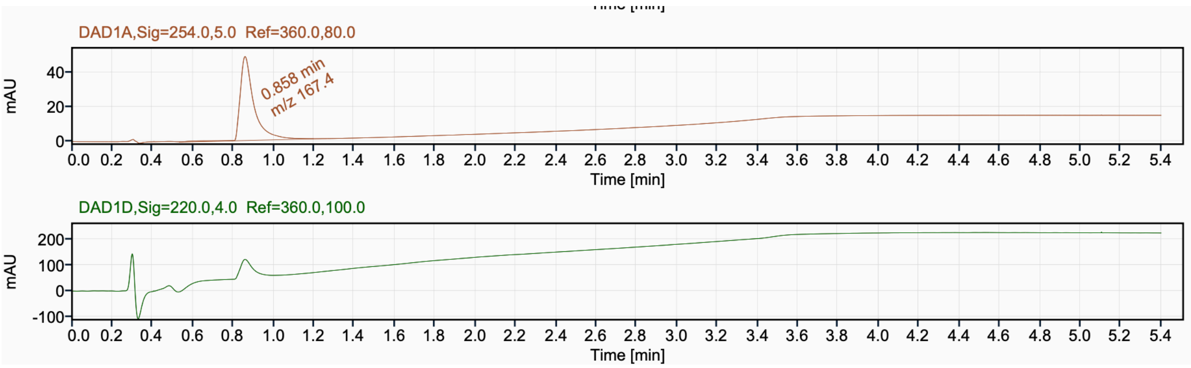

$^{1}H$ NMR (400 MHz, $CD_3OD$) for Compound **4**

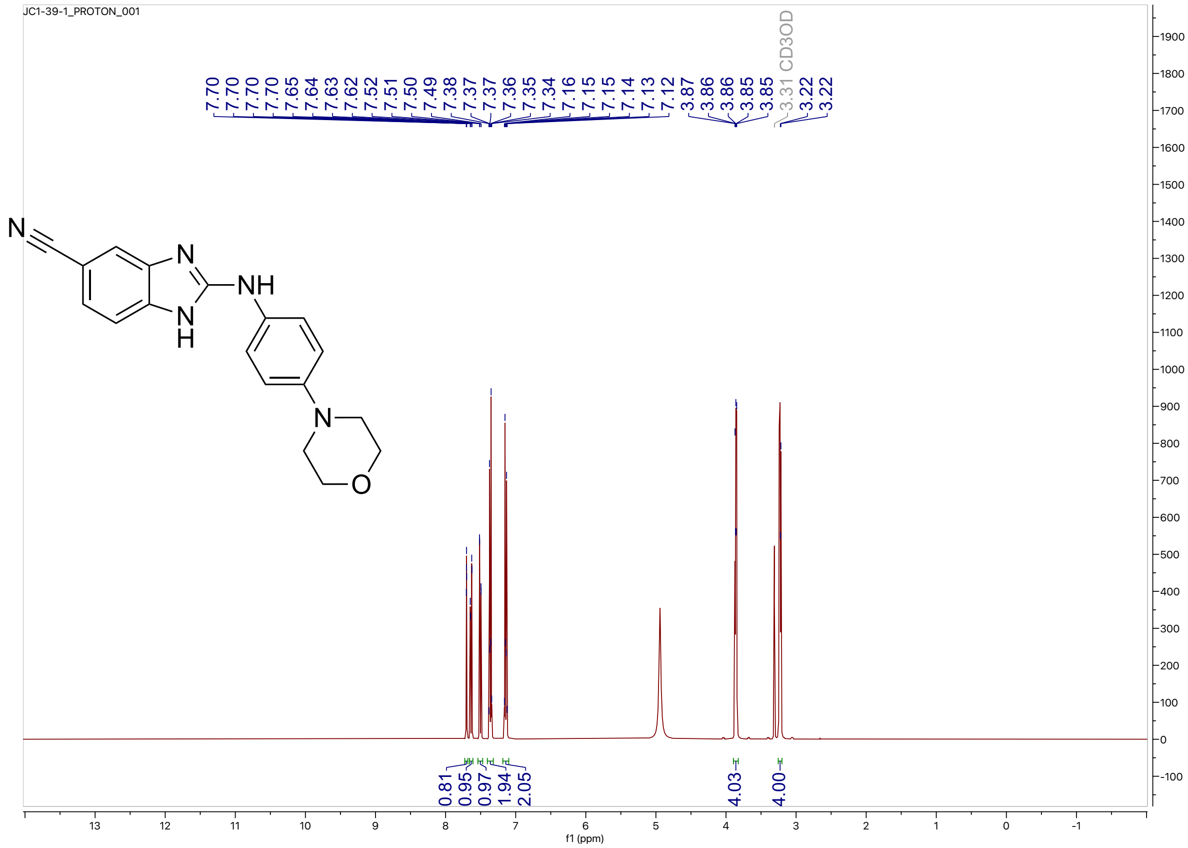


$^{13}C$ NMR (100 MHz, $CD_3OD$) for Compound **4**

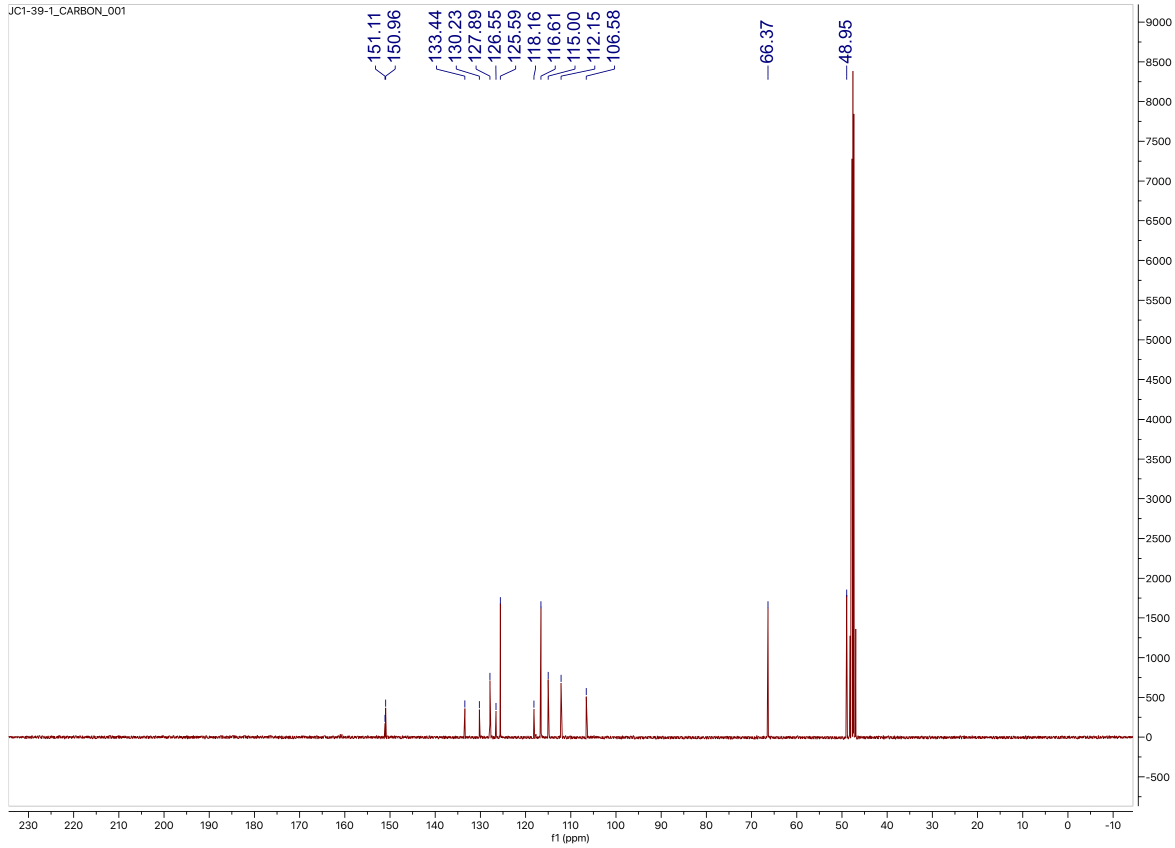

HPLC trace for compound **4**

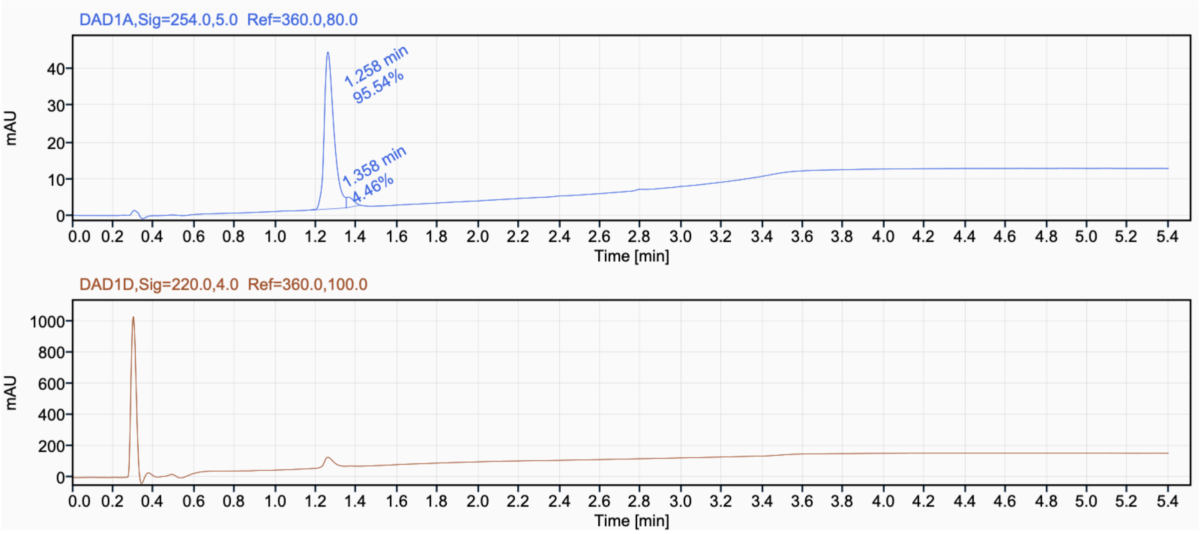

$^{1}H$ NMR (400 MHz, $CD_3OD$) for Compound **5**

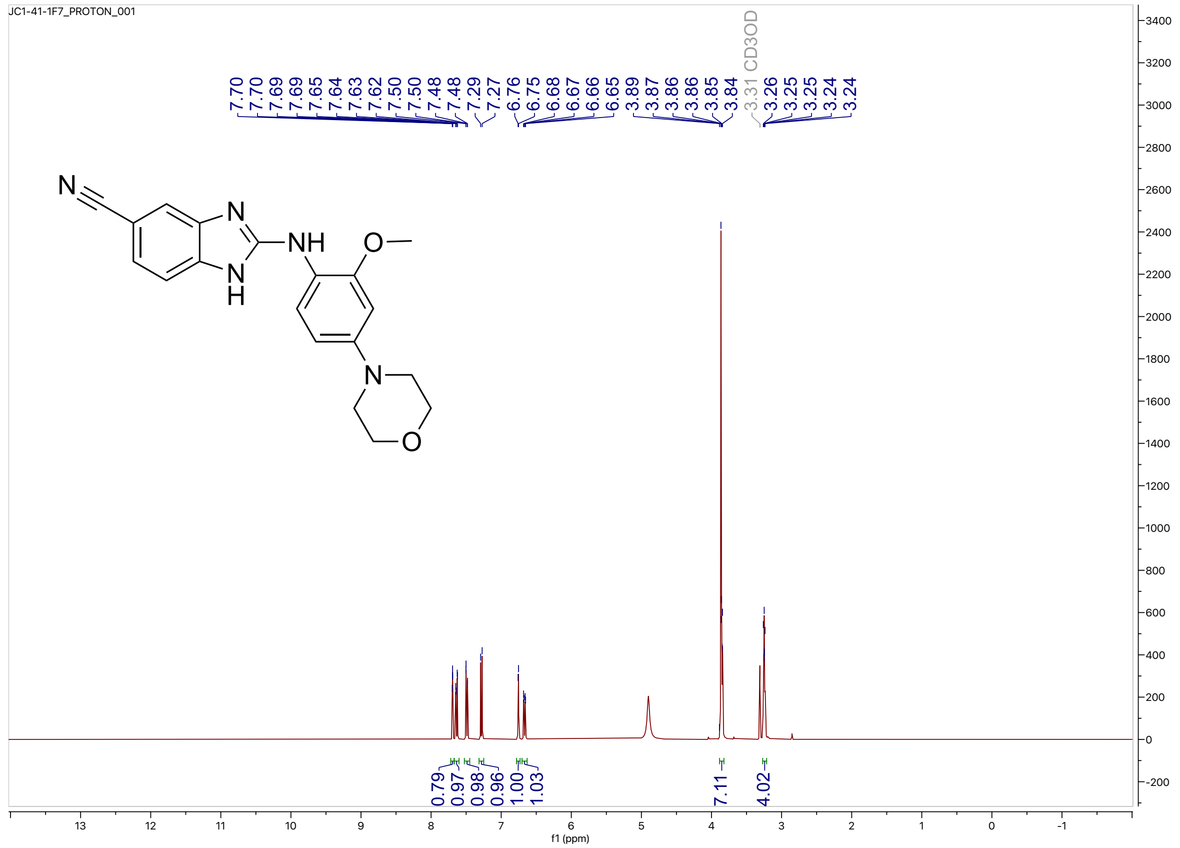


$^{13}C$ NMR (100 MHz, $CD_3OD$) for Compound **5**

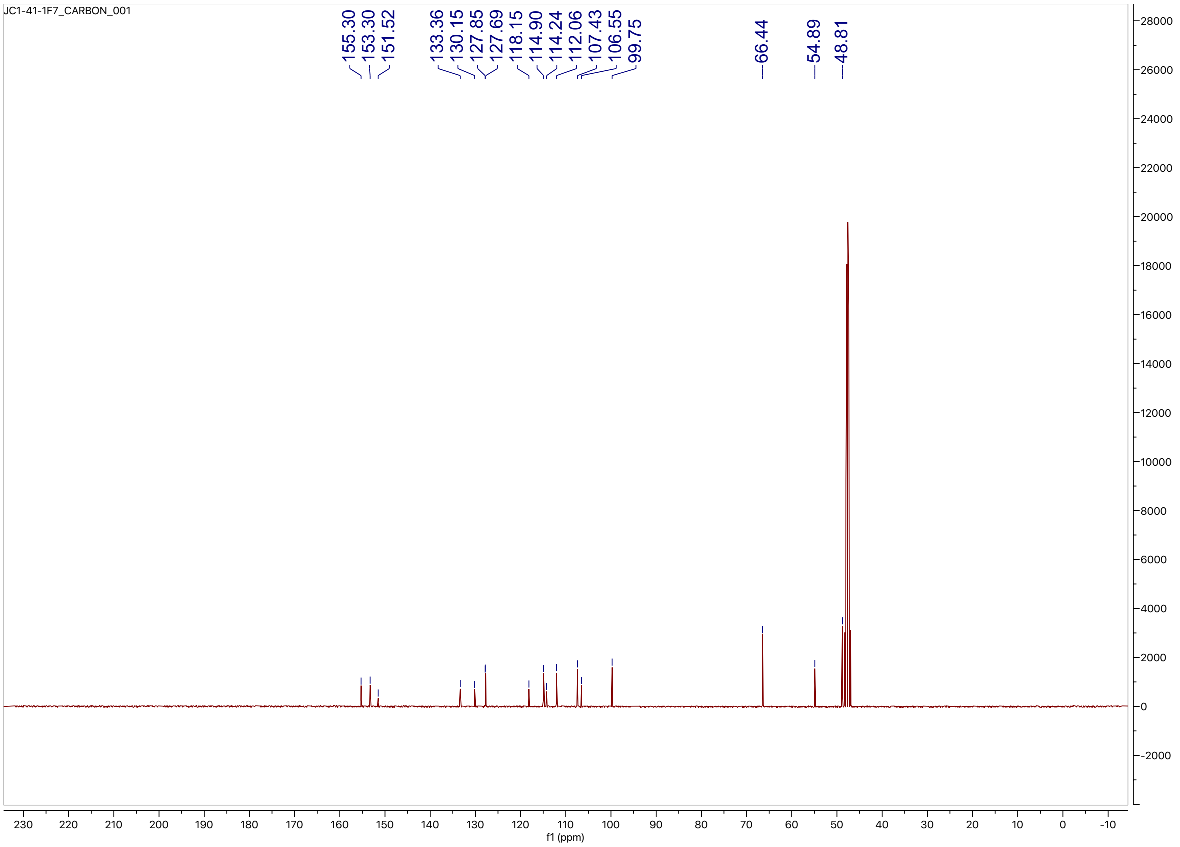

HPLC trace for Compound **5**

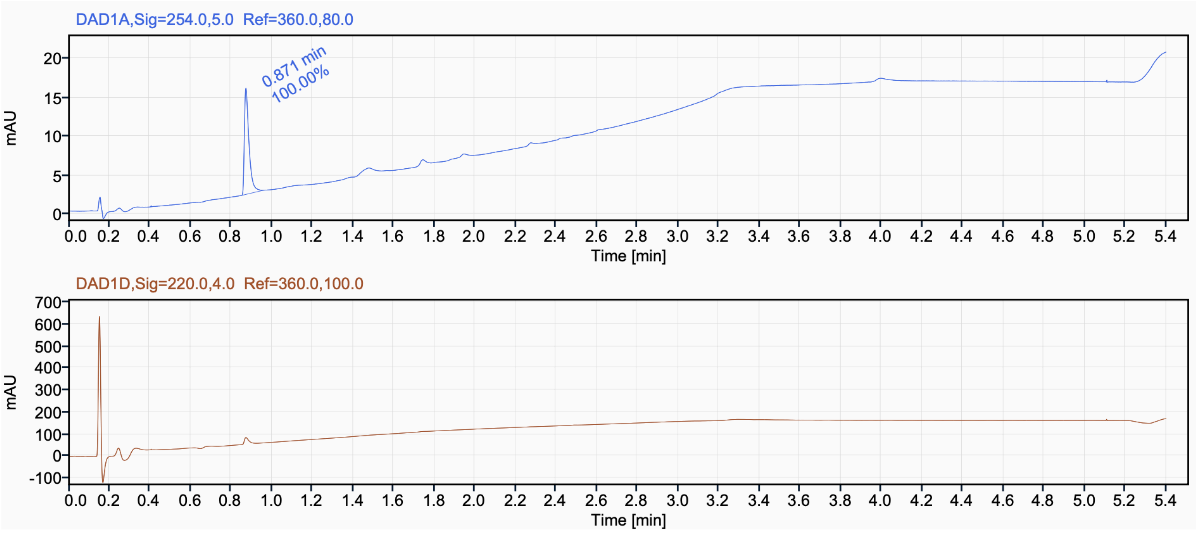

$^{1}H$ NMR (400 MHz, $CD_3OD$) for Compound **6**

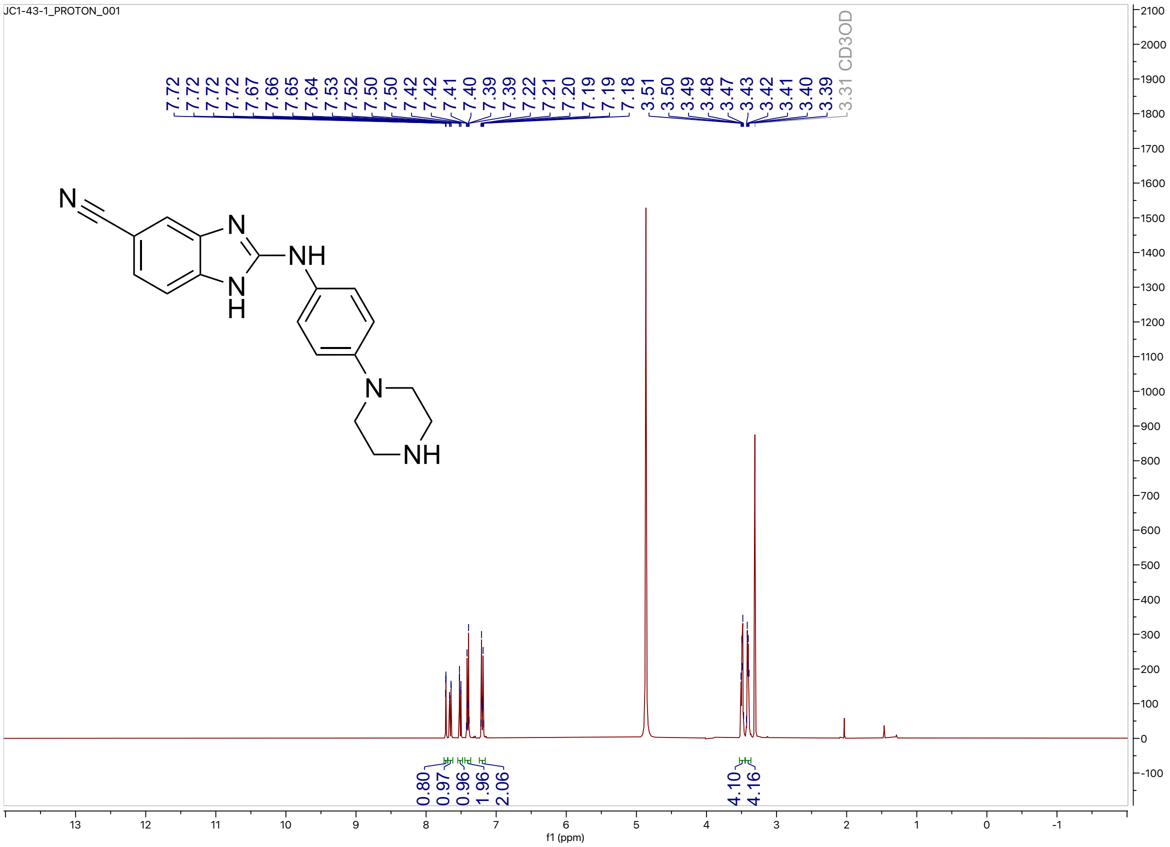


$^{13}C$ NMR (100 MHz, $CD_3OD$) for Compound **6**

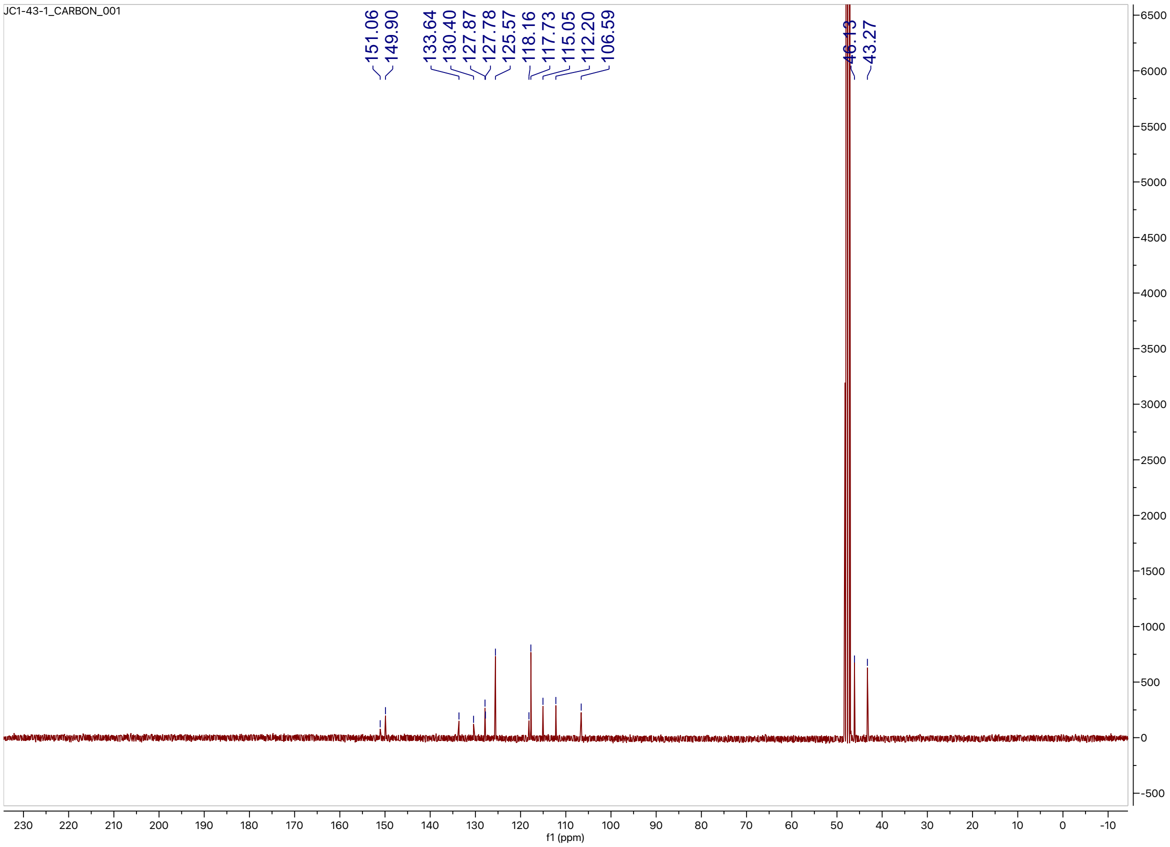

HPLC trace for compound **6**

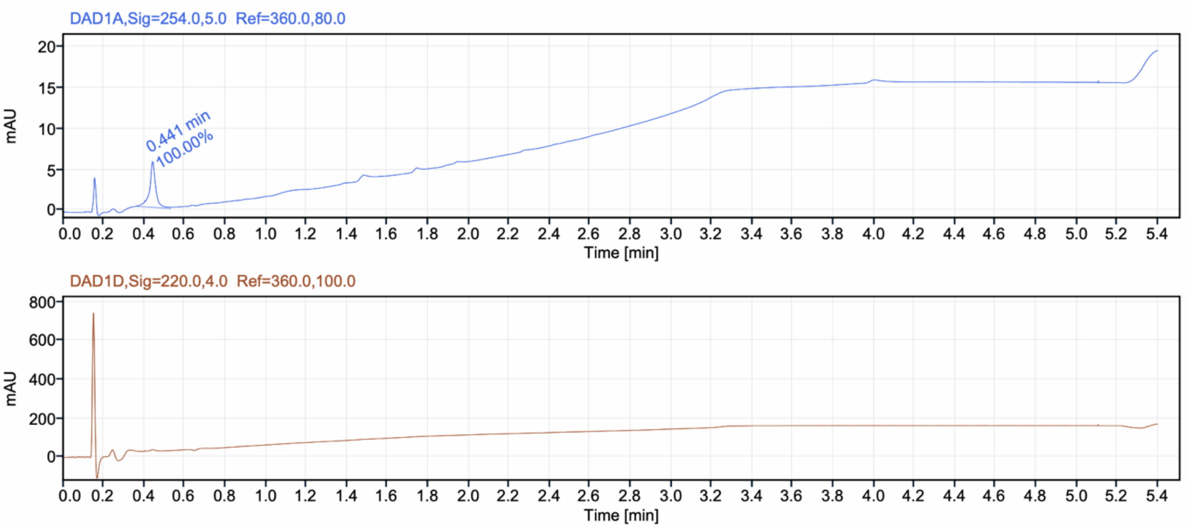

$^1H$ NMR (400 MHz, DMSO-$d_6$) for Compound **7**

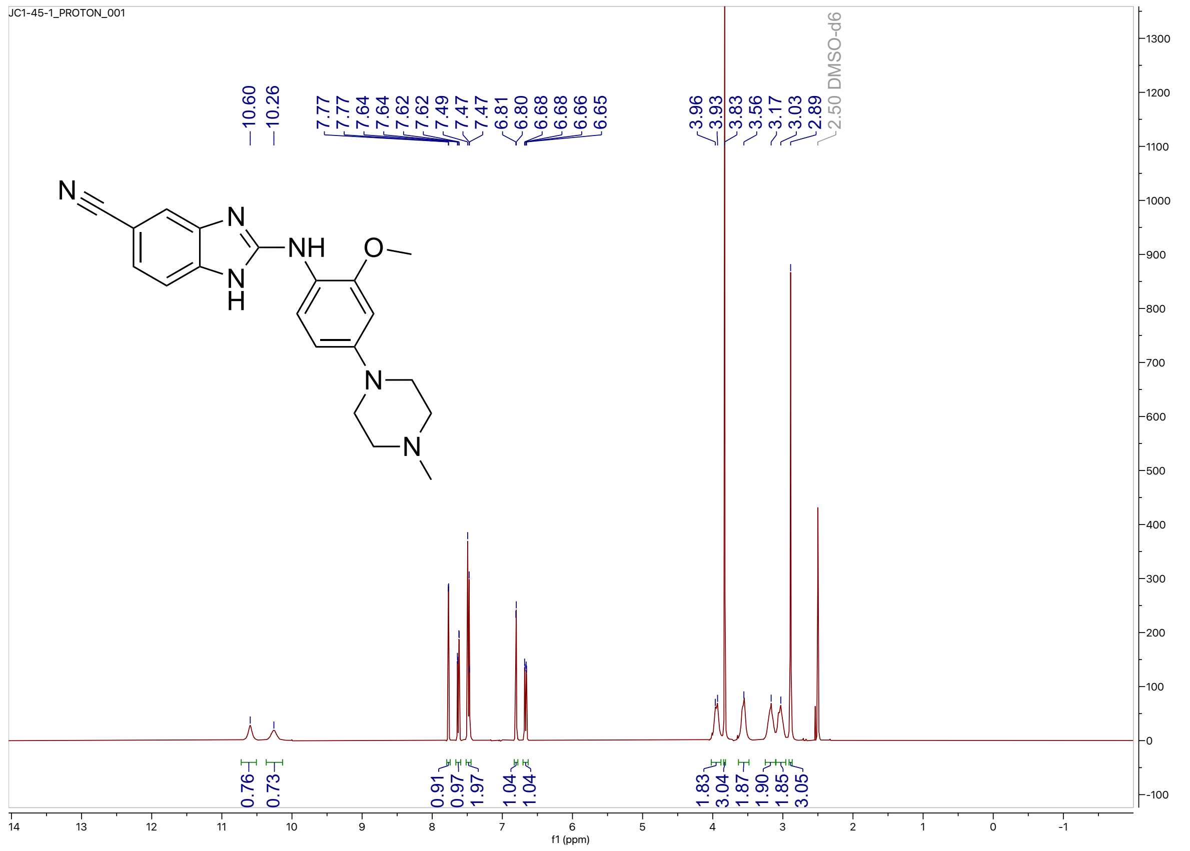

$^{13}C$ NMR (100 MHz, DMSO-$d_6$) for compound 7

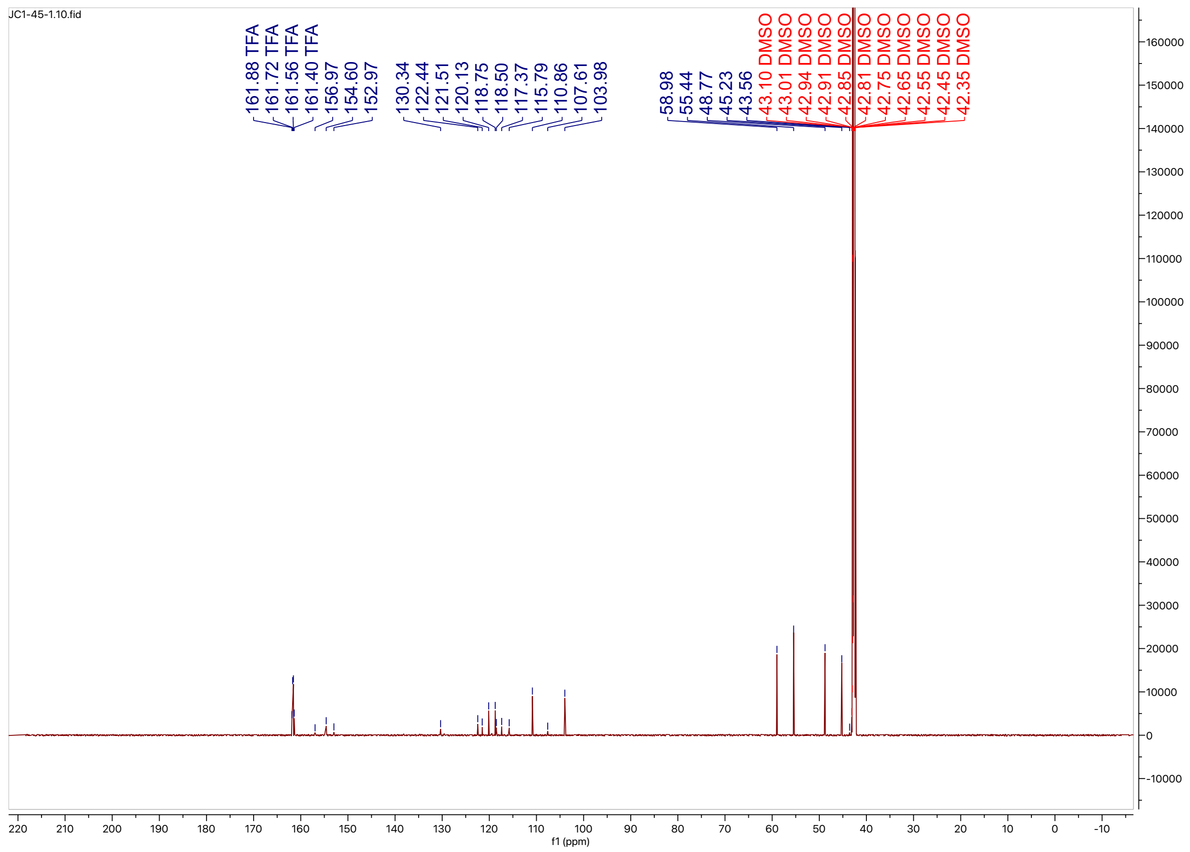

HPLC trace for compound **7**

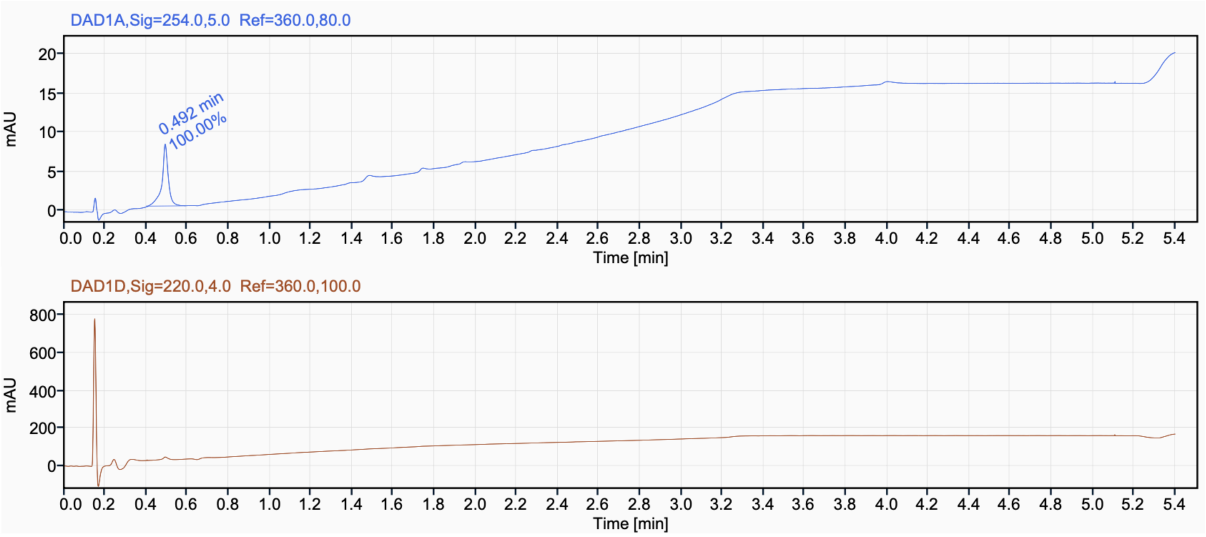

[1]H NMR (400 MHz, DMSO-$d_6$) for Compound **8**

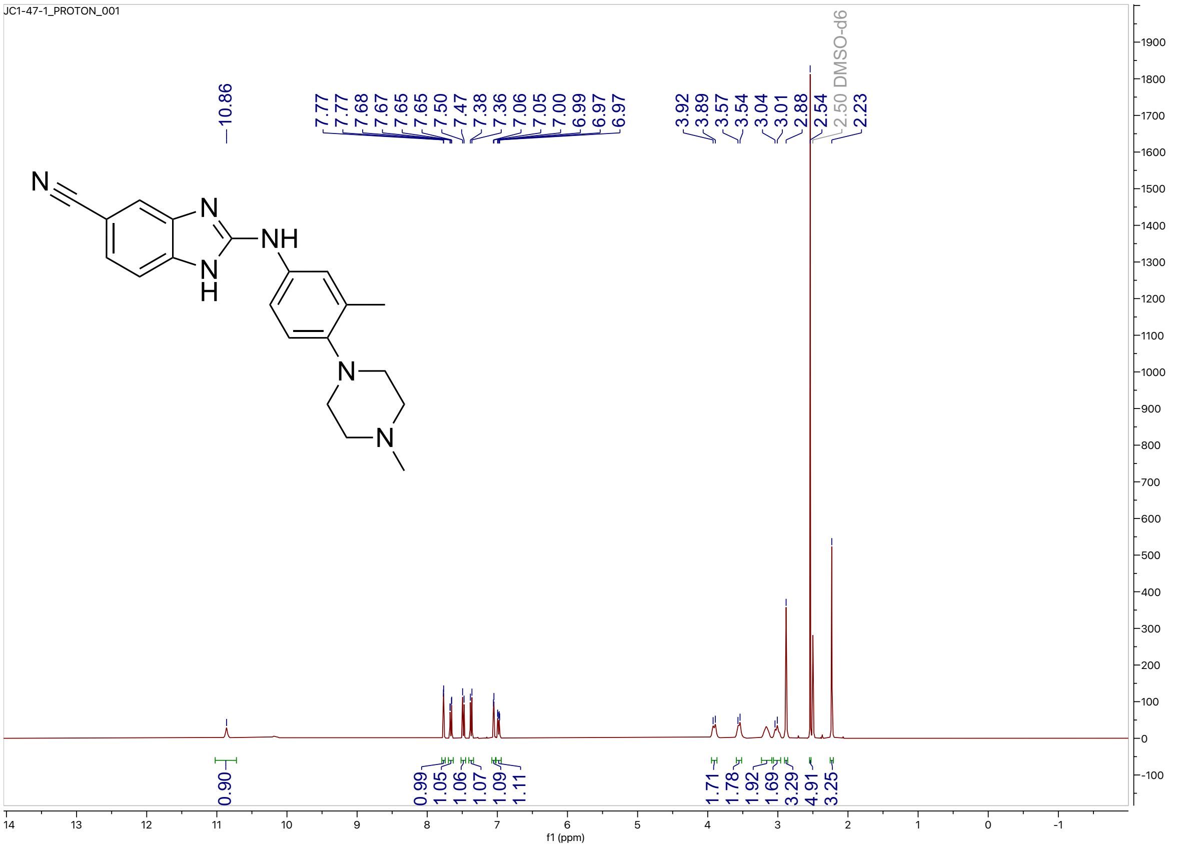


[13]C NMR (100 MHz, DMSO-$d_6$) for compound **8**

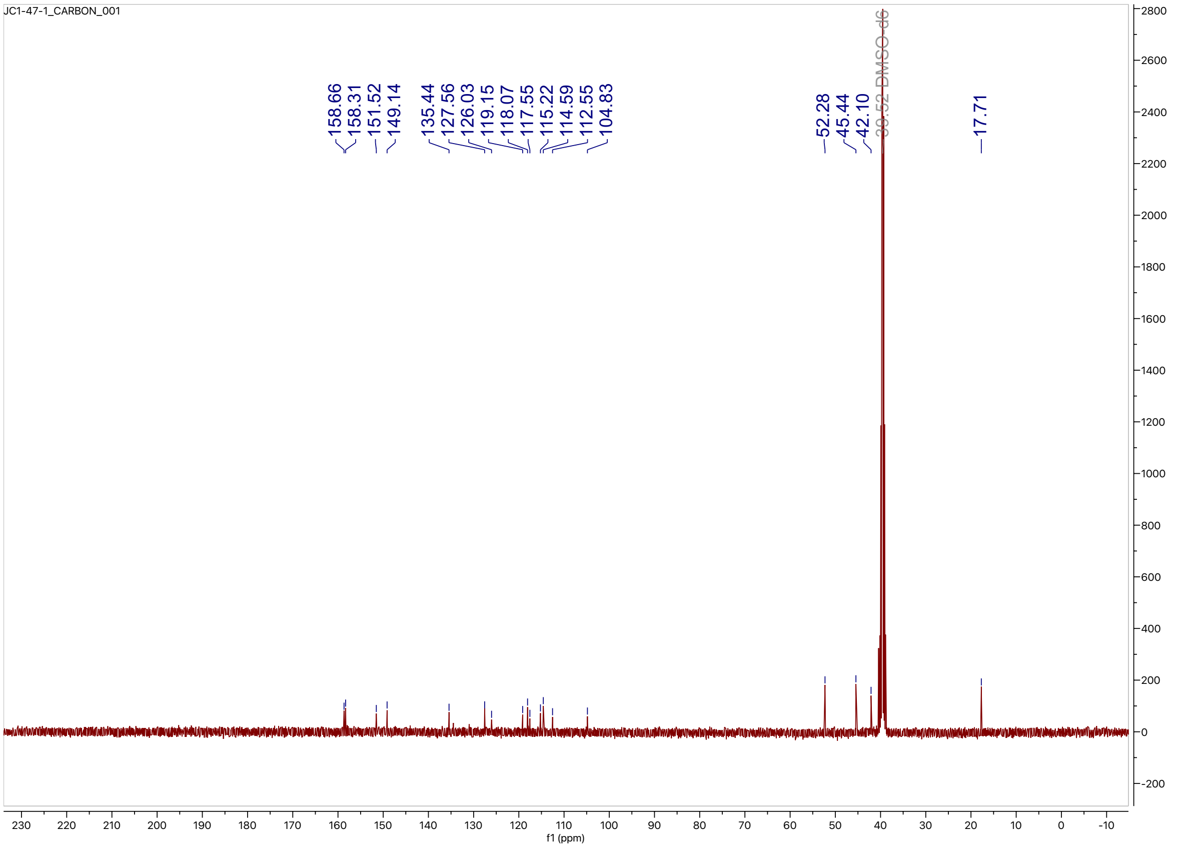

HPLC trace for compound **8**

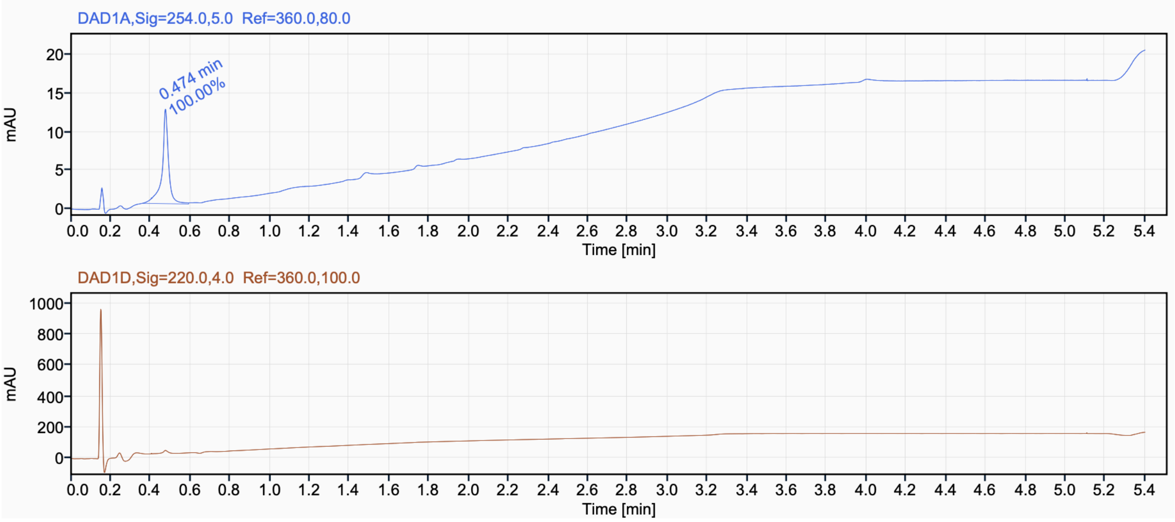

$^{1}H$ NMR (400 MHz, $CD_3OD$) for Compound **9**

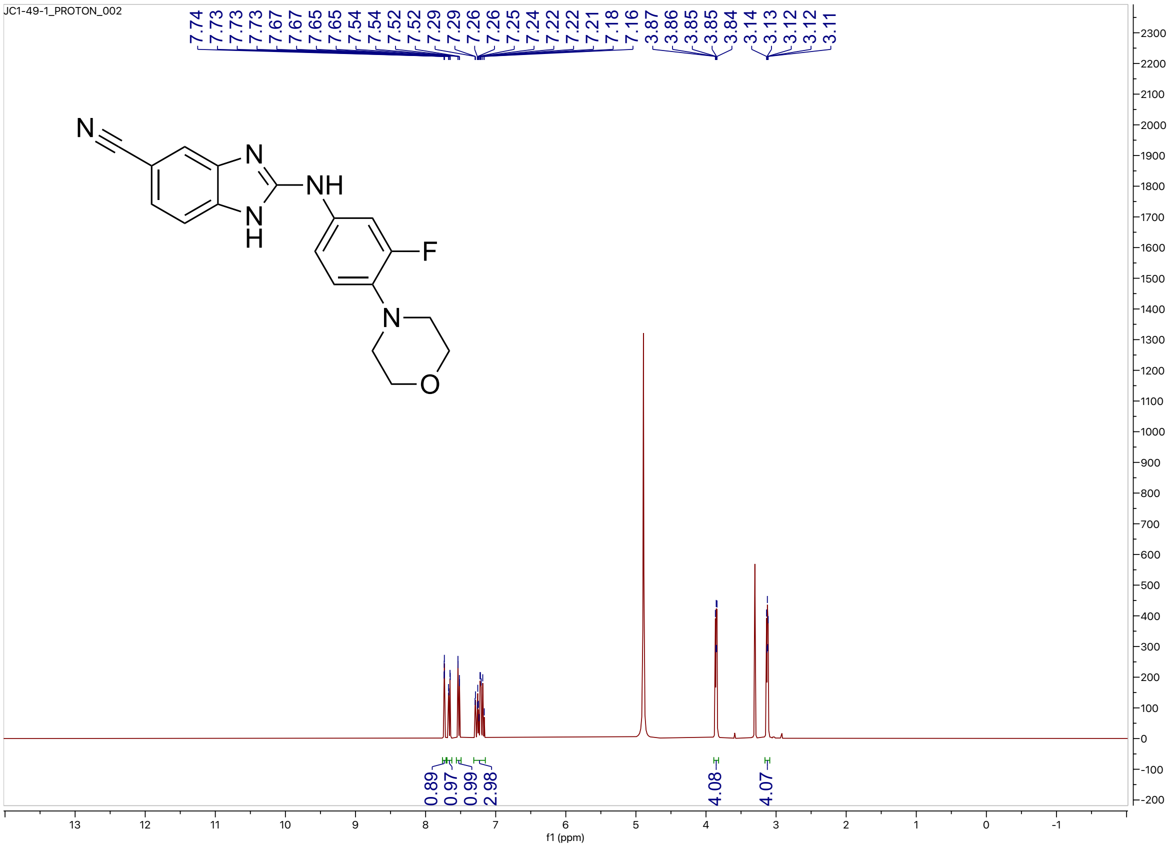


$^{13}C$ NMR (100 MHz, $CD_3OD$) for compound **9**

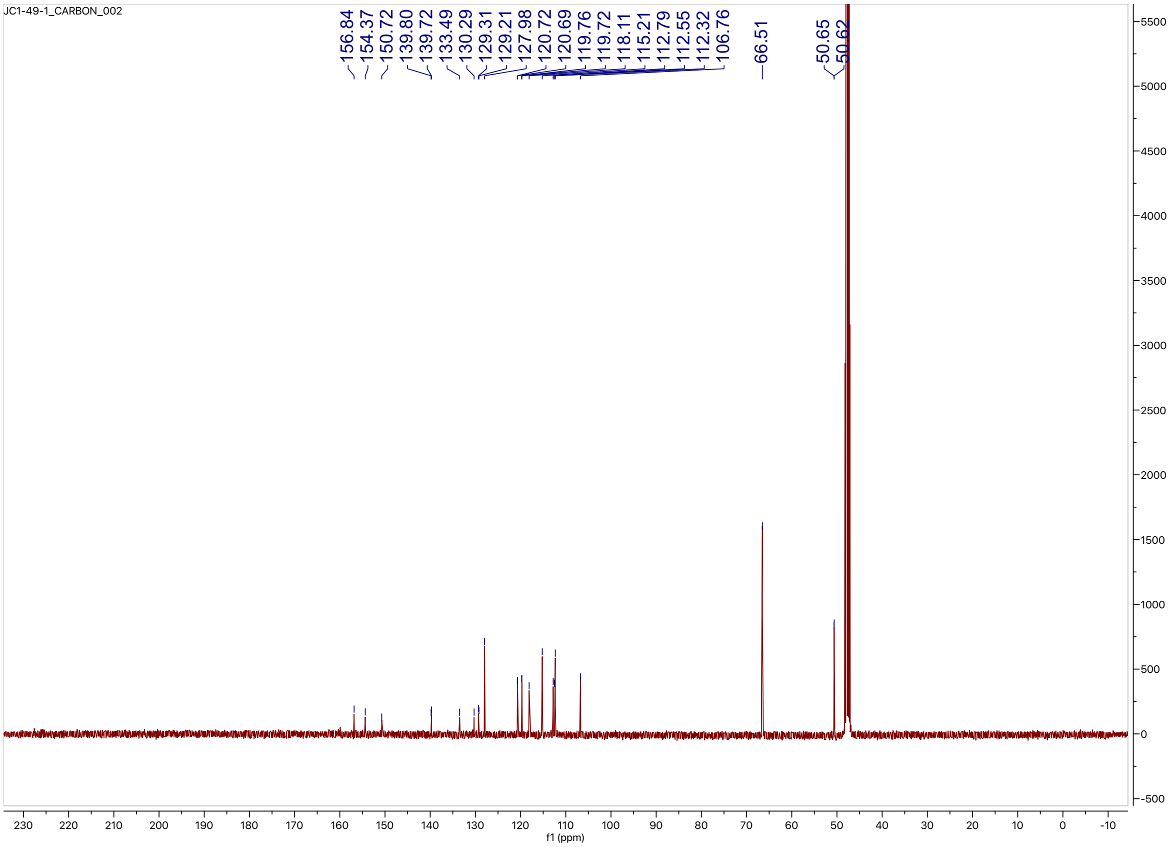

HPLC trace for compound **9**

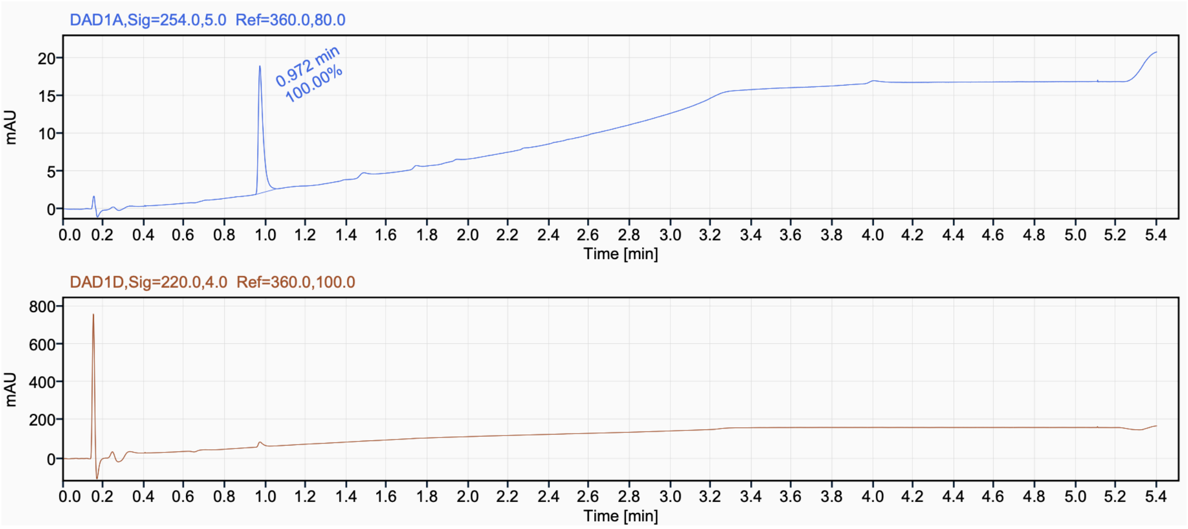

$^{1}H$ NMR (400 MHz, $CD_3OD$) for Compound **10**

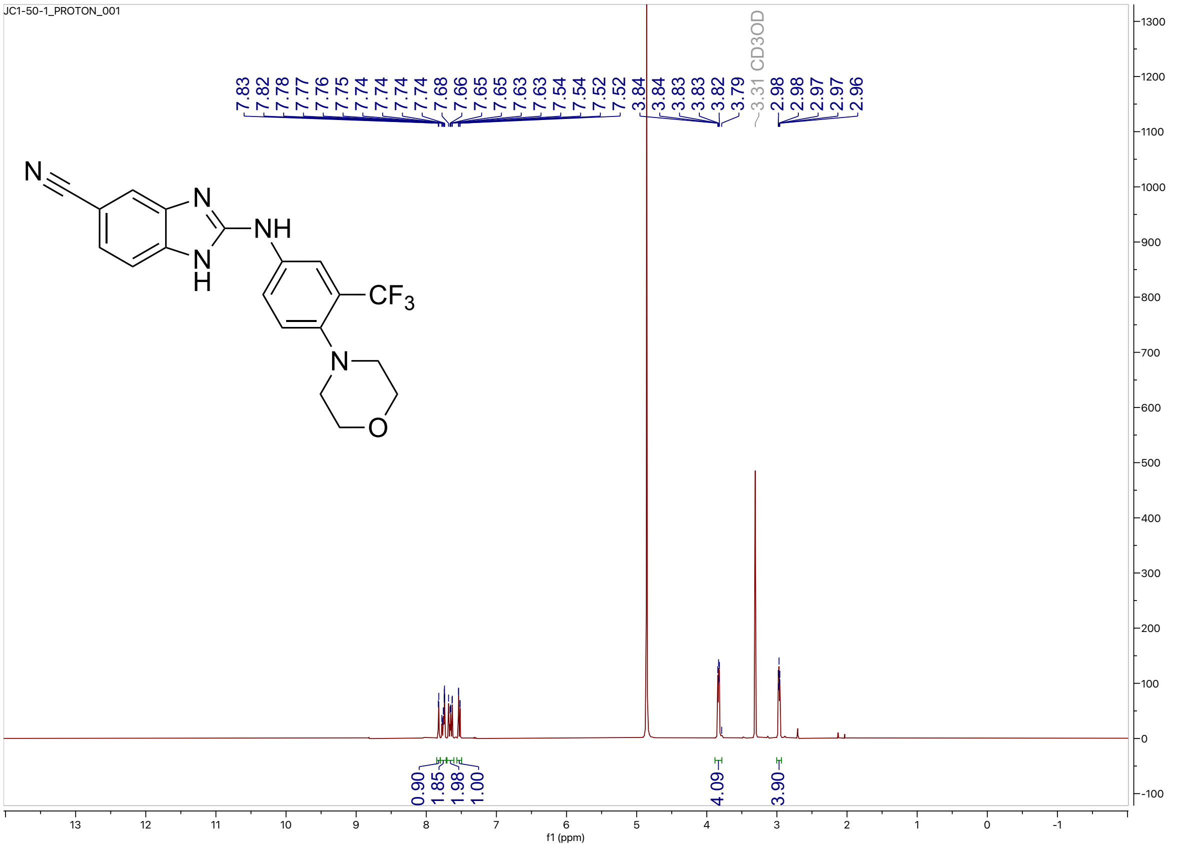


$^{13}C$ NMR (100 MHz, $CD_3OD$) for compound **10**

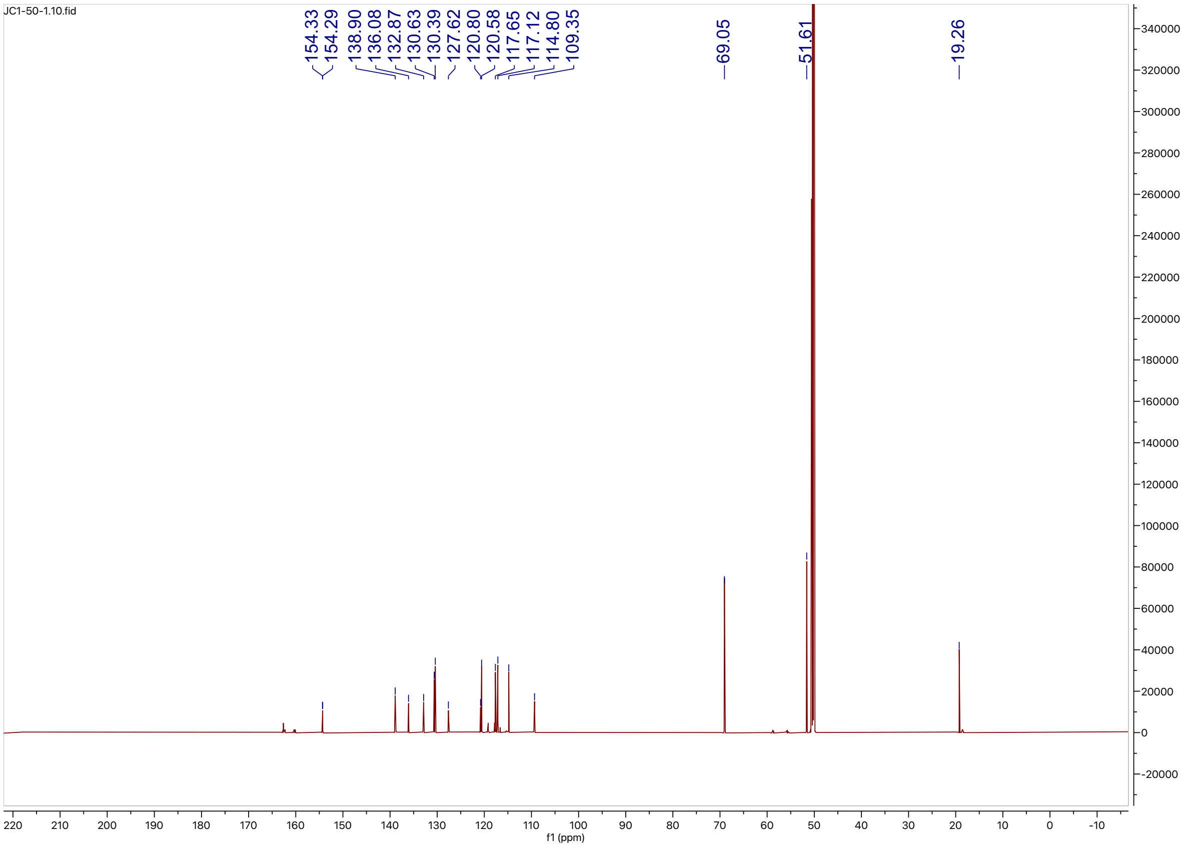

HPLC trace for compound **10**

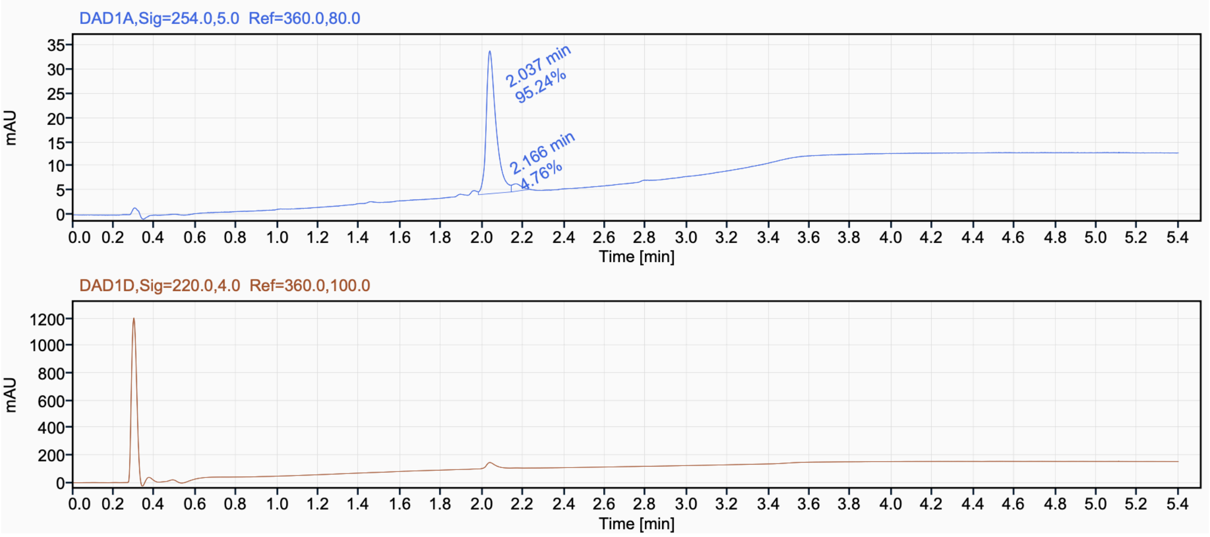

$^{1}H$ NMR (400 MHz, $CD_3OD$) for compound **11**

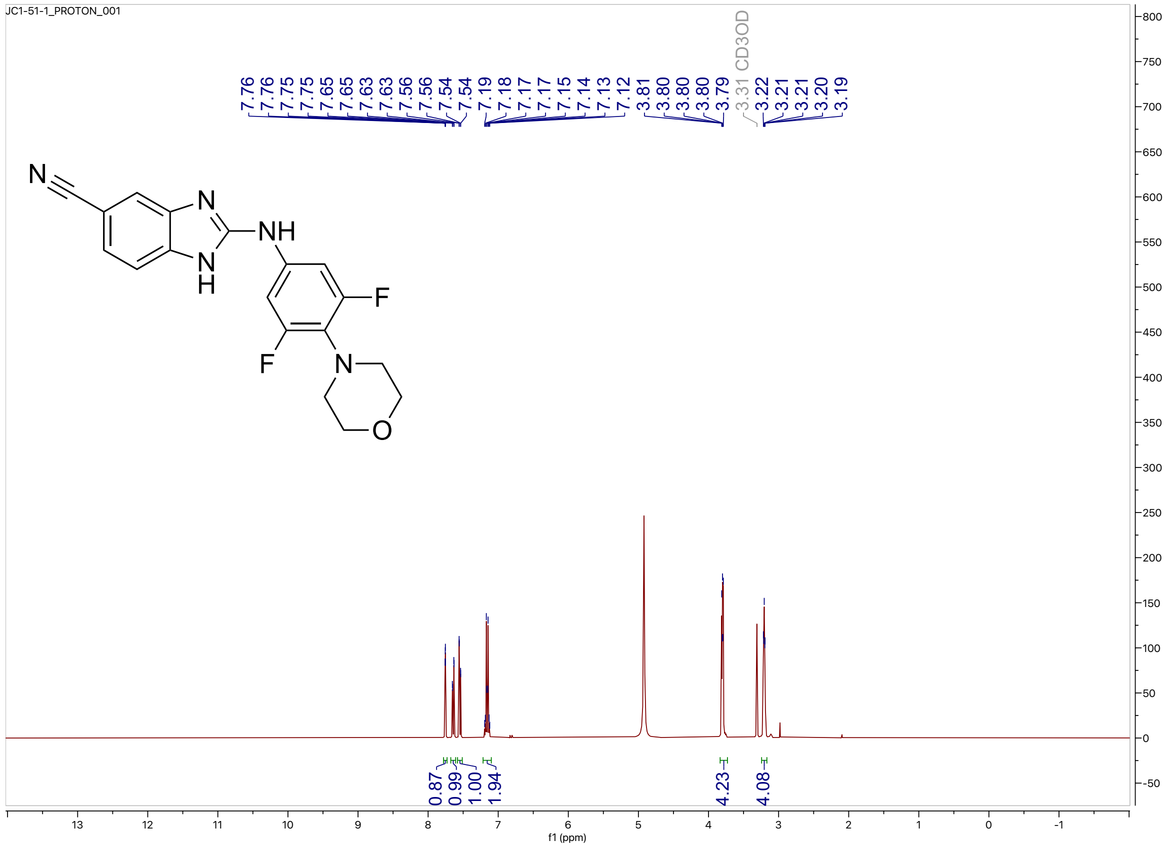


$^{13}C$ NMR (100 MHz, $CD_3OD$) for compound **11**

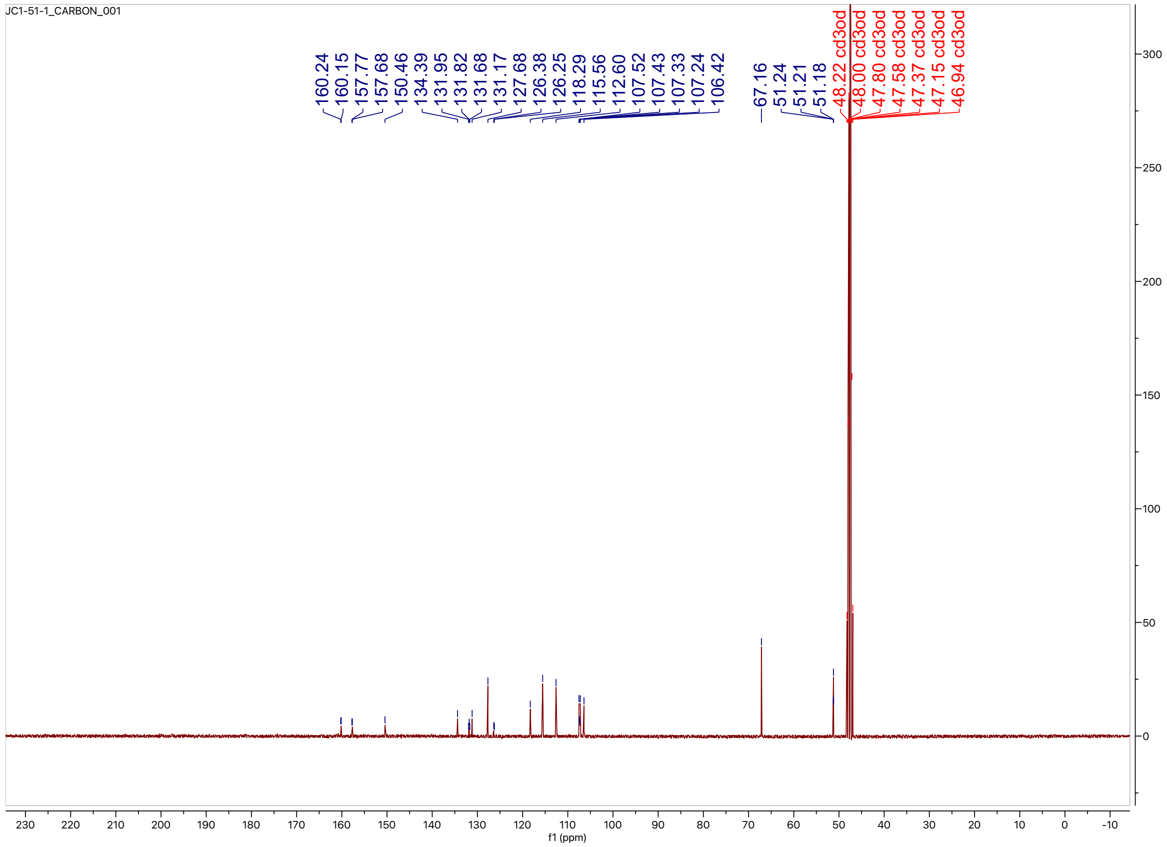

HPLC tracer for compound **11**

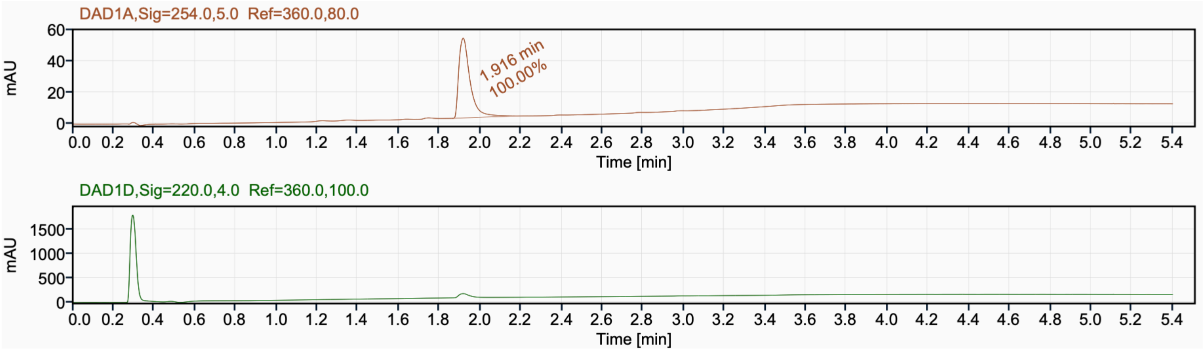

$^{1}H$ NMR (400 MHz, $CD_3OD$) for compound **12**

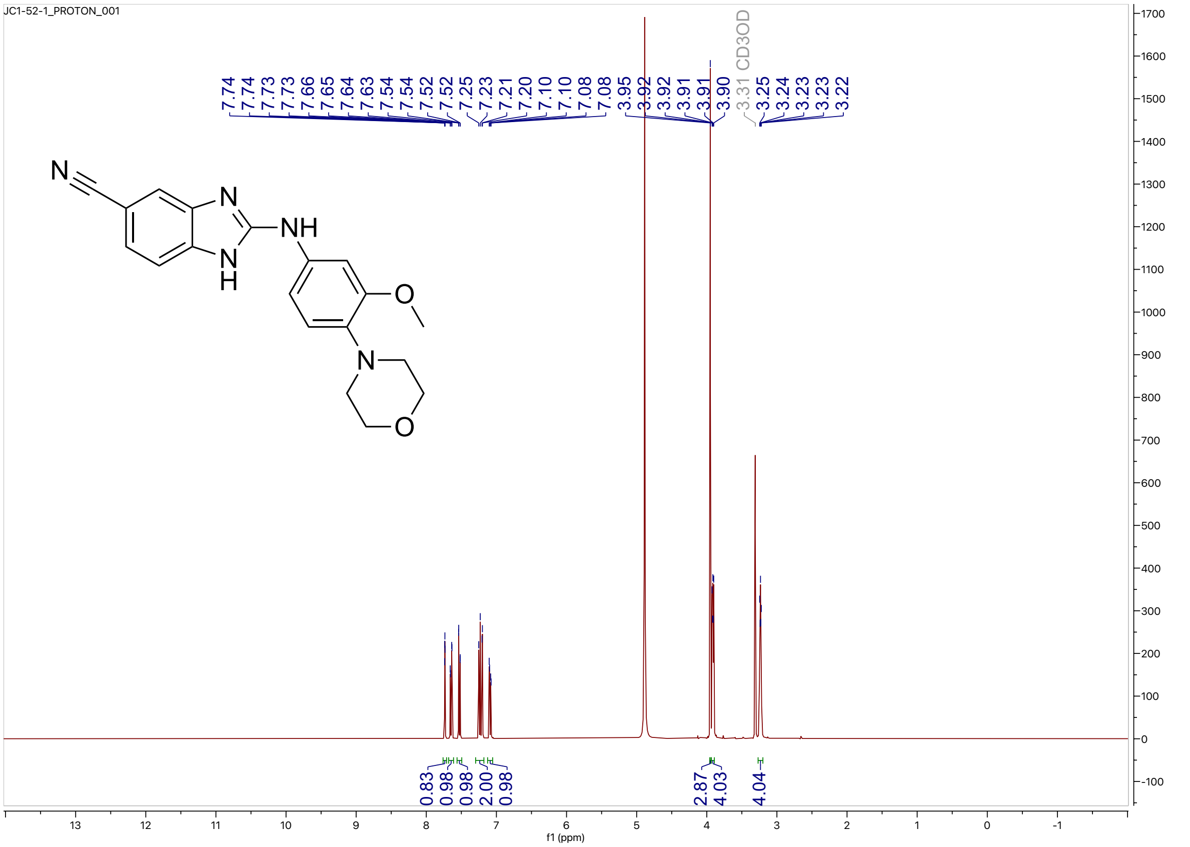


$^{13}C$ NMR (126 MHz, $CD_3OD$) for compound **12**

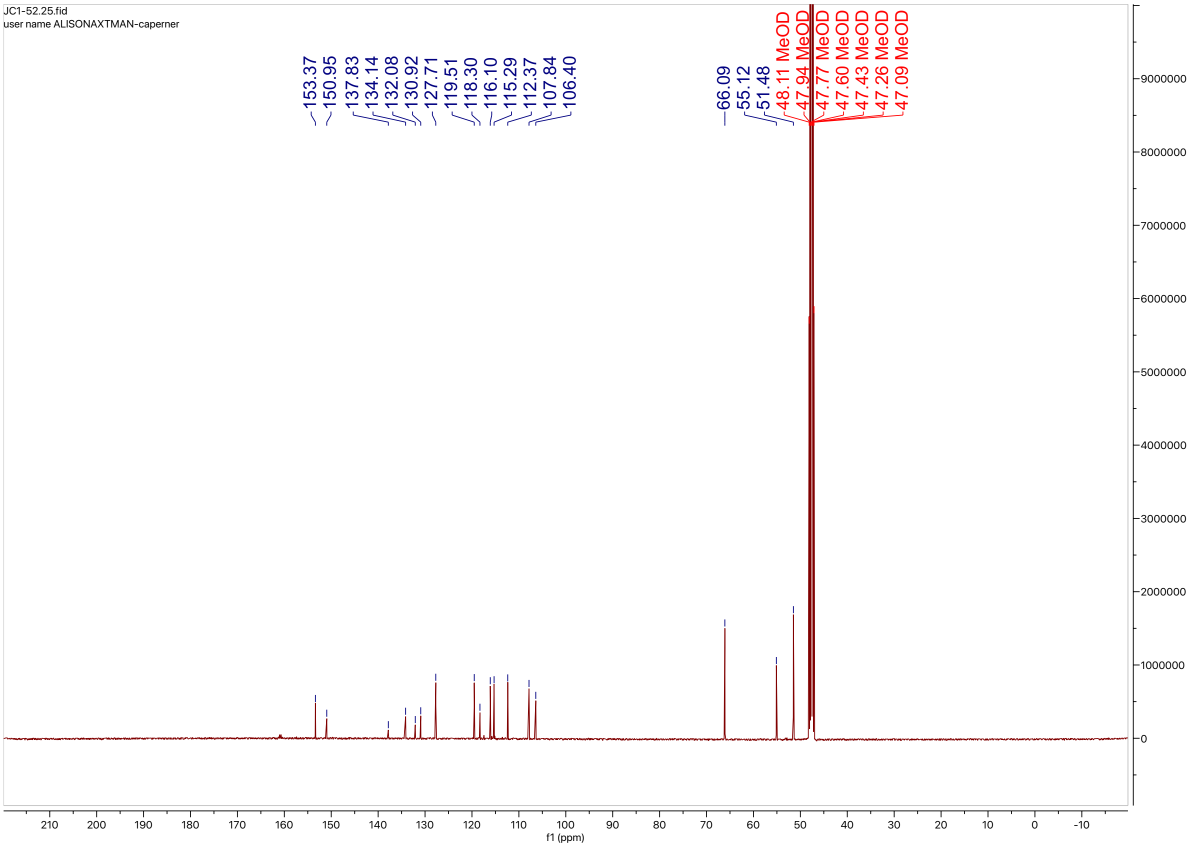

HPLC trace for compound **12**

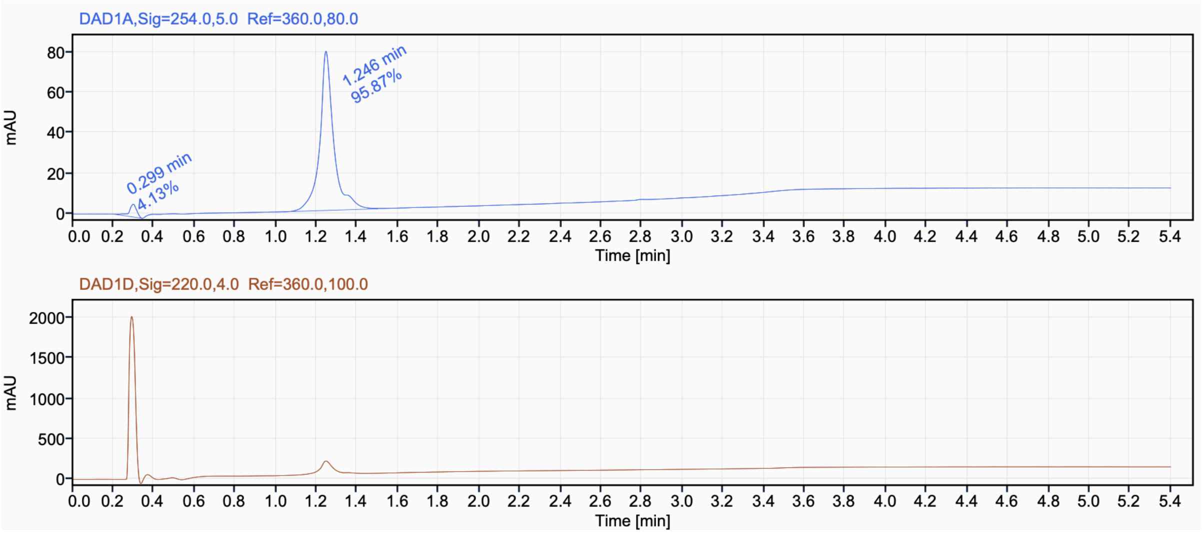

$^{1}H$ NMR (400 MHz, $CD_3OD$) for compound **13**

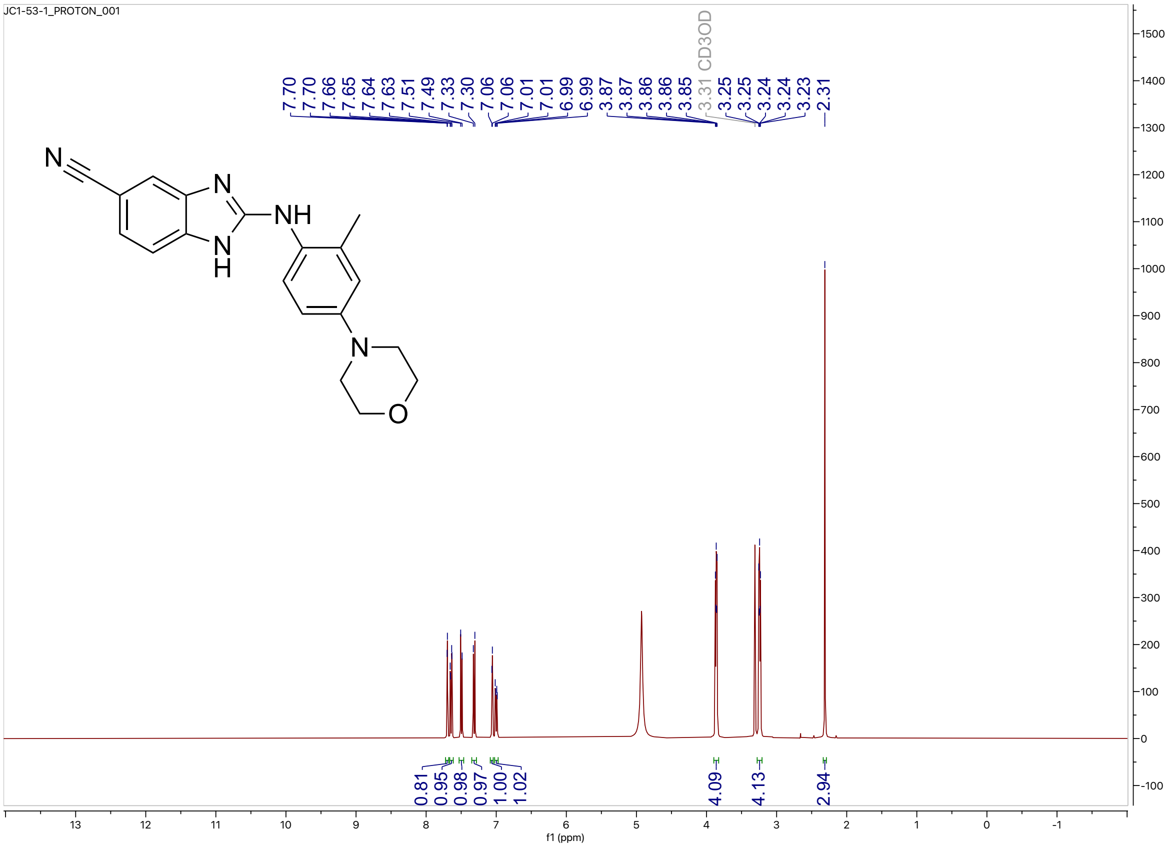


$^{13}C$ NMR (100 MHz, $CD_3OD$) for compound **13**

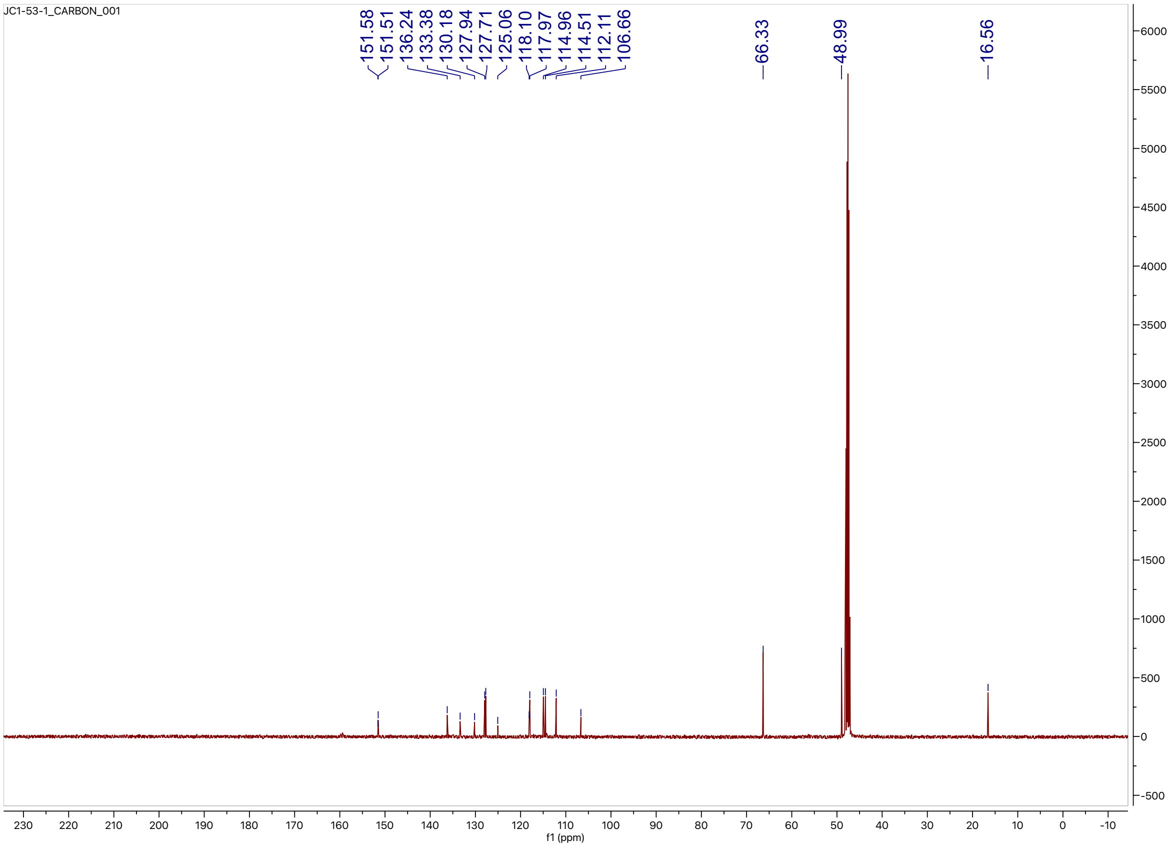

HPLC trace for compound 13

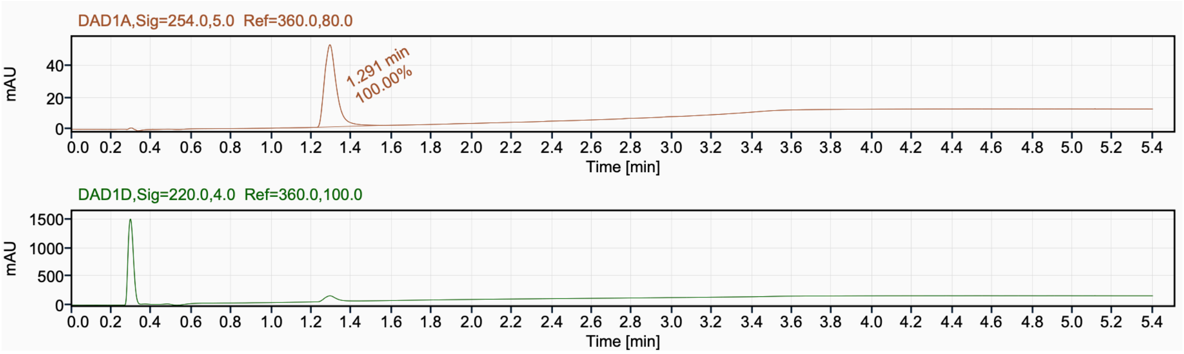

$^{1}H$ NMR (400 MHz, $CD_3OD$) for compound **14**

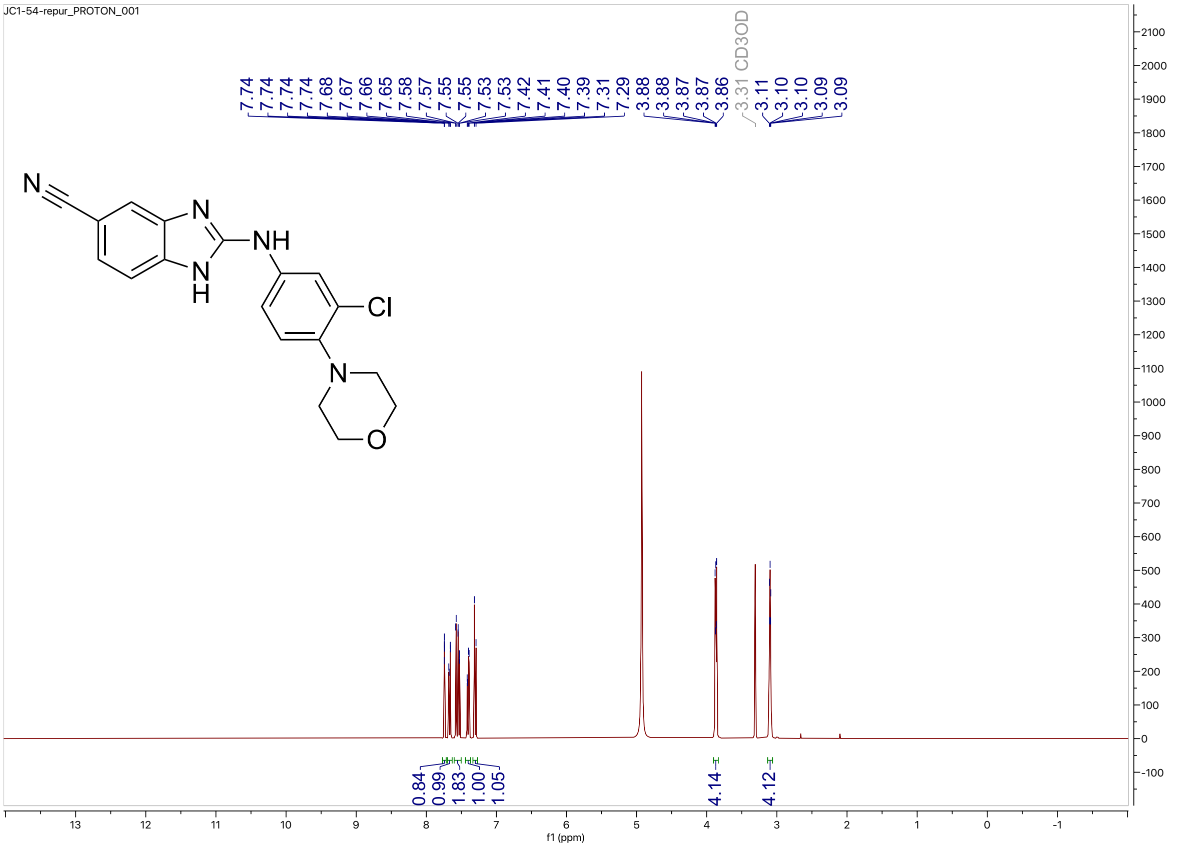


$^{13}C$ NMR (100 MHz, $CD_3OD$) for compound **14**

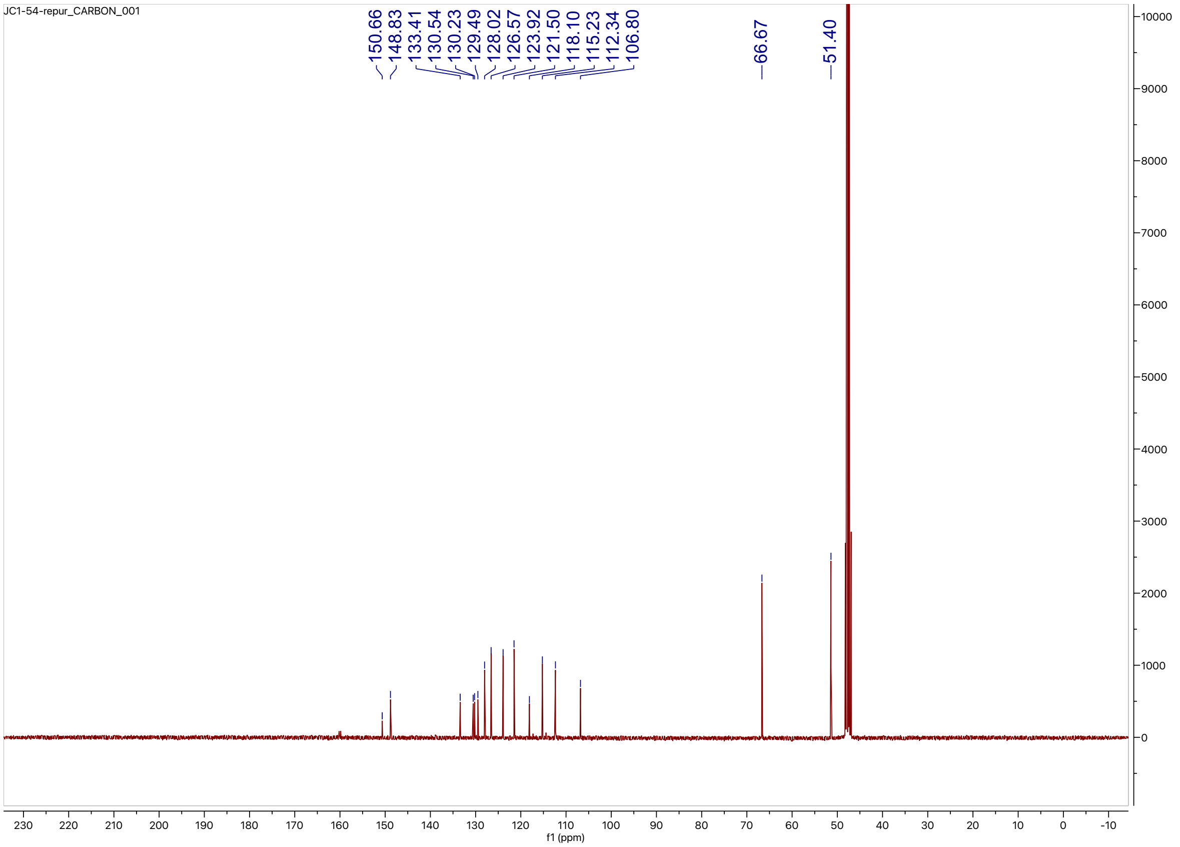

HPLC trace for compound **14**

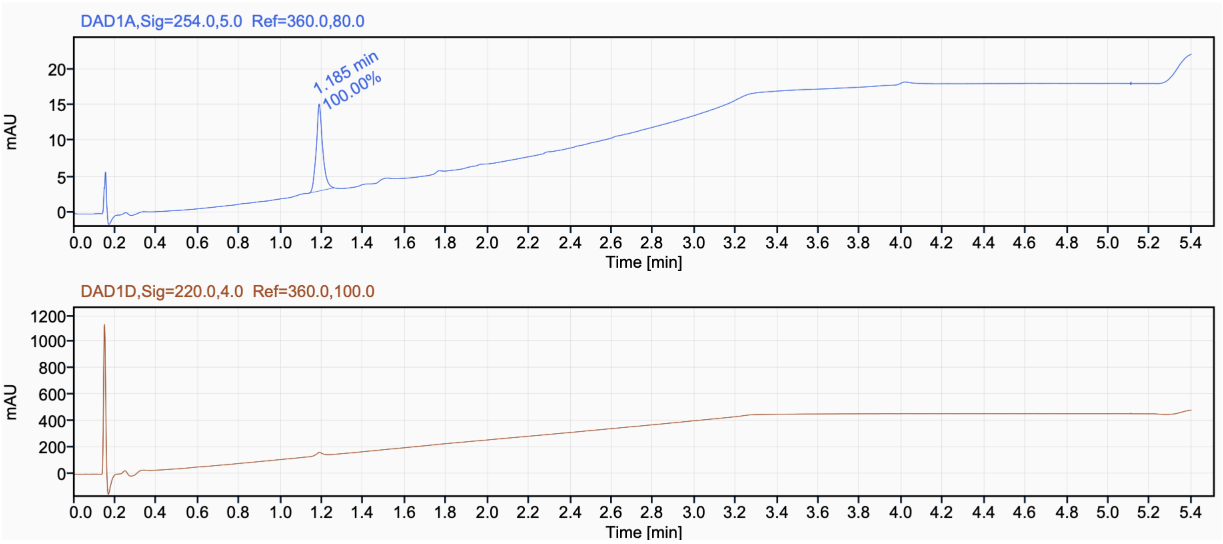

$^{1}H$ NMR (850 MHz, $CD_3OD$) for compound **15**

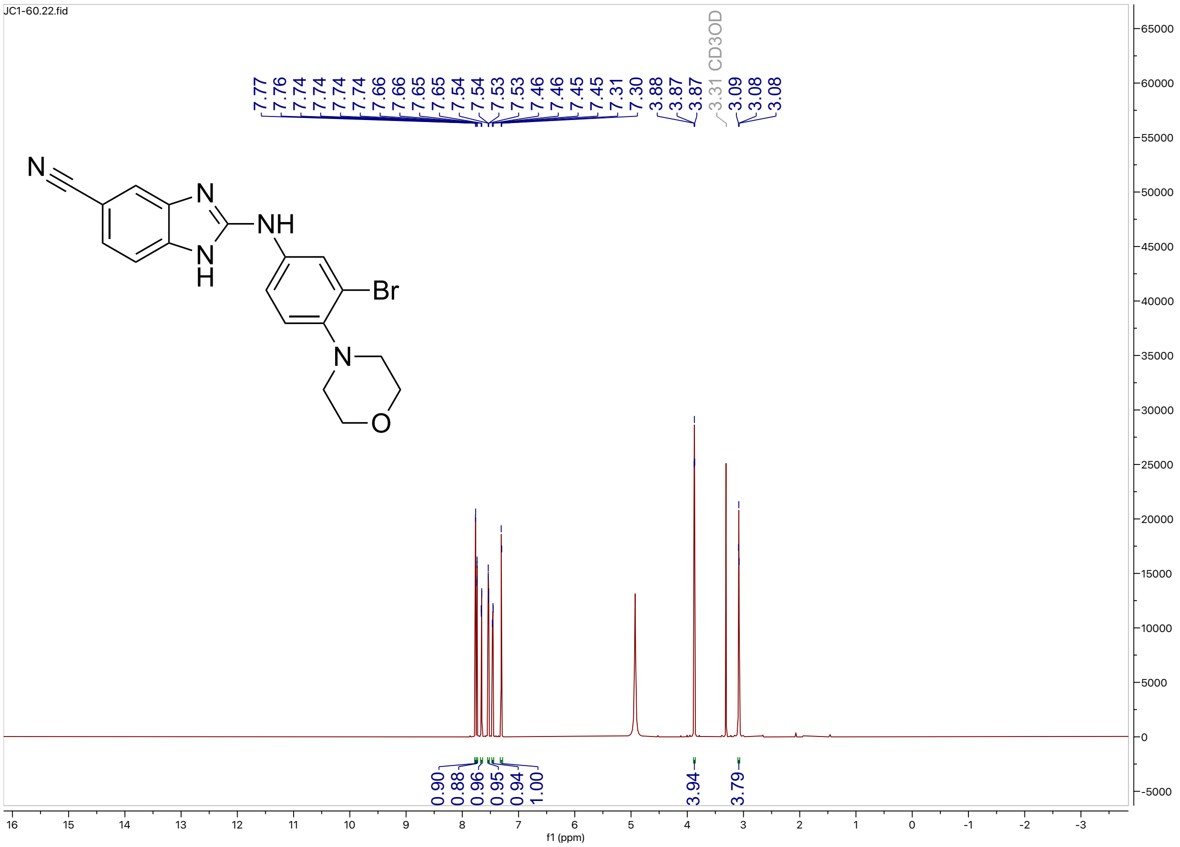


$^{13}C$ NMR (214 MHz, $CD_3OD$) for compound **15**

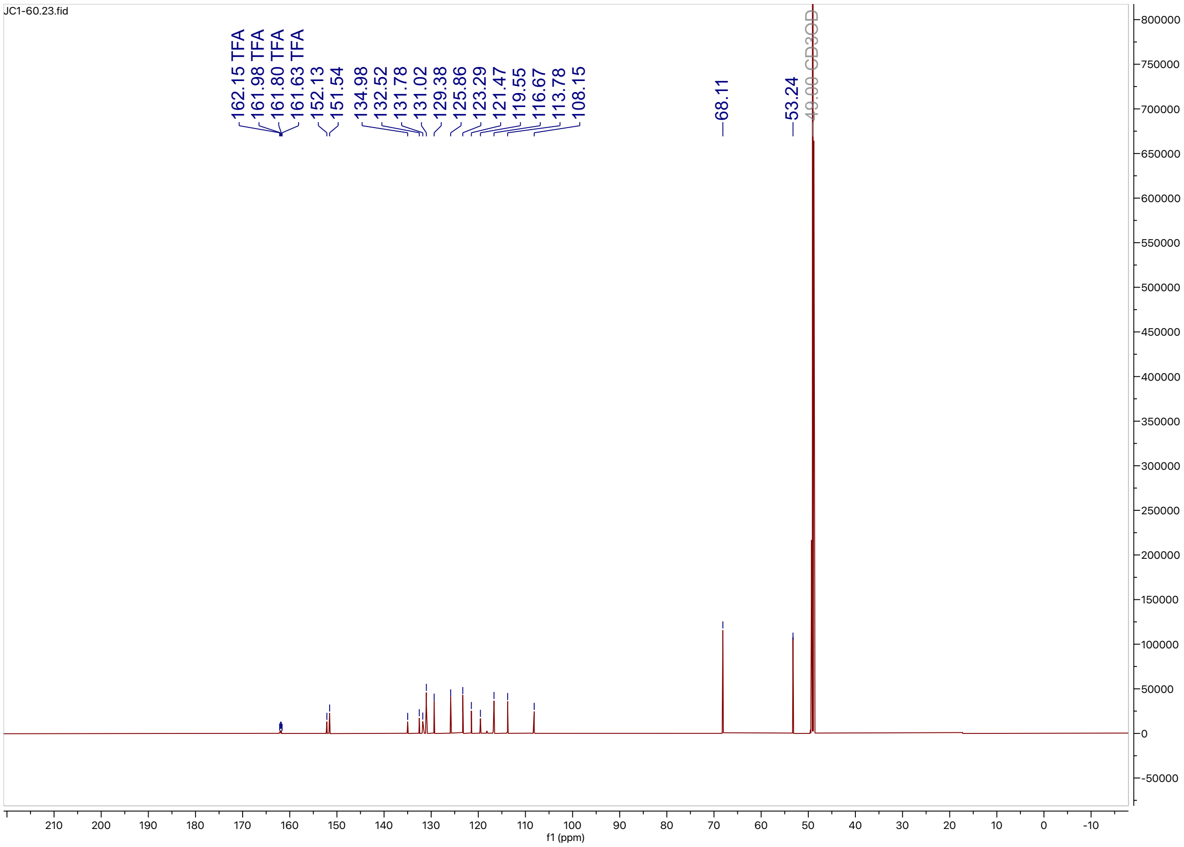

HPLC trace for compound **15**

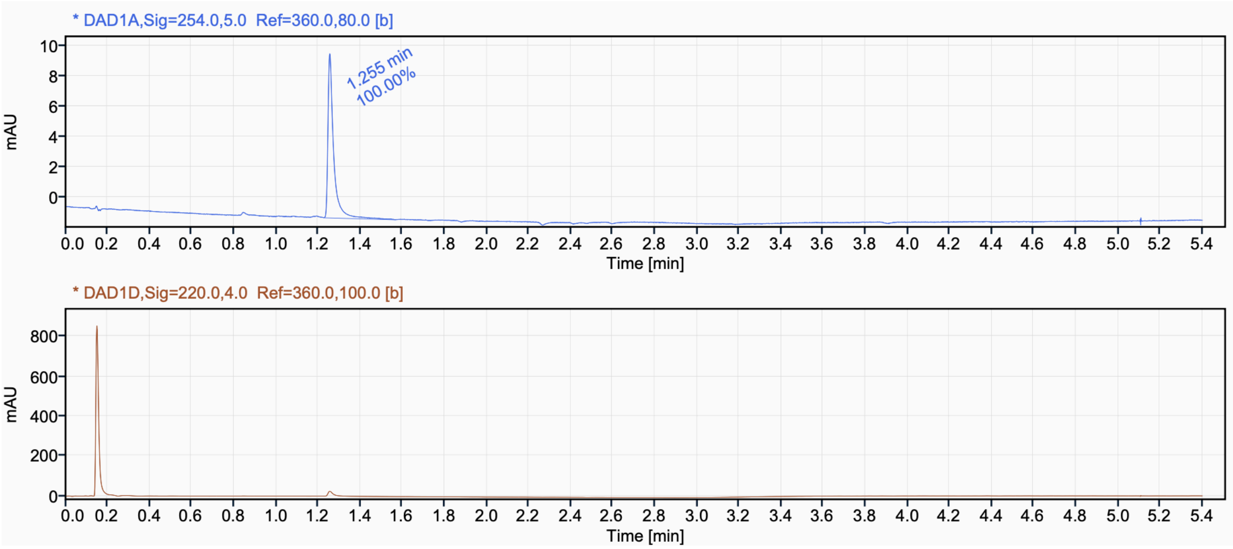

$^{1}H$ NMR (500 MHz, $CD_3OD$) for compound **16**

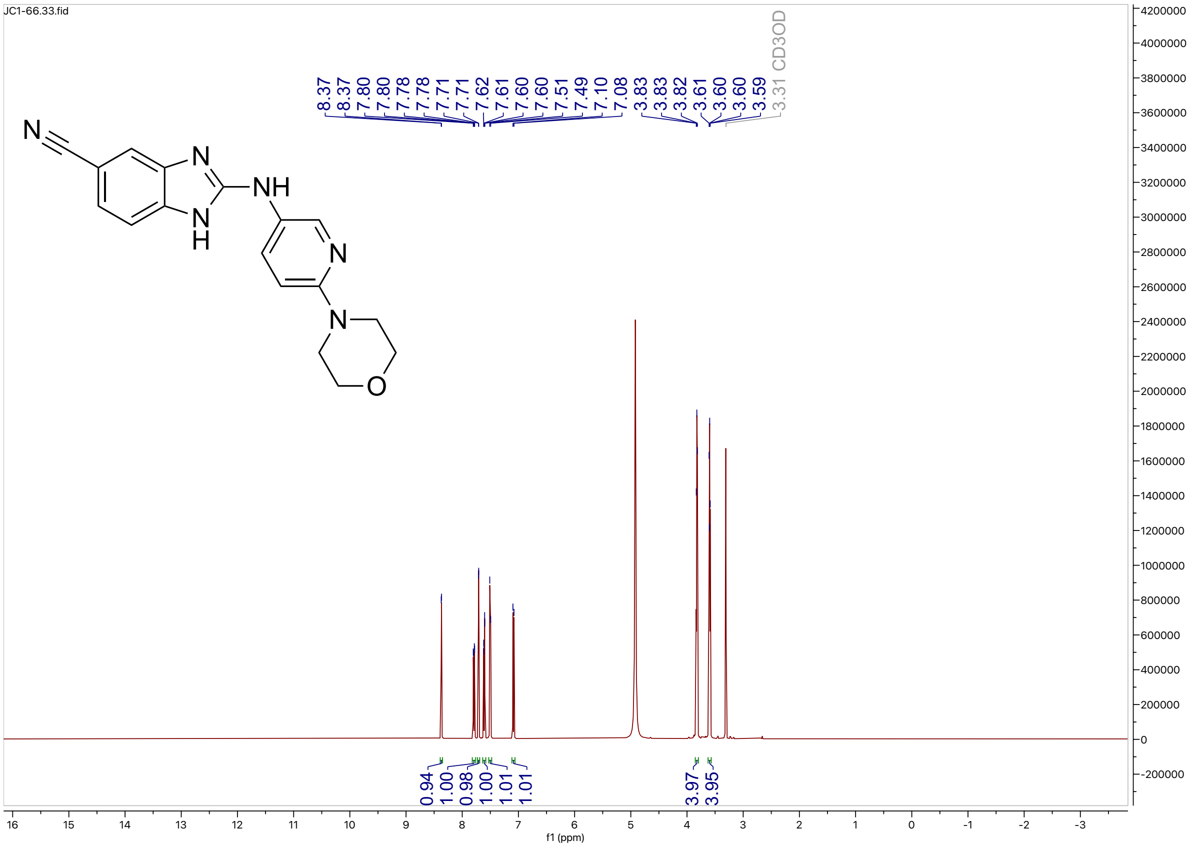


$^{13}C$ NMR (126 MHz, $CD_3OD$) for compound **16**

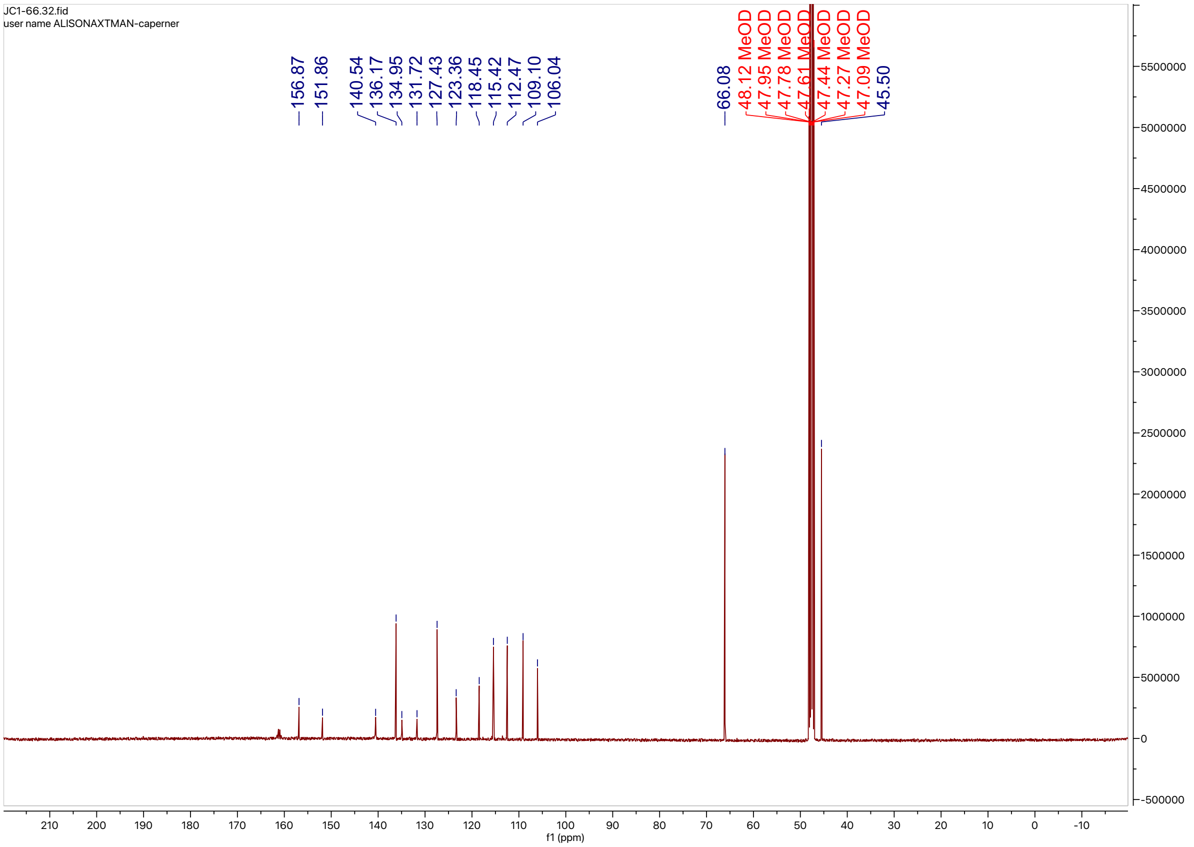

HPLC trace for compound **16**

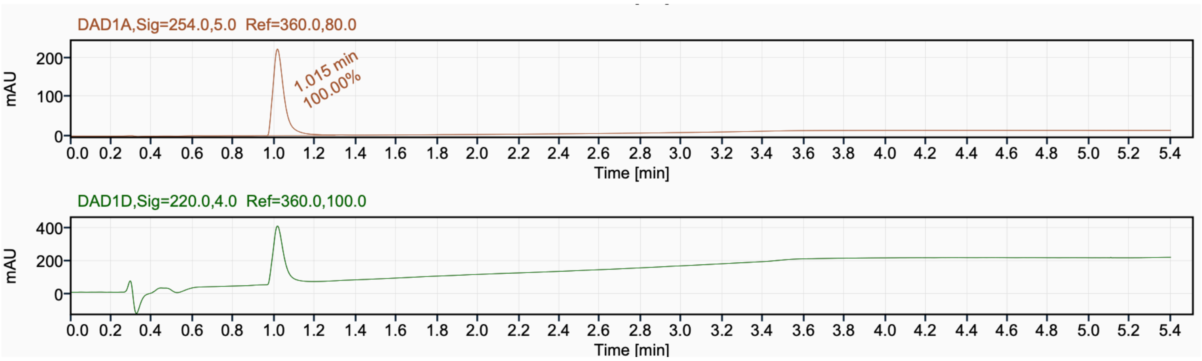

$^{1}H$ NMR (500 MHz, $CD_3OD$) for compound **17**

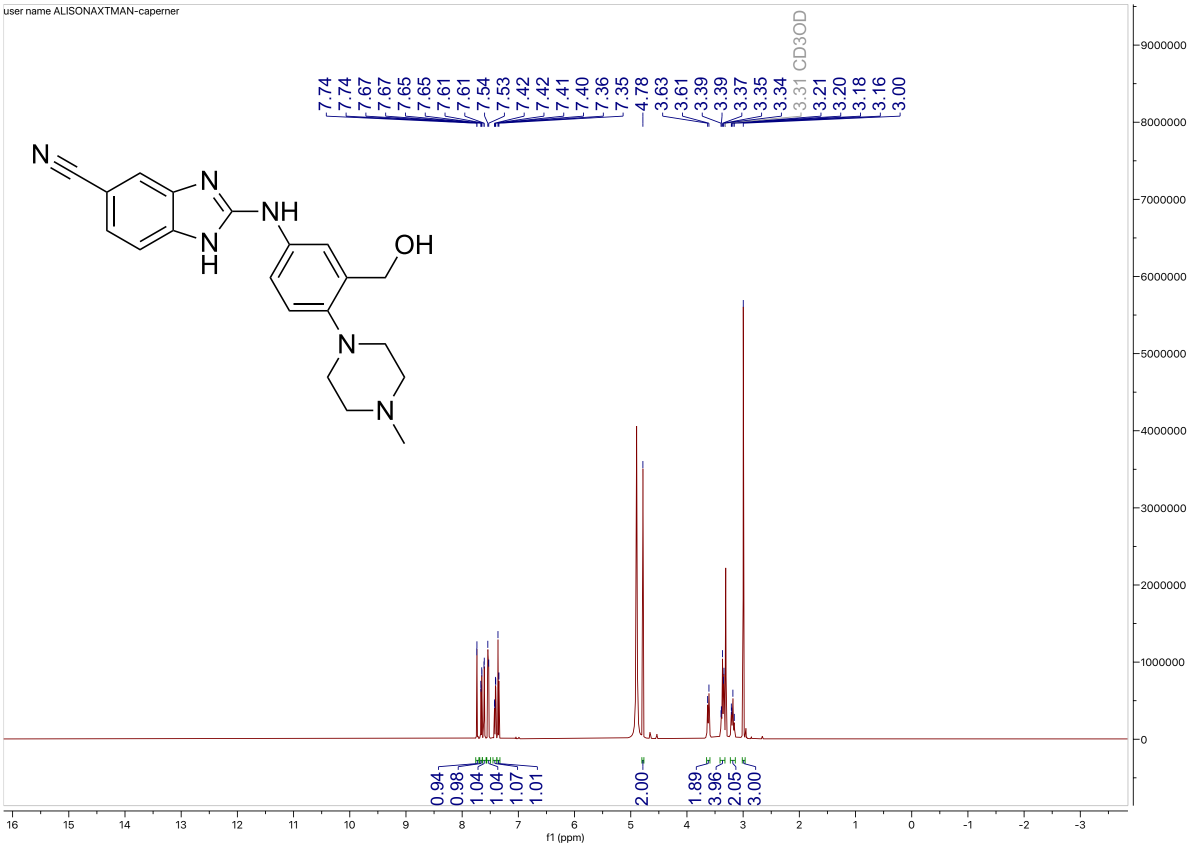


$^{13}C$ NMR (126 MHz, $CD_3OD$) for compound **17**

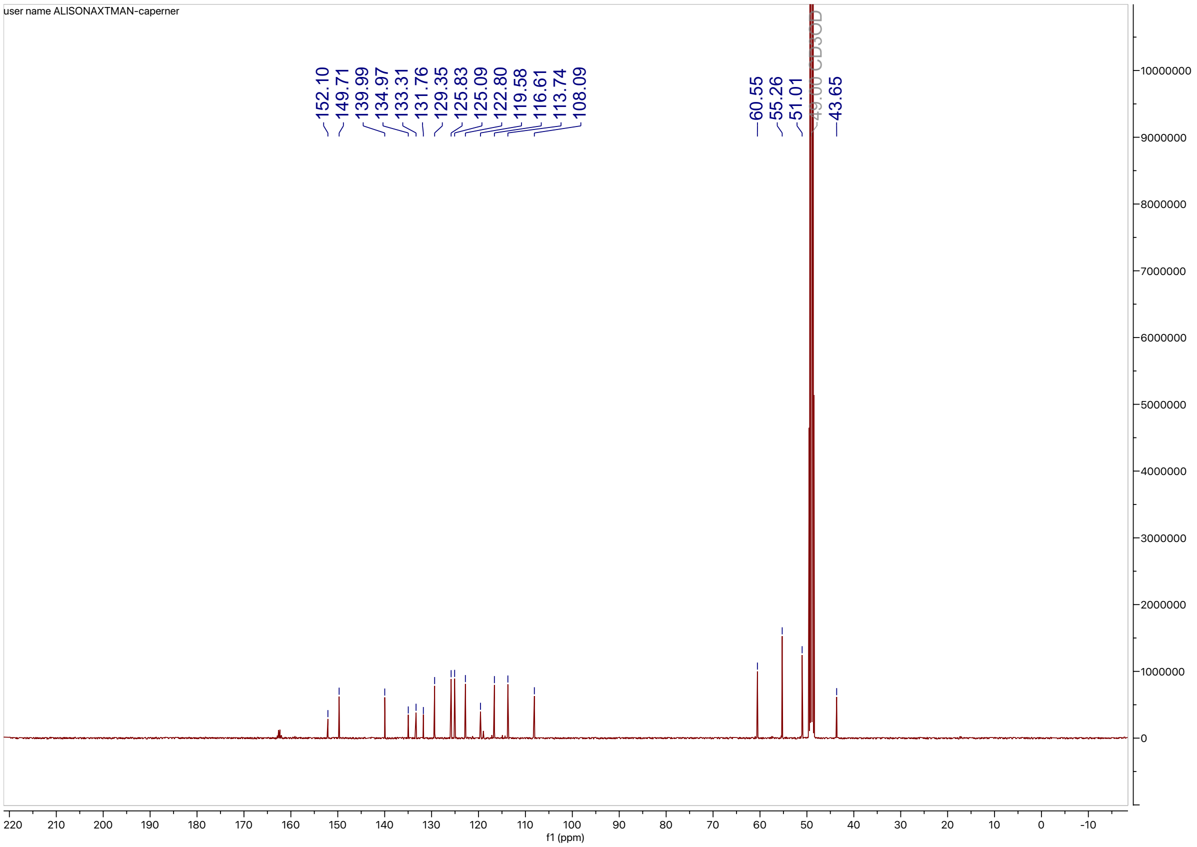

HPLC trace for compound **17**

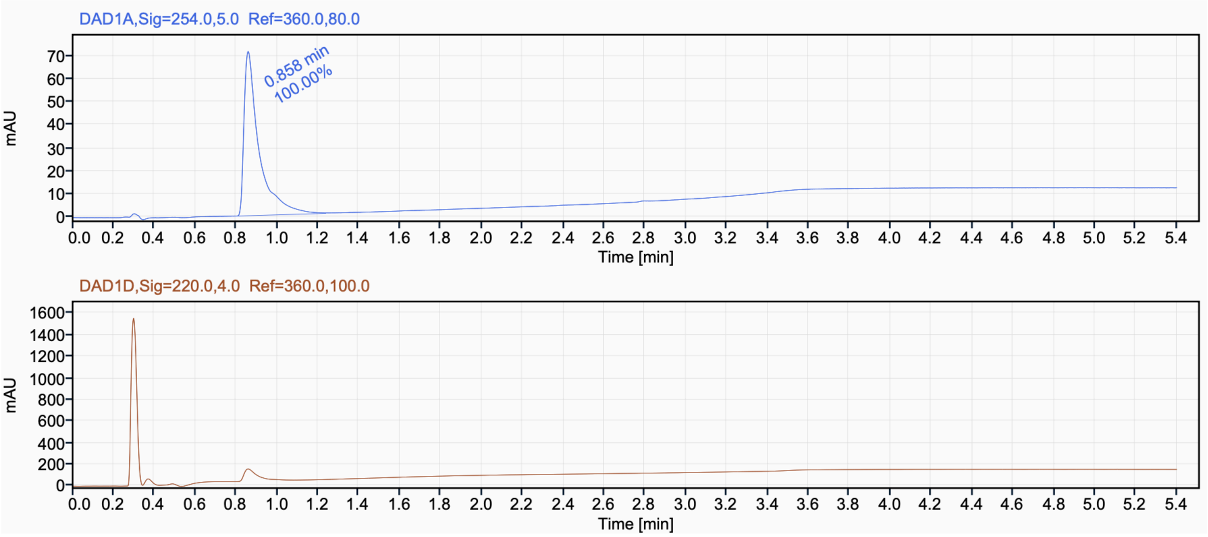

$^{1}H$ NMR (500 MHz, DMSO-$d_6$) for compound **18**

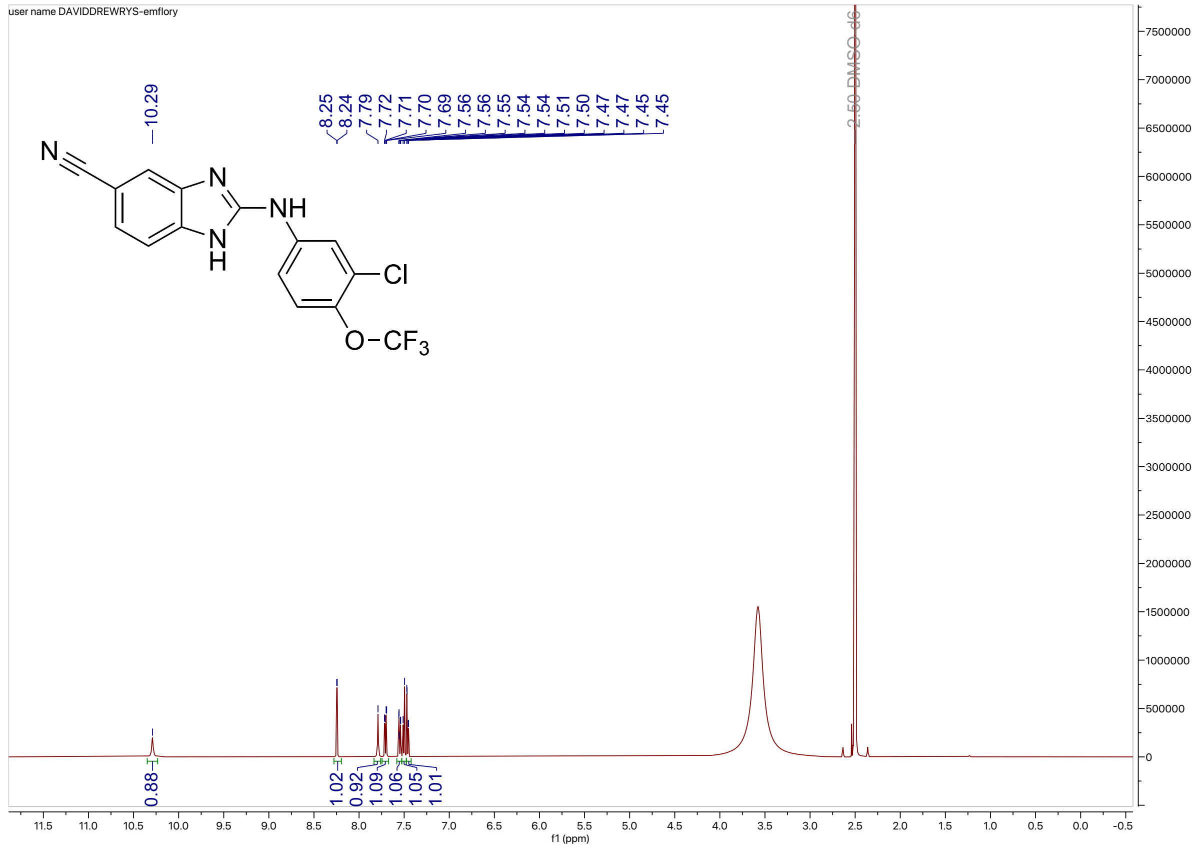

$^{13}C$ NMR (100 MHz, DMSO-$d_6$) for compound **18**

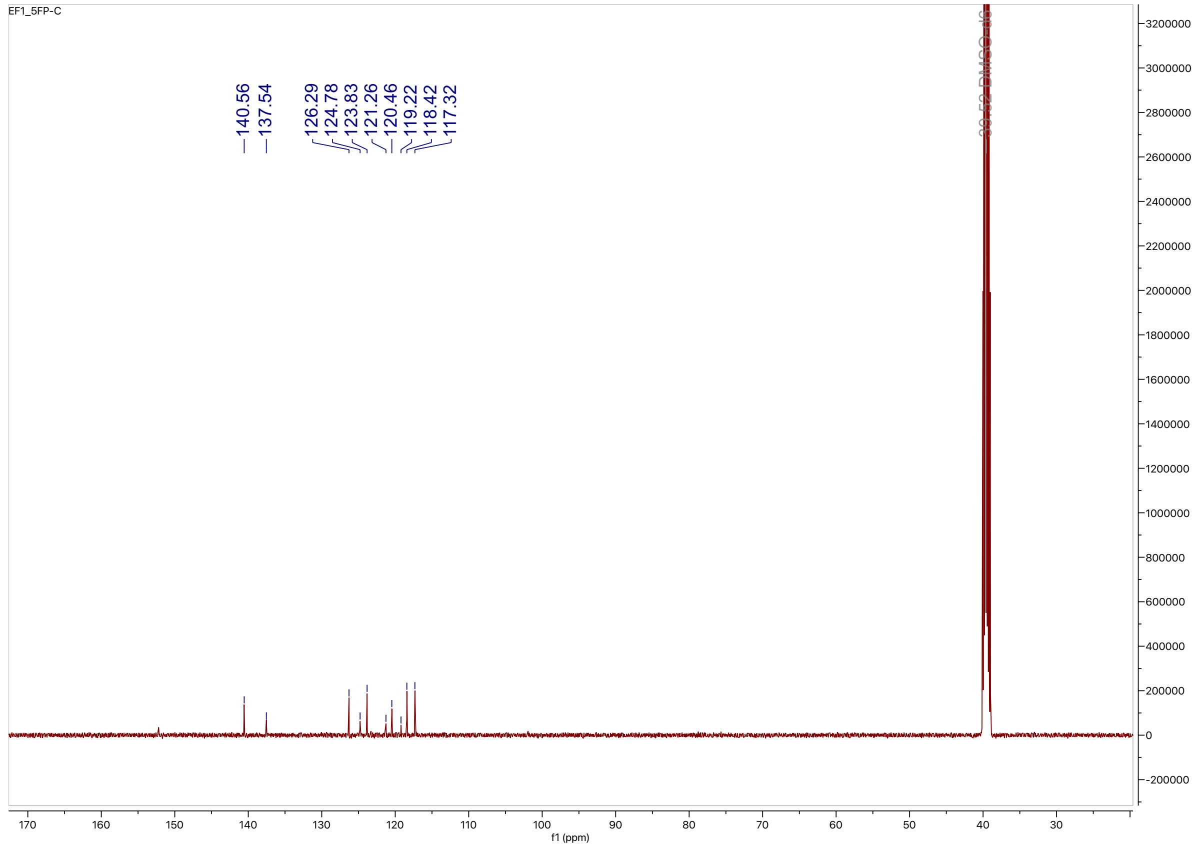

HPLC trace for compound **18**

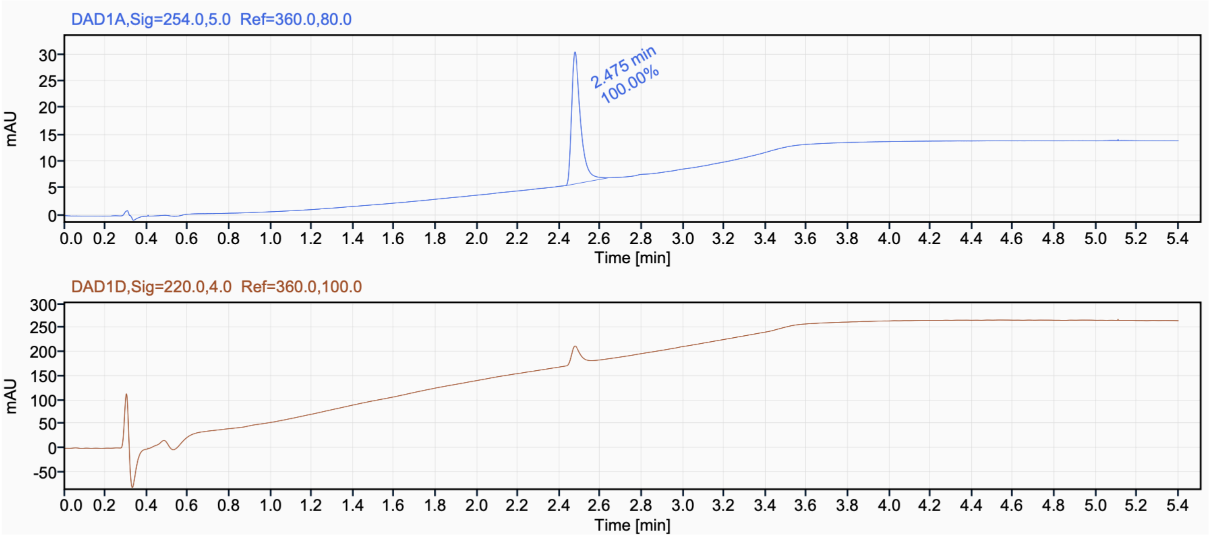

$^{1}H$ NMR (500 MHz, DMSO-$d_6$) for compound **19**

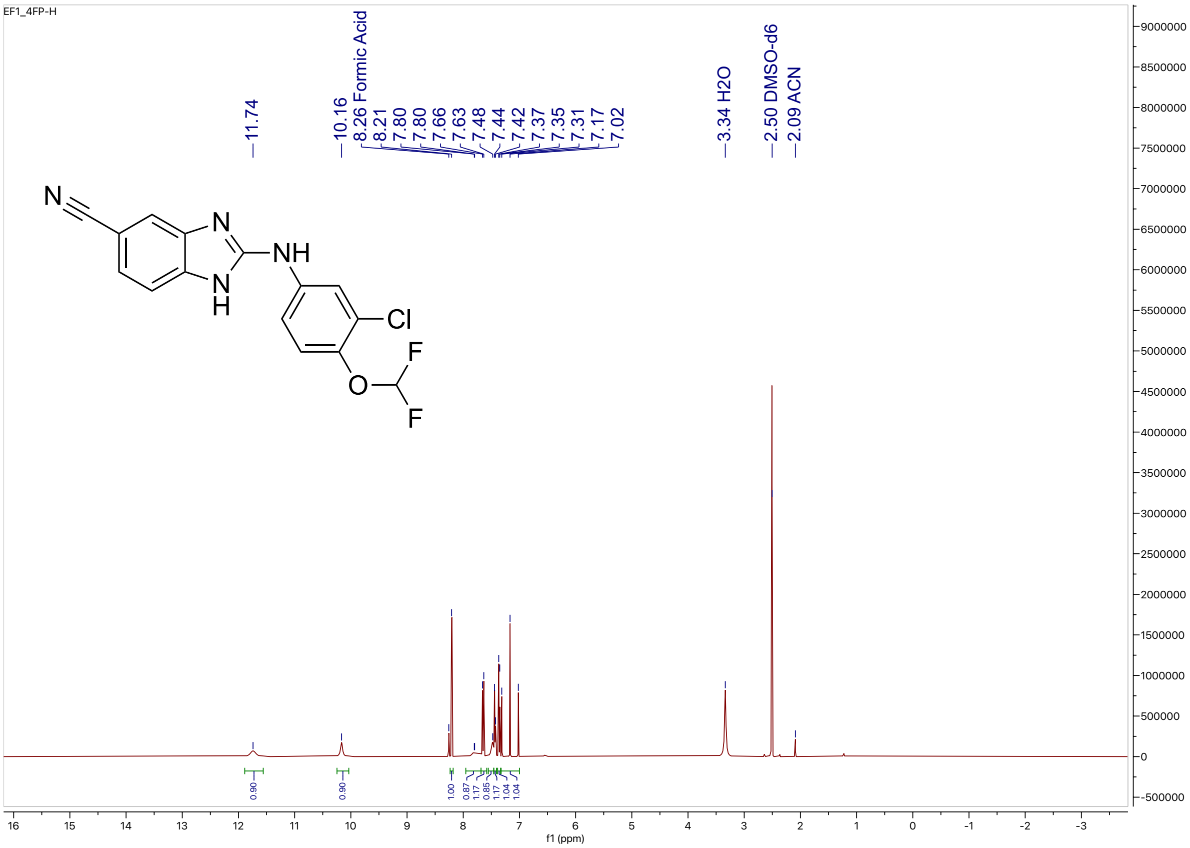


$^{13}C$ NMR (100 MHz, DMSO-$d_6$) for compound **19**

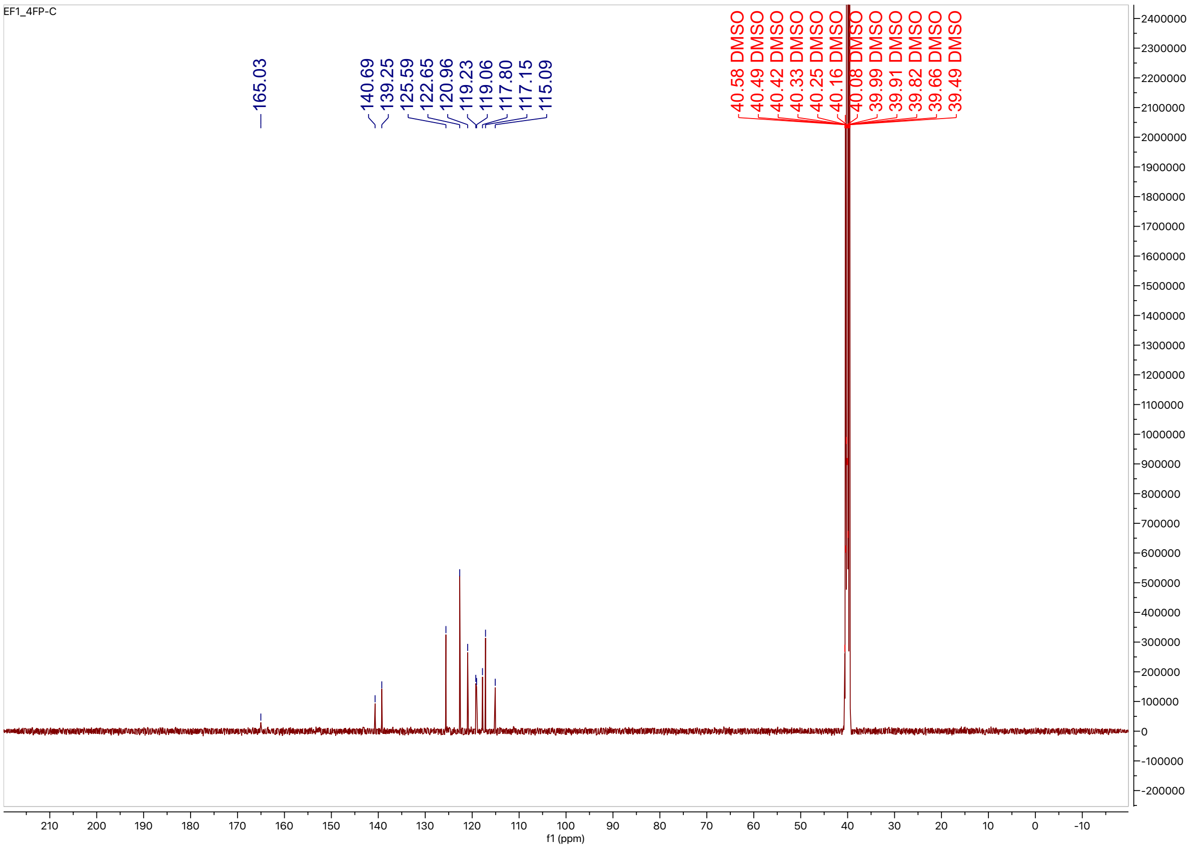

HPLC trace for compound **19**

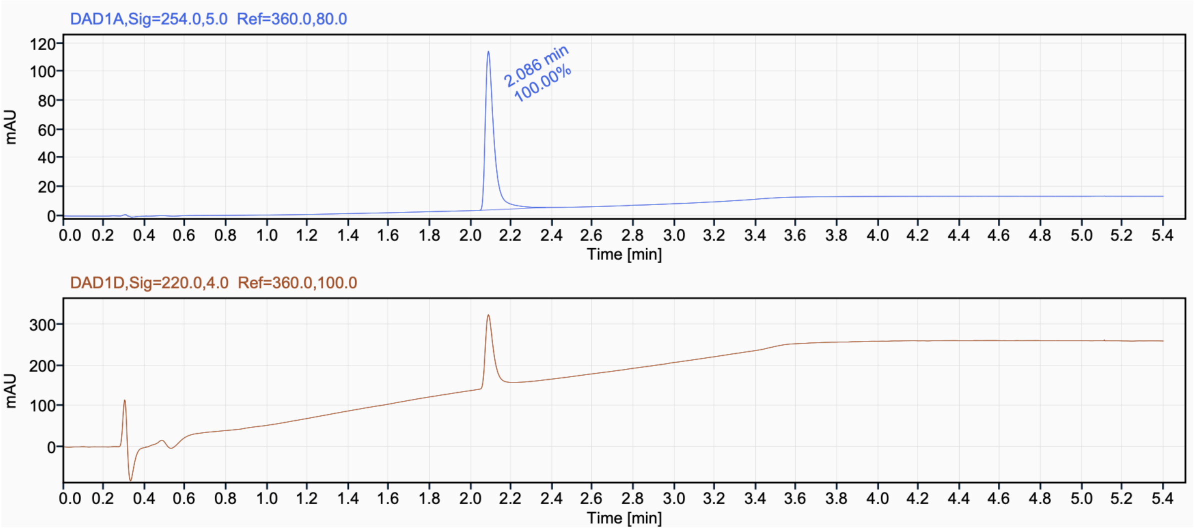

[1]H NMR (500 MHz, $CD_3OD$) for compound **20**

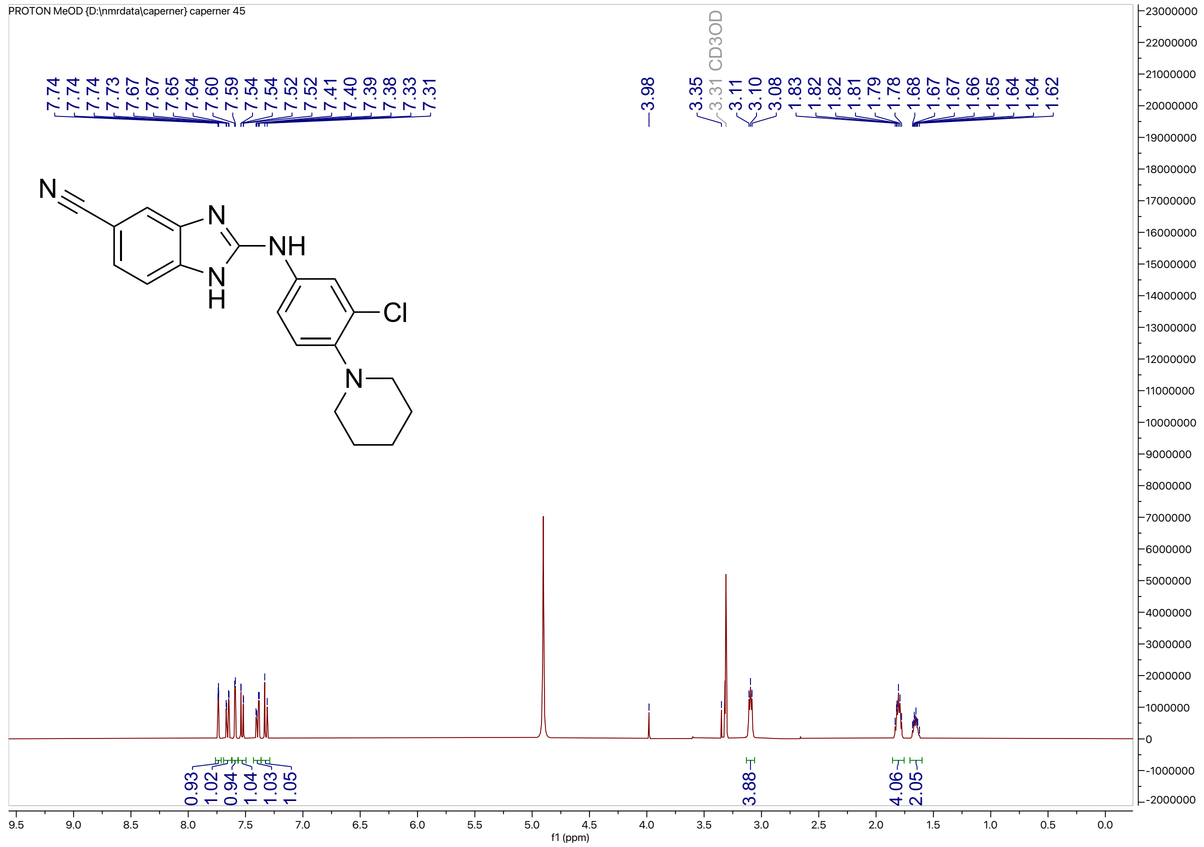

[13]C NMR (100 MHz, $CD_3OD$) for compound **20**

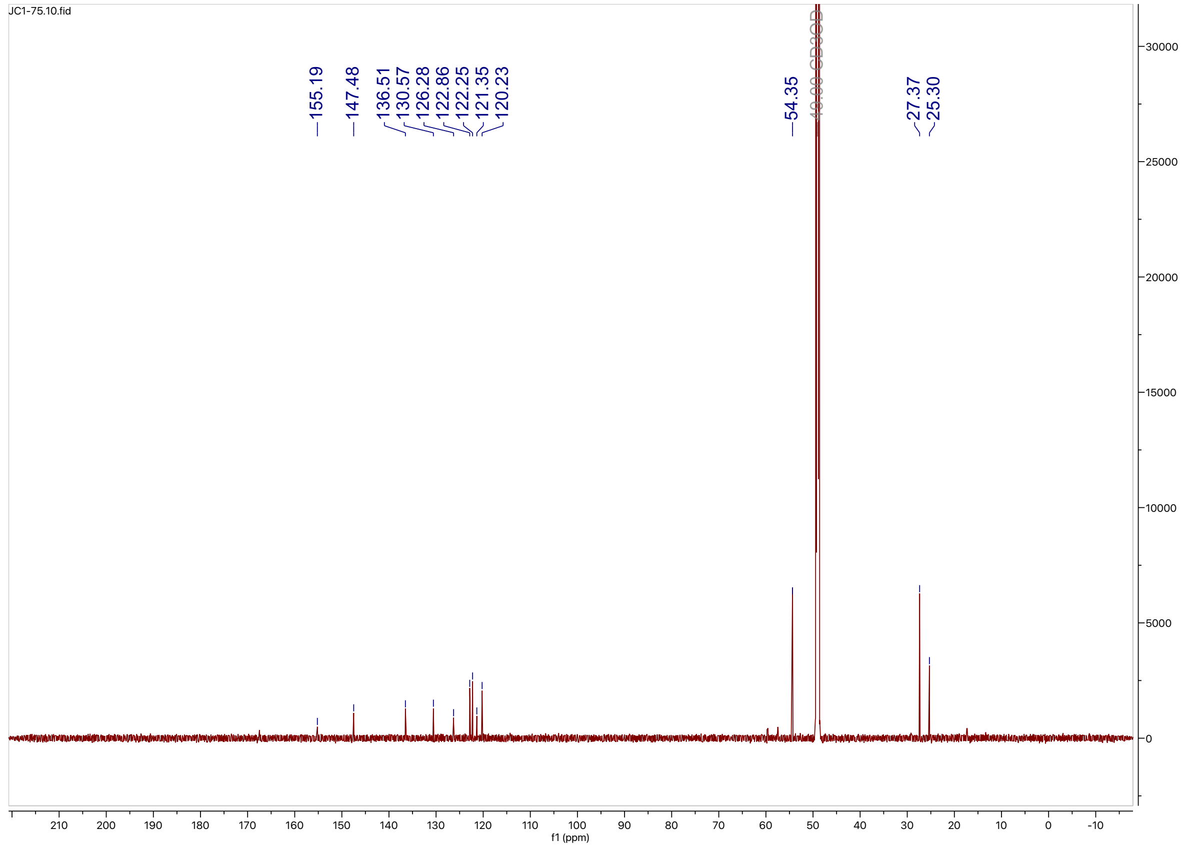

HPLC trace for compound **20**

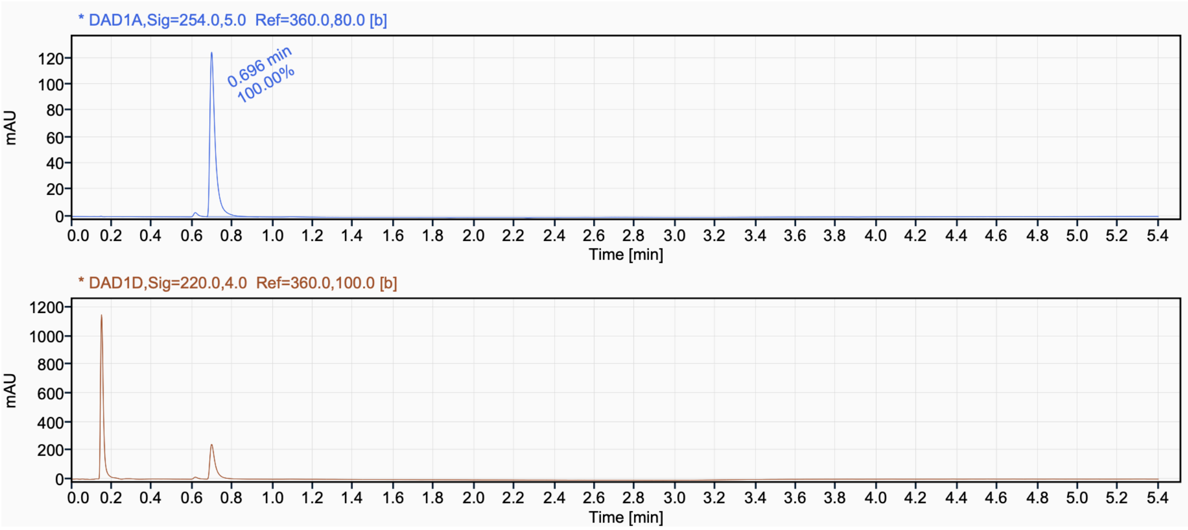

[1]H NMR (400 MHz, $CD_3OD$) for compound **21**

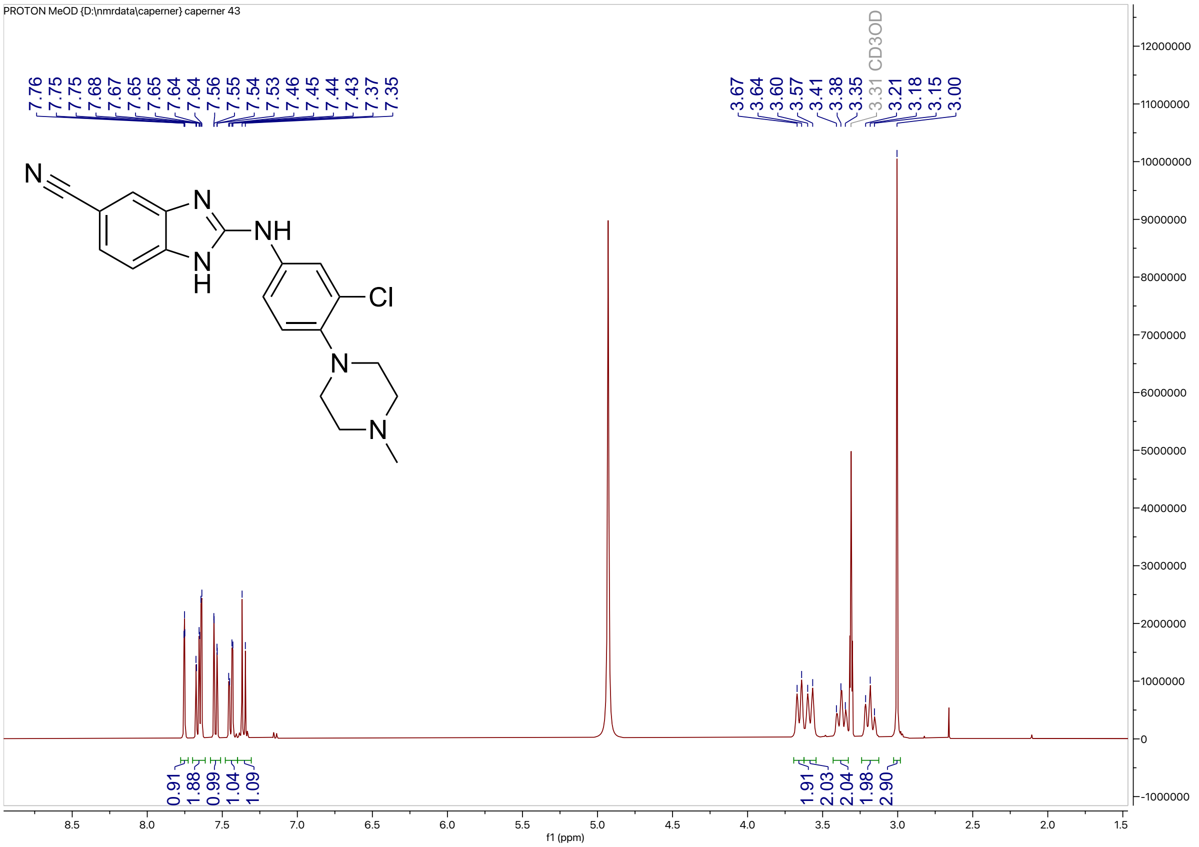


[13]C NMR (100 MHz, $CD_3OD$) for compound **21**

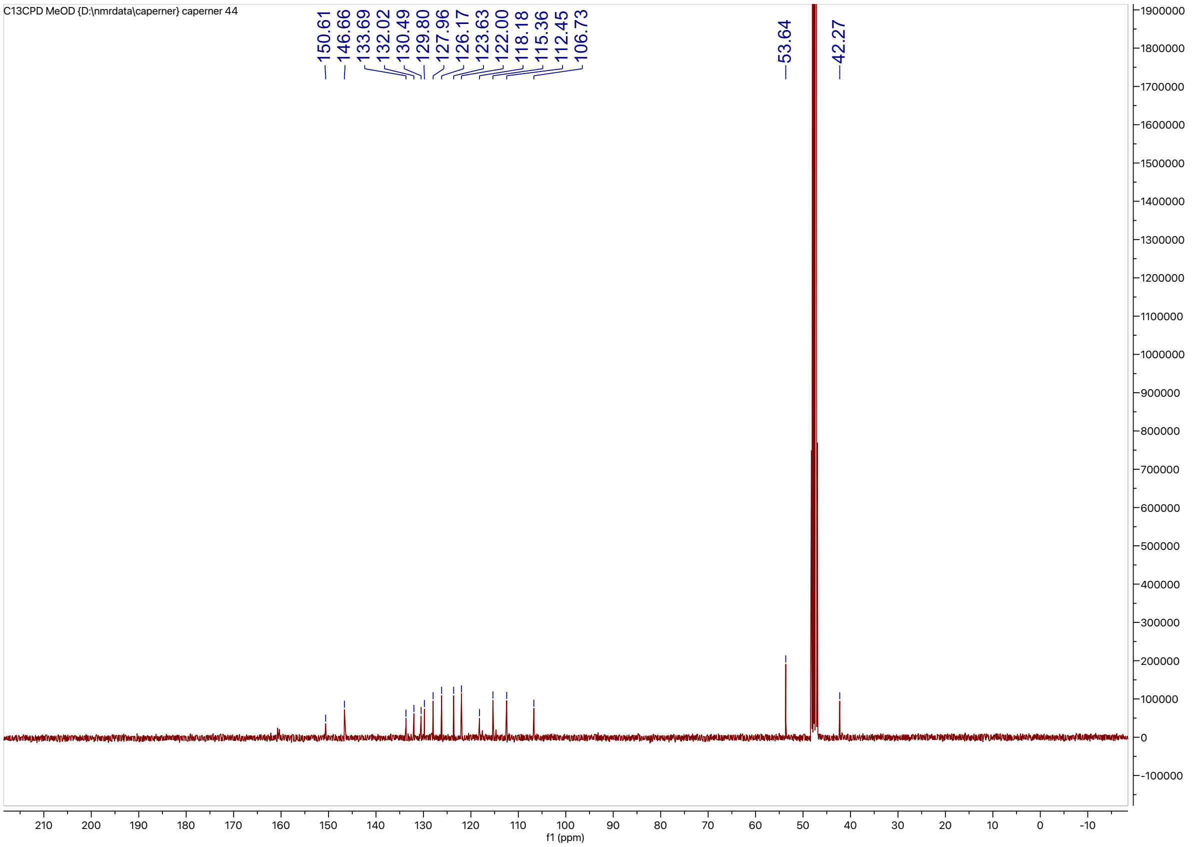

HPLC trace for compound **21**

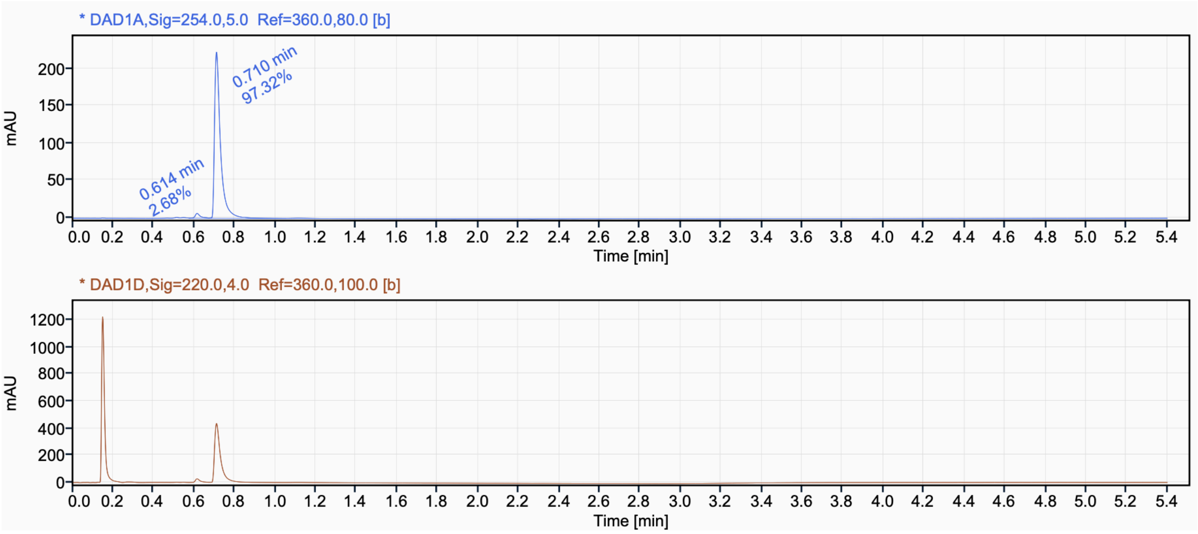

$^{1}H$ NMR (500 MHz, $CD_3OD$) for compound **22**

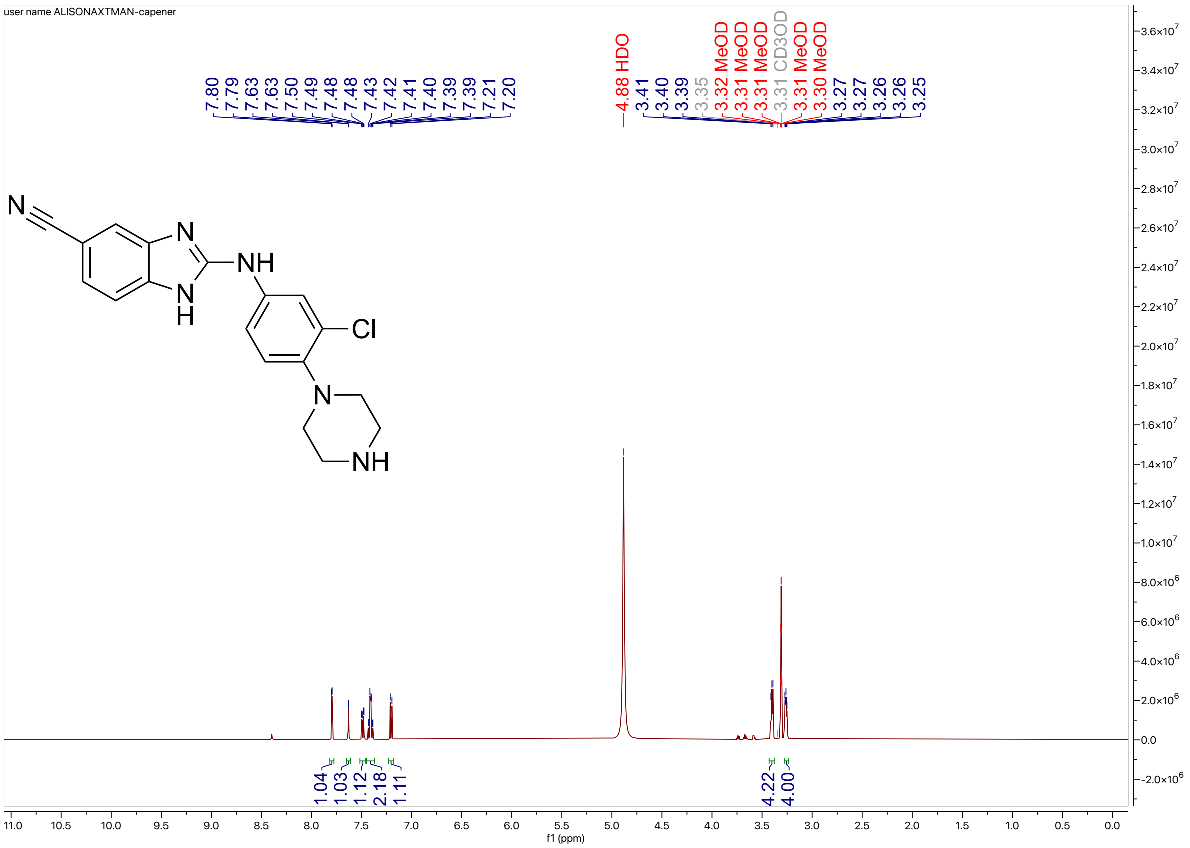


$^{13}C$ NMR (126 MHz, $CD_3OD$) for compound **22**

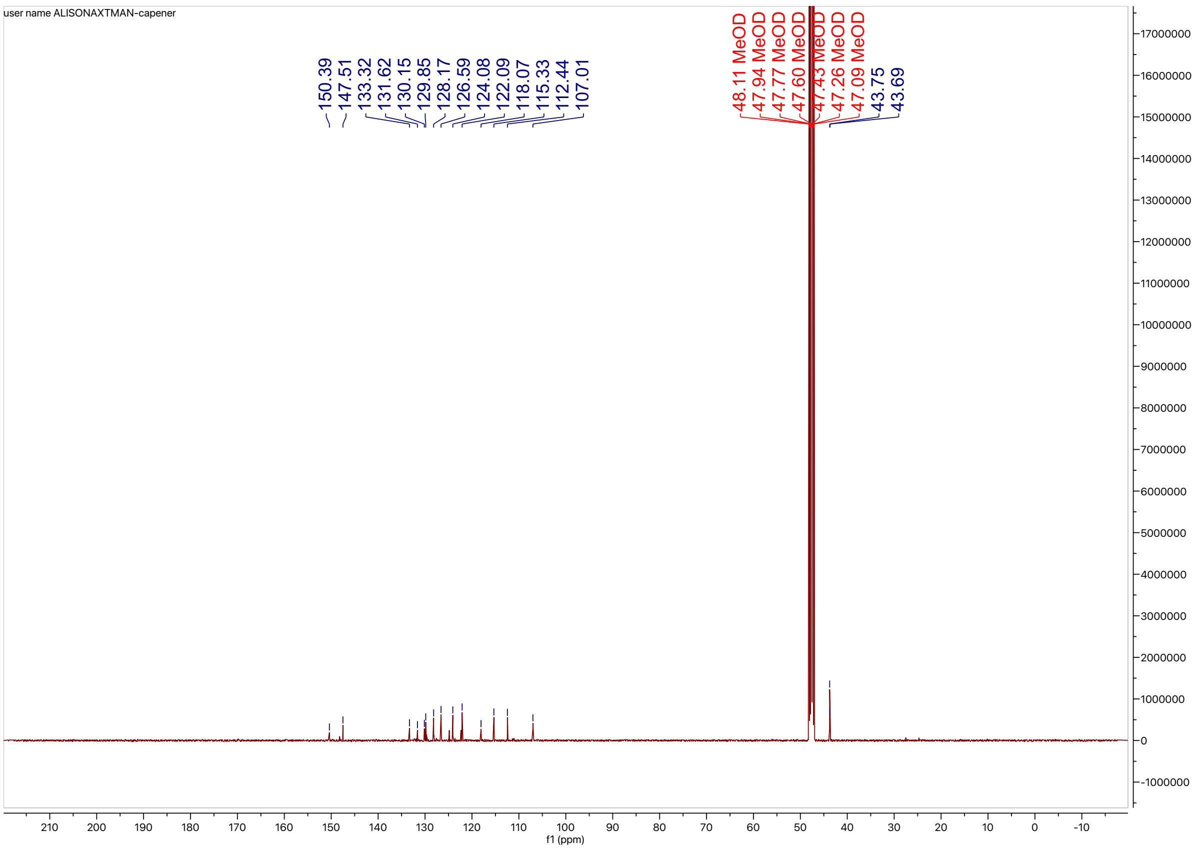

HPLC trace for compound **22**

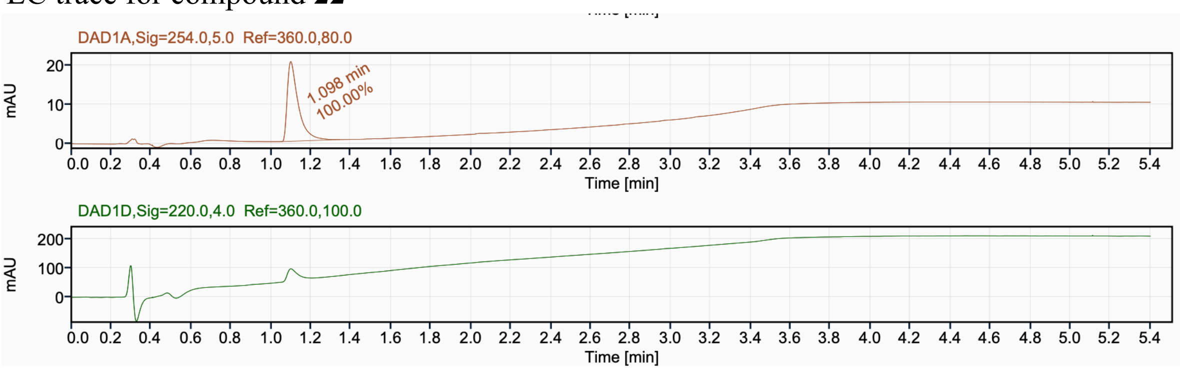

$^{1}H$ NMR (500 MHz, $CD_3OD$) for compound **23**

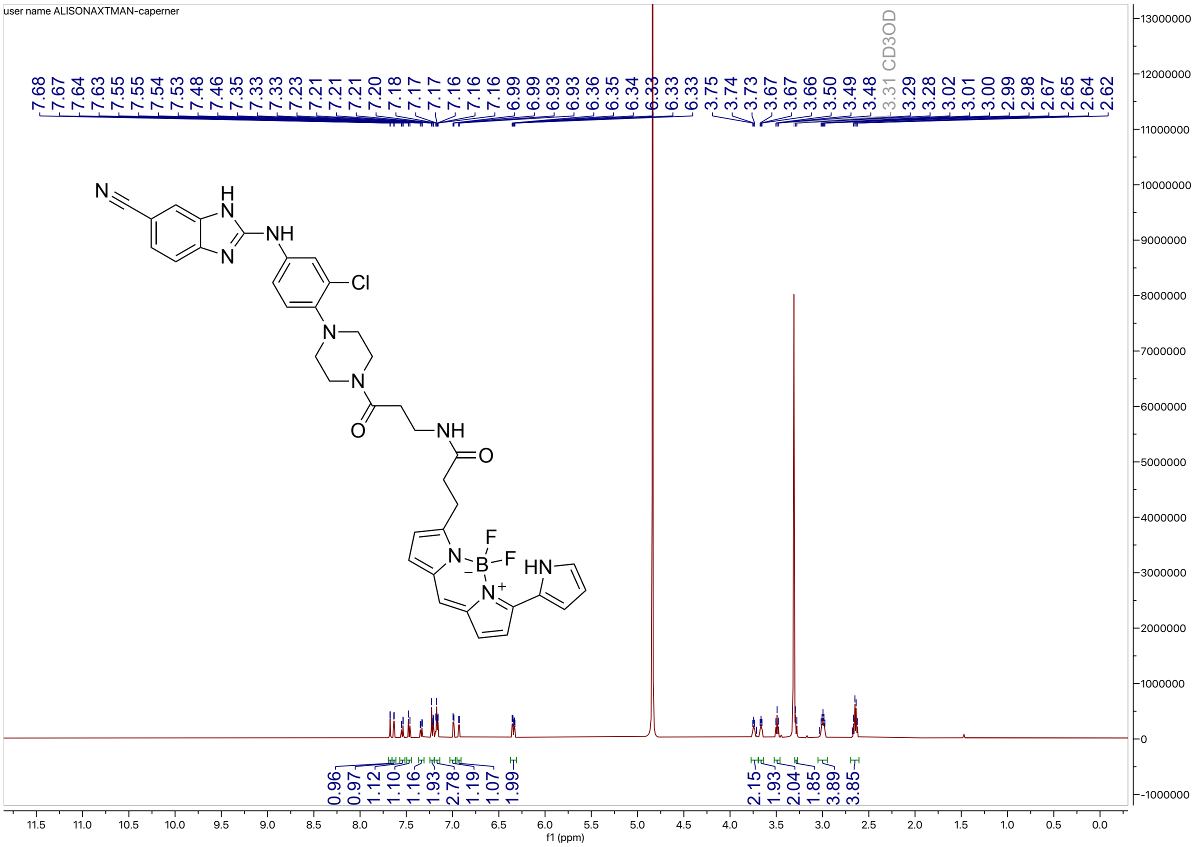


HPLC trace for compound **23**

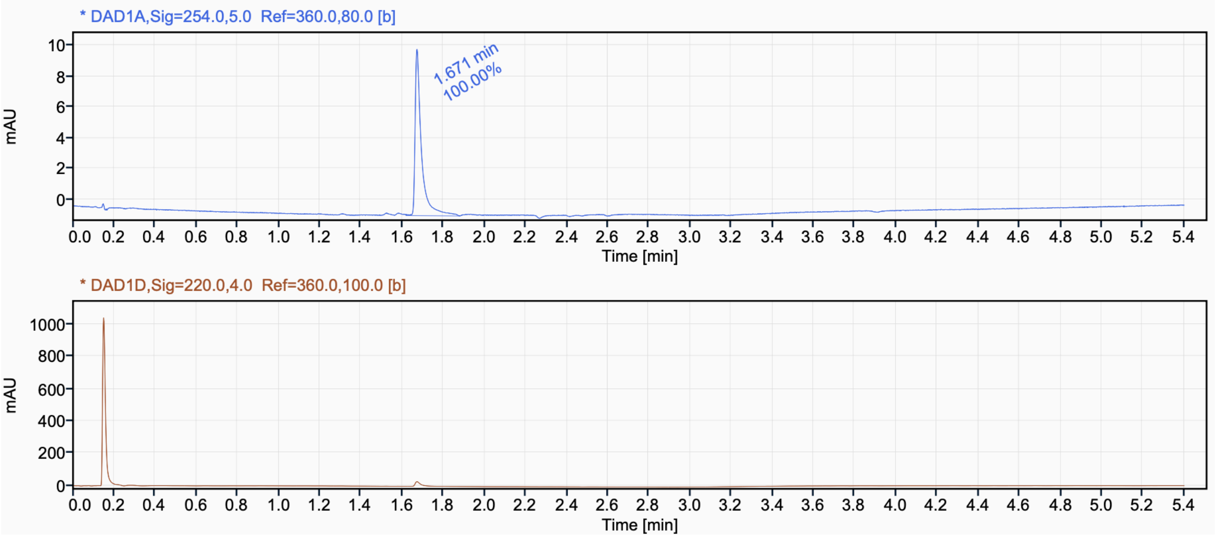

[1]H NMR (500 MHz, $CD_3OD$) for compound 1h

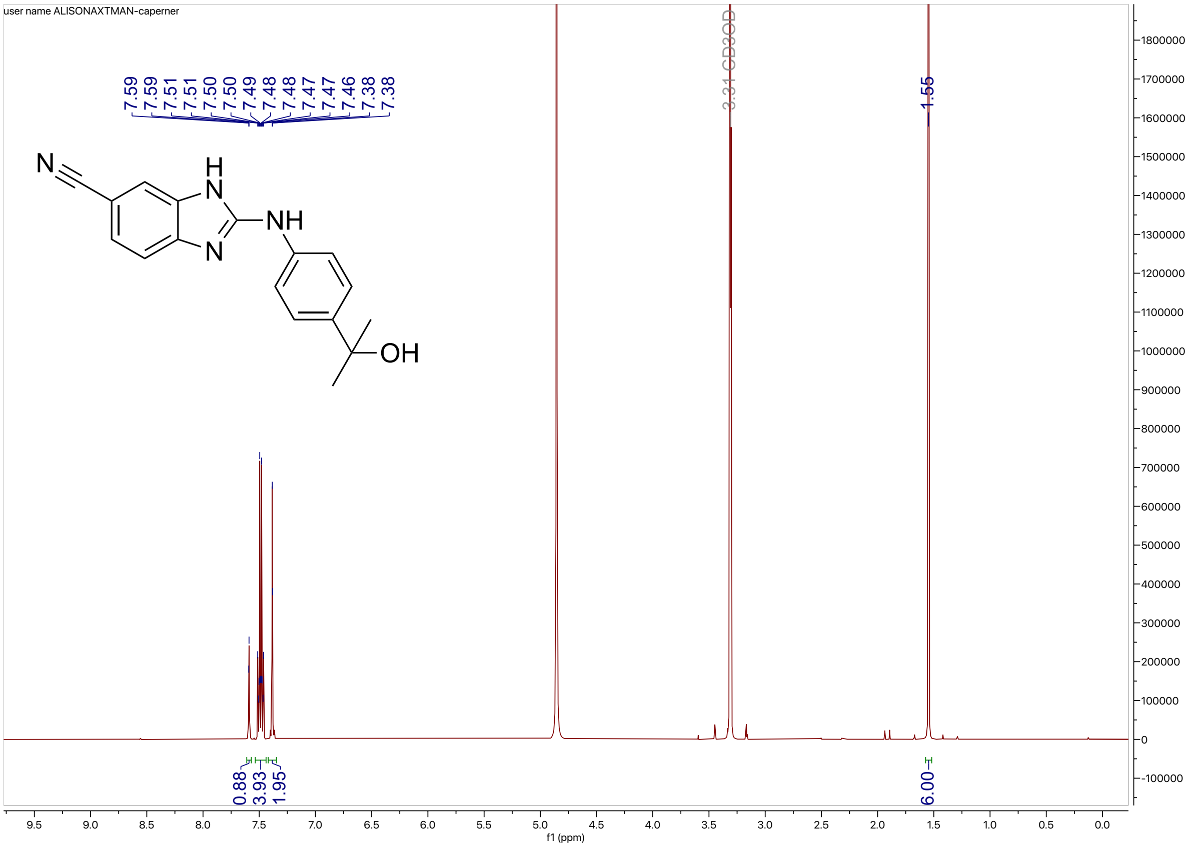


HPLC trace for compound 1h

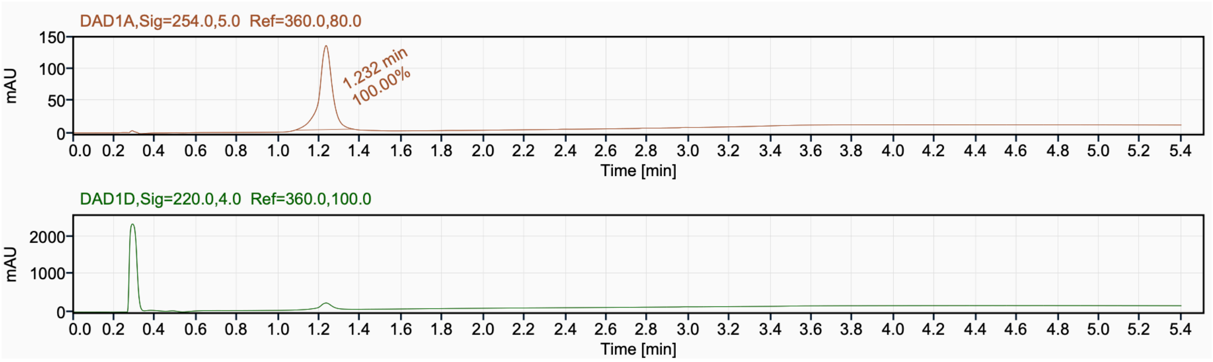
